# Shared and Distinct Genetic Risk Factors for Childhood Onset and Adult Onset Asthma: Genome- and Transcriptome-wide Studies

**DOI:** 10.1101/427427

**Authors:** Milton Pividori, Nathan Schoettler, Dan L. Nicolae, Carole Ober, Hae Kyung Im

**Author notes:** These authors contributed equally (Milton Pividori, Nathan Schoettler). Corresponding Author: HKI.

## Abstract

**Background:** Childhood and adult onset asthma differ with respect to severity and co-morbidities. Whether they also differ with respect to genetic risk factors has not been previously investigated in large samples. The goals of this study were to identify shared and distinct genetic risk loci for childhood and adult onset asthma, and the genes that may mediate the effects of associated variation.

**Methods:** We used data from UK Biobank to conduct genome-wide association studies (GWASs) in 37,846 subjects with asthma, including 9,433 childhood onset cases (onset before age 12) and 21,564 adult onset cases (onset between ages 26 and 65), and 318,237 subjects without asthma (controls; older than age 38). We conducted GWASs for childhood onset asthma and adult onset asthma each compared to shared controls, and for age of asthma onset in all 37,846 asthma cases. Enrichment studies determined the tissues in which genes at GWAS loci were most highly expressed, and PrediXcan, a transcriptome-wide gene-based test, was used to identify candidate risk genes.

**Findings:** We detected 61 independent asthma loci: 23 were childhood onset specific, one was adult onset specific, and 37 were shared. Nineteen loci were associated with age of asthma onset. Genes at the childhood onset loci were most highly expressed in skin, blood and small intestine; genes at the adult onset loci were most highly expressed in lung, blood, small intestine and spleen. PrediXcan identified 113 unique candidate genes at 22 of the 61 GWAS loci.

**Interpretation:** Genetic risk factors for adult onset asthma are largely a subset of the genetic risk for childhood onset asthma but with overall smaller effects, suggesting a greater role for non-genetic risk factors in adult onset asthma. In contrast, the onset of disease in childhood is associated with additional genes with relatively large effect sizes, and SNP-based heritability estimates that are over 3-times larger than for adult onset disease. Combined with gene expression and tissue enrichment patterns, we suggest that the establishment of disease in children is driven more by dysregulated allergy and epithelial barrier function genes whereas the etiology of adult onset asthma is more lung-centered and environmentally determined, but with immune mediated mechanisms driving disease progression in both children and adults.

**Funding:** This work was supported by the National Institutes of Health grants R01 MH107666 and P30 DK20595 to HKI, R01 HL129735, R01 HL122712, P01 HL070831, and UG3 OD023282 to CO; NS was supported by T32 HL007605.

**Research in Context:** *Evidence before this study:* Genome-wide association studies in large samples that include both childhood onset and adult onset asthma have identified many loci associated with asthma risk. However, little was known about the shared or distinct effects of those or other loci on age of asthma onset, or about the genes that may mediate the effects of loci associated with childhoon and/or adult onset asthma.

*Added value of this study:* Leveraging the resources of UK Biobank, we identified loci with both age of onset specific effects and shared effects. We further showed a significantly greater contribution of genetic variation to childhood onset asthma, implying a greater role for environmental risk factors in adult onset asthma, and different biological pathways and tissue enrichments for genes at loci associated with childhood vs adult onset asthma.

*Implications of all the available evidence:* Our results suggest that childhood onset specific loci and those associated with age of onset play a role in disease initiation, whereas the other associated loci reflect shared mechanisms of disease progression. The childhood onset specific loci highlight skin as a primiary target tissue for early onset disease and support the idea that asthma in childhood is due to impaired barrier function in the skin and other epithelial surfaces.

## Introduction

Asthma is the most prevalent chronic respiratory disease worldwide^1^. Its diagnosis is based on the presence of reversible airflow obstruction and clinical symptoms that include wheeze, cough, and shortness of breath. Despite these shared features, asthma is likely many different conditions. In particular, childhood onset asthma and adult onset asthma differ with respect to sex ratios, triggers of exacerbation, associated co-morbidities, severity^2, 3^, and potentially also for genetic risk factors^4, 5^. For example, the most often replicated and most statistically significant genome-wide association study (GWAS) single nucleotide polymorphisms (SNPs) at the 17q12-21 locus are specific to asthma with onset of symptoms in early life^6^. Only one asthma GWAS to date was performed in adult onset cases. The GABRIEL Consortium included 1,947 cases with adult onset asthma (defined as 16 years of age or older) and 3,669 adult controls^7^. Overall, genome-wide significant variants in the combined sample of 10,365 cases and 16,110 controls revealed larger odds ratios (ORs) in the childhood onset group compared to the adult onset group, but no loci reached significance in the adult onset cases, likely due to low power. It remains unknown, therefore, whether loci other than 17q12-21 contribute specifically to childhood onset or adult onset asthma. Delineating genetic risk factors affecting age of onset from those that are shared between childhood and adult onset asthma could provide insights into the molecular mechanisms contributing to different clinical manifestations of asthma that is diagnosed at different ages.

To address this question and directly compare genetic risk architectures of adult onset and childhood onset asthma, we leveraged UK Biobank (UKB), a large-scale prospective study collecting demographic, clinical, medical history, and genetic data for nearly 500,000 participants^8^. We performed three asthma GWASs, one in 9,433 adults with a diagnosis of asthma before age 12 years (childhood onset cases) and one in 21,564 adults with a diagnosis of asthma between age 26 and 65 years (adult onset cases), each compared to 318,237 adults (>38 years) who did not have a diagnosis of asthma or chronic obstructive lung disease (COPD) (controls). In our study sample, the proportion of self-reported doctor diagnosed asthma was 37,846/318,237 (11·9%), lower than the UK lifetime prevalence of “patient reported clinician diagnosed asthma” of 15·6%^9^ and consistent with the known healthy volunteer bias in UK Biobank^10^. In addition, we conducted an age of onset GWAS in 37,846 asthma cases. Our goals were to identify shared and distinct genetic risk loci for childhood and adult onset asthma, and to identify genes that may mediate the effects of associations at age of onset specific loci and those shared between childhood and adult onset asthma.

## Methods

### Sample composition and case definitions

Data for 376,358 British white individuals from UKB data release July 2017 were used^8^. We extracted disease status (asthma, allergic rhinitis, atopic dermatitis, food allergy, COPD, emphysema, chronic bronchitis), age of onset of asthma, and sex from self-reported questionnaires and hospital records (ICD10 codes) by querying our in-house protected UKB database server^11^. See our ‘reproducible research’ and other details in the appendix (pp 4-10). Among the 37,846 asthma cases included in our study, all had self-reported doctor diagnosed asthma. Of those, 15,519 also had hospital records with an asthma diagnosis (ICD10 codes).

We defined childhood and adult onset asthma using strict age of onset criteria that would minimize the likelihood of misclassification, and considered asthma cases with onset < 12 years of age as childhood onset cases (n=9,433; 3,462/9,433 [36·7%] with ICD10 codes) and with onset > 25 and before 66 years of age as adult onset cases (n=21,564; 9,260/21,564 [42·9%] with ICD10 codes). UKB participants without an asthma diagnosis were included as controls (n=318,237). Individuals with COPD, emphysema or chronic bronchitis were excluded from adult onset cases and controls. For the age of onset GWAS, we included an additional 6,849 subjects with asthma with onset between the ages of 12 and 25 years of age (2,797/6,849 [40·8%] with ICD10 codes). Characteristics of the sample are shown in Table 1. After genotype quality control, 10,894,596 variants were available for analyses. See appendix (pp 4-7) for additional details on genotype QC and phenotype definitions.

**Table 1.** Characteristics of the asthma cases and controls. NA, not applicable.

|  | **Childhood**<br>**Onset**<br>**(N=9,433)** | **Adolescent or**<br>**Young Adult**<br>**Onset (N=6,849)** | **Adulthood Onset**<br>**(N=21,564)** | **Controls**<br>**(N=318,237)** |
| --- | --- | --- | --- | --- |
| **Age at recruitment (yr)** |  |  |  |  |
| Mean (±SD) | 55±8 | 52±8 | 57±8 | 57±8 |
| Range | 40 - 70 | 40 - 70 | 40 - 70 | 39 - 73 |
| **Age at asthma onset (yr)<sup>a</sup>** |  |  |  |  |
| Mean (±SD) | 6±3 | 19±4 | 44±10 | NA |
| Range | 0 - 11 | 12 - 25 | 26 - 65 | NA |
| **Sex (% female)** | 40.7 | 57.4 | 63.6 | 53.5 |
| **Asthma medication use in the**<br>**past 12 months, N (%)** | 1,383 (14.7) | 1,124 (16.4) | 4,081 (18.9) | NA |
| **Current smokers N (%)** | 802 (8.5) | 688 (10.0) | 1,379 (6.4) | 30,056 (9.4) |
| **Allergic diseases ever** |  |  |  |  |
| Allergic rhinitis, N (%) | 2,537 (26.9) | 2,042 (29.8) | 4,660 (21.6) | 27,289 (8.6) |
| Atopic dermatitis, N (%) | 1,126 (11.9) | 473 (6.9) | 927 (4.3) | 7,168 (2.3) |
| Food allergy, N (%) | 138 (1.5) | 63 (0.9) | 172 (0.8) | 1,229 (0.4) |
| Any allergic disease, N (%) | 3,205 (34.0) | 2,321 (33.9) | 5,306 (24.6) | 33,808 (10.6) |

### Genome-wide association studies

We conducted a childhood onset and adult onset GWAS by logistic regression and an age of onset GWAS by linear regression, both using the allele dosages under an additive genetic model, as implemented in Hail (https://github.com/hail-is/hail). All analyses were performed using the Bionimbus Protected Data Cloud^12^. In all three GWASs, we included sex and the first 10 genetic principal components as covariates and used a genome-wide significance threshold of 5×10^−8^. FUMA^13^, an integrative post-GWAS annotation web-based tool, was used to define independent risk loci and identify enriched tissues. FUMA defines independent loci using linkage disequilibrium (LD) information from the 1000 Genomes project^14^. We specified an *r^2^* threshold >0·6 between genome-wide significant SNPs to represent a single locus. Tissue enrichments were calculated in FUMA using a hypergeometric test to determine overrepresentation of genes mapped to risk loci (by physical distance) among highly expressed genes in each tissue in GTEx relative to all others.

Childhood onset and adult onset specific loci were defined as those that were genome-wide significant in either the childhood onset or adult onset GWAS but were not associated with asthma at p<0·05 in the other group, and the 95% confidence intervals (CIs) of the respective ORs did not overlap. All other GWAS loci that were genome-wide signficant in at least one of the two GWASs were considered to be shared.

### SNP-based heritability estimation

We used Linkage Disequilibrium Score regression (LDSC) to estimate the SNP-based heritability from the childhood onset, adult onset and age of onset GWAS summary statistics, using SNPs overlapping with HapMap3 variants, as recommended^15^, and estimated population prevalence of 8·68% for childhood onset asthma and 9·55% for adult onset asthma^16^. Heritability estimates for binary traits are reported in the liability scale.

### Sensitivity analysis to adult onset misclassification

We performed two sets of sensitivity analyses to assess 1) the effects of potential poor recall of age of onset among subjects with adult onset asthma, and 2) the effects of misclassification of COPD as asthma among the adult onset cases, even with exclusion of cases with a reported diagnosis of COPD, emphysma or chronic bronchitis. First, to assure that the adult onset cases did not include a significant proportion of childhood onset asthma in which symptoms remitted in early life but then relapsed in adulthood, we replaced adult onset cases with increasing proportions of randomly selected childhood onset cases, and then tested for association at the two most significant childhood onset specific loci. This procedure was repeated 20 times for each proportion to quantify the sampling variability (appendix pp 7-8). Second, we performed two analyses in which we removed either subjects with ages of asthma onset between 46 to 65 years or adult-onset cases and controls with FEV1/FVC <0·70. For each, we compared p-values and ORs to the GWAS including all adult onset cases (appendix pp 8-9).

### Predicted transcriptome association test

We used the PrediXcan^17^ framework to identify genes that may mediate associations between genetic variants and asthma risk. PrediXcan is a software tool that estimates tissue-specific gene expression profiles from an individual’s SNP genotype profile using prediction models trained in large references databases of genotypes and tissue-specific gene expression profiles. Using these genotype-imputed expression profiles, PrediXcan can perform gene-based association tests that correlate predicted expression levels with phenotypes (e.g., asthma) to identify candidate causal genes from GWAS data. We used a summary version of PrediXcan, which has high concordance with the individual level version (R^2^ >0·99)^18^. For predictions, we downloaded elastic net models trained with reference transcriptome data from the GTEx consortium^19^ (http://predictdb.org) for 49 tissues (appendix Table 1).

PrediXcan was run separately in the childhood onset and adult onset cases, each with the same controls. Significance was determined using a Bonferroni correction for the 38,608 genes (p<1·29×10^−6^) that were expressed in the five tissues determined by FUMA to be enriched (skin, lung tissue, whole blood, small intestine and spleen; see Results). We defined childhood onset specific genes as those whose predicted expression is significantly associated with childhood onset asthma and the variants that predict their expression are within childhood onset specific loci. Adult onset specific genes were similarly defined. Because SNPs at shared loci may predict the expression of genes that are associated only in childhood onset or adult onset cases, we also considered genes to be age of onset specific if they were significantly associated with asthma at p<1·29×10^−6^ in one age group and not associated with asthma at p<0·05 in the other. All other genes were considered shared.

## Results

### Genome-wide association studies of asthma

We first conducted GWASs of childhood onset and adult onset asthma. These studies revealed 61 independent loci associated with asthma, 52 were significant in the childhood onset asthma GWAS and 19 were significant in the adult onset asthma GWAS (p<5×10^−8^) (Figure 1). The GWAS results were robust to inclusion of varying numbers of PCs (10, 14, or 20) (appendix Figure 1) and to limiting the sample to cases with diagnoses based on ICD10 codes (appendix Figure 2). Twenty-eight of the 61 loci, in 27 chromosomal regions, were not previously reported in the GWAS catalog^20^, including one study also conducted in UKB subjects but focused on the phenotype asthma+allergies^21^. Among the 28 new loci, 17 were significant in the childhood onset GWAS, one was significant in the adult onset GWAS, and 10 were significant in both (Table 2). Some of these loci contained genes that have been associated with asthma in candidate gene studies (e.g., *FADS2*^22^, *MUC5AC*^23, 24^ and *TBX2*1^25^), which provide independent validation of our GWAS findings. The lead SNP or an LD surrogate SNP (*r*^2^>0·40) was reported for 23 of the 28 new loci in the TAGC GWAS^26^. SNPs at 16 of the 23 loci were associated with asthma in TAGC (p<0·05) and had the same direction of effects as in the UKB GWASs (appendix Table 2). To test for potential effects of asthma diagnostic criteria on the GWAS results, we conducted a GWAS comparing individuals with asthma based on self-reported doctor diagnosis alone compared to those with self-reported doctor diagnosis plus ICD10 codes. There were no differences detected between these two groups (appendix Figure 3), indicating that differences between these groups are not influencing the GWAS results discussed above.

**Figure 1.**
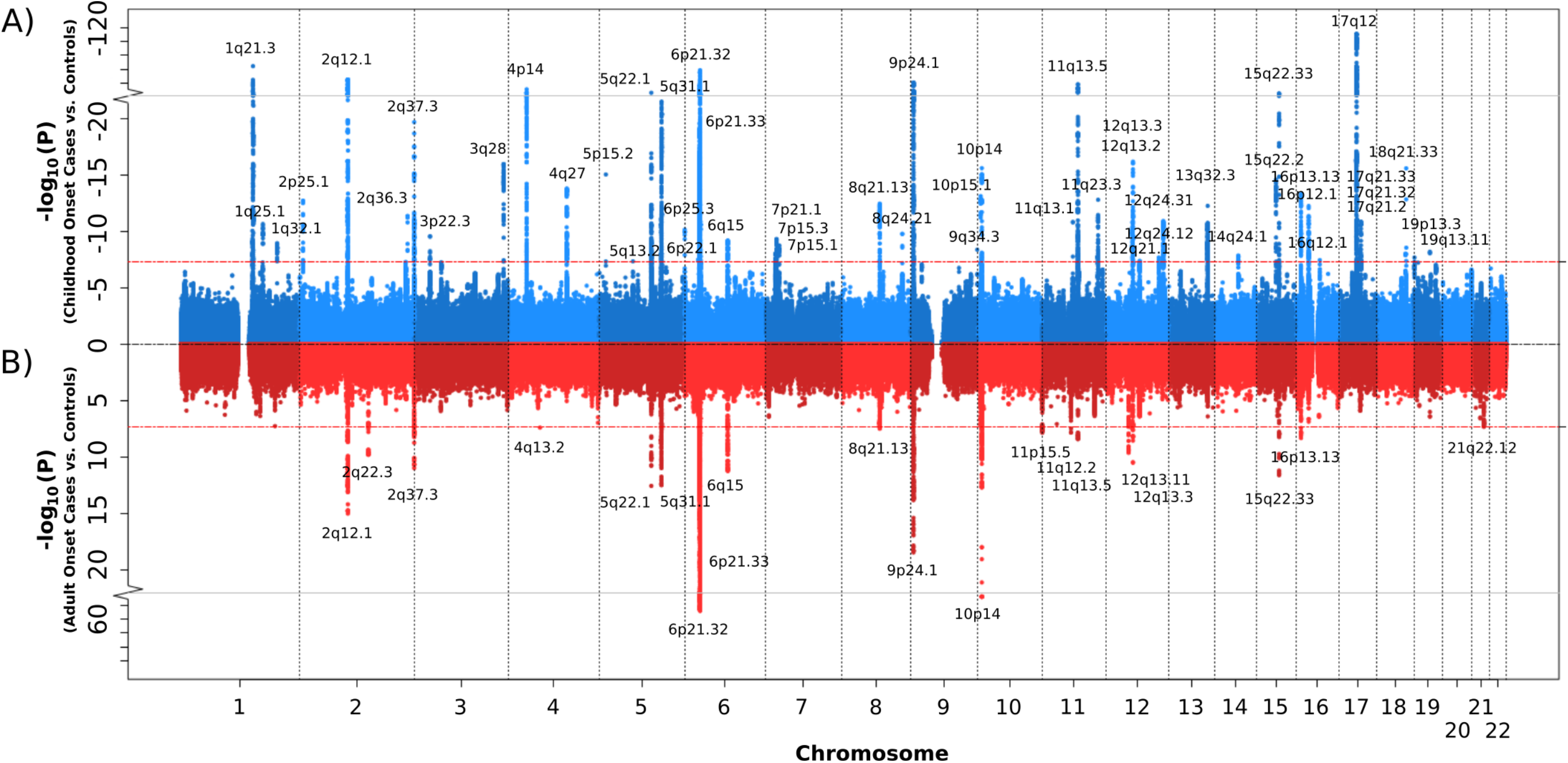
GWAS of childhood onset and adult onset asthma. Miami plot showing results for the childhood onset versus controls GWAS (blue, panel A) and adult onset versus controls GWAS (red, panel B). Each point corresponds to a SNP; the y-axes show the −log_10_p-values from the childhood onset GWAS (panel A) and adult onset GWAS (panel B). The x-axis shows the position of each SNP along the 22 autosomes. See appendix Figure 6 for the age of asthma onset Manhatten plot.

**Table 2.**
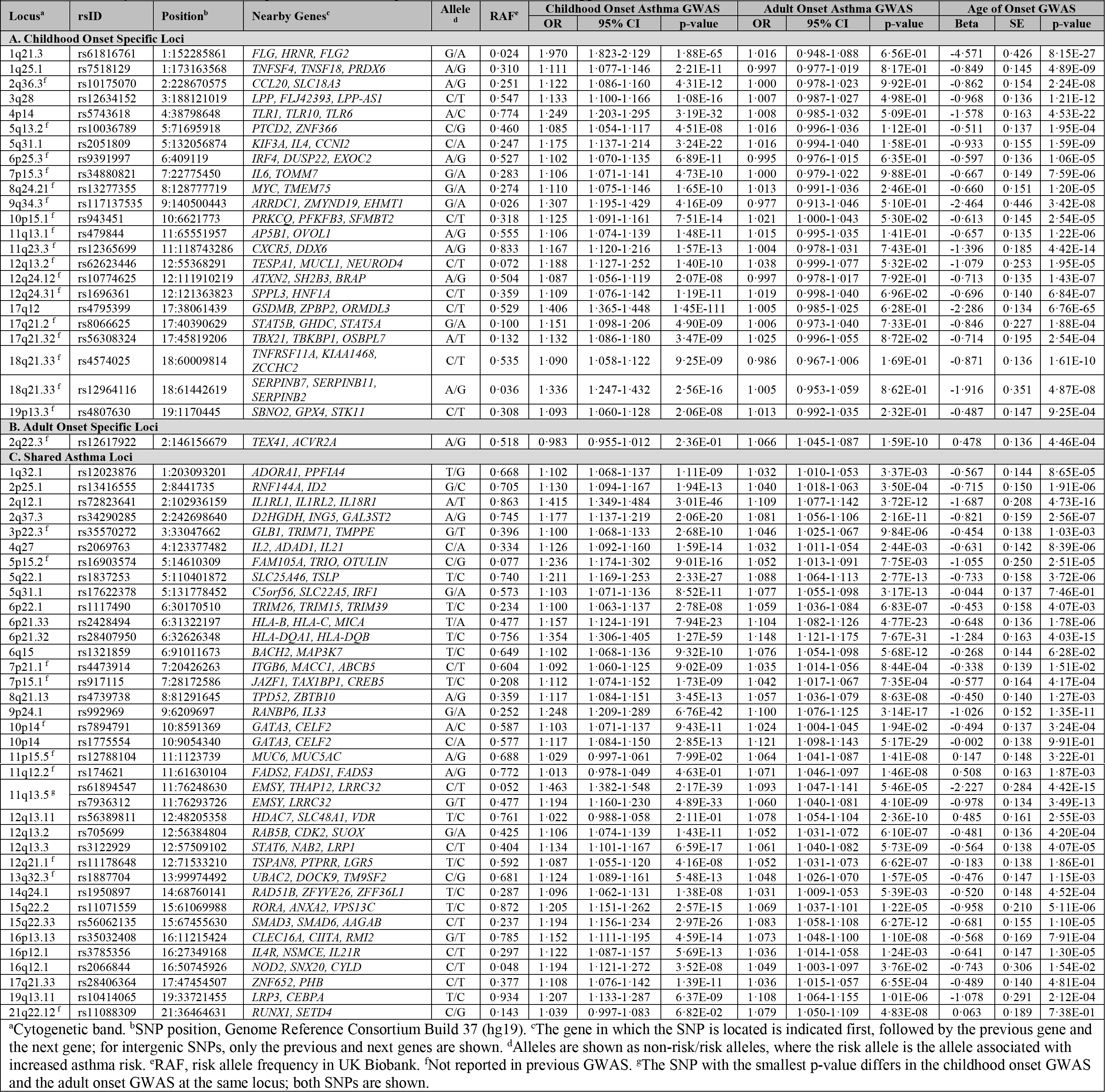
Regions with SNPs that were genome-wide significant in the childhood onset or adult onset GWAS. Information for the SNP with smallest p-value at each locus is shown. Summary statistics for the 19 age of asthma onset significant SNPs are shown in appendix Table 5.

As in previous GWASs comprised largely of children, the most significant locus in the childhood onset GWAS is at 17q12^27^, with the lead SNP in the *GSDMB* gene (Figure 1A). However, the estimated ORs for the lead SNPs at three other loci were similar to or larger than the lead SNP at the 17q locus. The lead SNPs in *IL1RL1* at 2q12.1 and in *EMSY* at 11q13.5 had effect sizes on childhood onset asthma similar to the lead SNP in *GSDMB* on 17q (Table 2). Both loci have been prominent in previous asthma GWAS. The lead SNP at the 1q21.3 locus, corresponding to a nonsense mutation (R501X) in the filaggrin (*FLG*) gene at 1q21.3, had the largest OR overall. Variants in *FLG* have been robustly associated with food allergies^28–30^ and atopic dermatitis^31–33^, but previous associations with asthma have been in the context of other allergic conditions^21, 34^. Whether variants in *FLG* are associated with risk for childhood onset asthma independent of its effects on early life allergic disease is yet unknown.

To address this possibility, we repeated the childhood onset GWAS after excluding 3,205 childhood onset cases and 5,785 controls who reported having a history of allergic rhinitis, AD or food allergy. As expected in a smaller sample the p-values were overall larger, but the ORs were strikingly similar (appendix Figure 4 and Table 3). Even though the OR at the *FLG* locus on 1q21.3 decreased from 1·97 (95% CI 1·82, 2.13) to 1·61 (95% CI 1·49, 1·74), it remained both highly significant (p=2·45×10^−19^) and the largest OR for childhood onset. These results suggest both a critical role for the allergic diathesis in the development of asthma in childhood and a shared architecture between allergic disease and childhood onset asthma, as previously discussed^34, 35^.

The most significant association in the adult onset GWAS was in the HLA region, with independent associations at the HLA-C/B (6p21·33) and HLA-DR/DQ (6p21·32) loci (Figure 1B). Compared to the childhood onset GWAS, effect sizes were quite small in adult onset cases, with ORs reaching 1·1 at only five loci (2q12.1, 6p21.33, 6p21.32, 9p24.1, 10p14) (Table 2).

Among the 61 asthma loci, 23 were specific to childhood onset asthma and one was specific to adult onset asthma (Table 2A-B). Regional association plots for the 23 loci with childhood or adult onset specific effects are shown in appendix Figure 5. Among the remaining 38 shared loci (Table 2C), mean ORs were larger in the childhood onset cases at all but six loci (permutation test p<10^−4^; appendix pp 8-9), indicating that both more loci contribute to childhood onset asthma and even among shared loci, effect sizes are larger in childhood onset asthma cases (Figure 2). Colocalization analyses using GWAS-PW^36^ yielded results that supported our classification of age of onset specific and shared loci (appendix Table 4).

**Figure 2.**
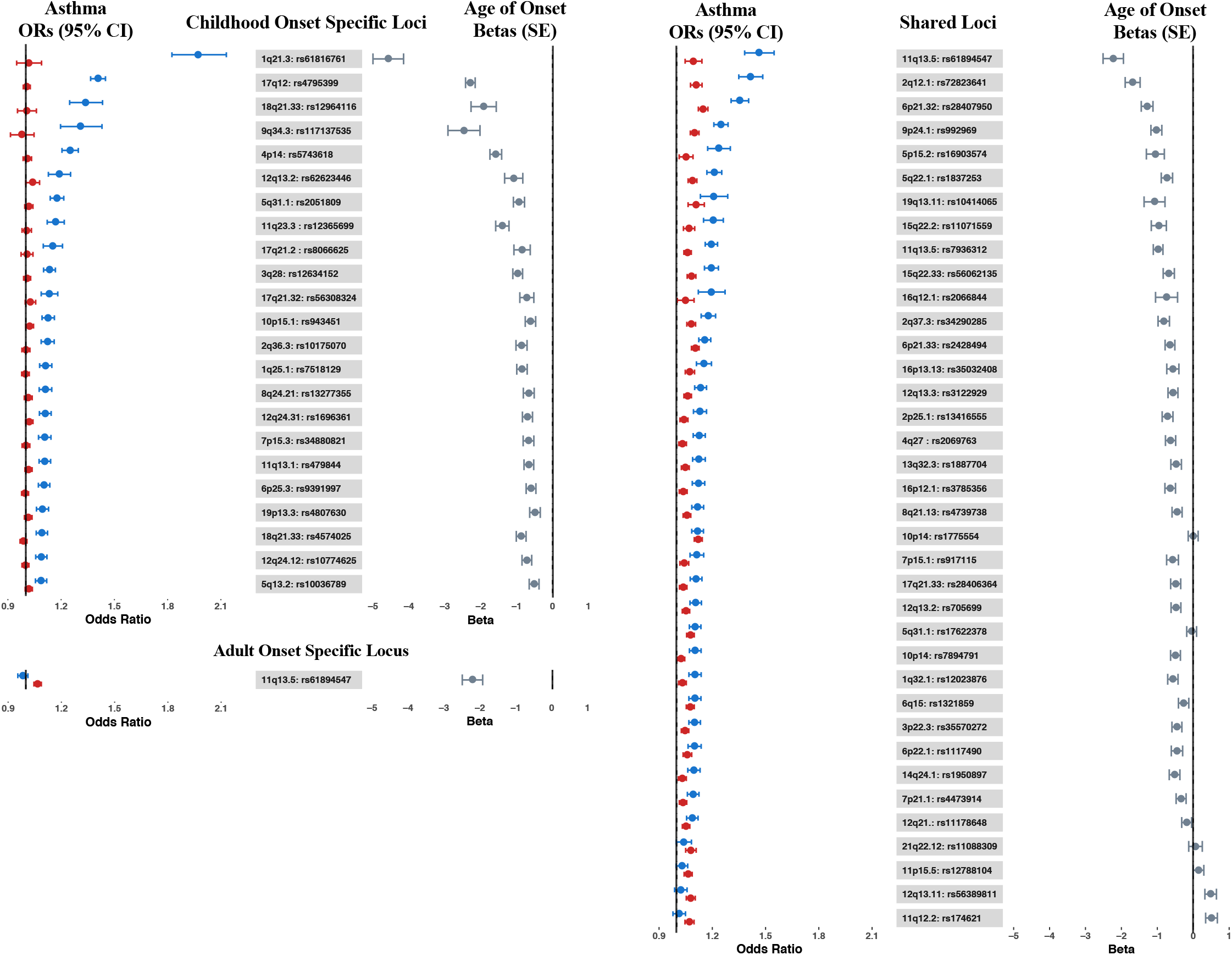
Forest plot showing the odds ratios (ORs) and 95% confidence intervals from the childhood onset (blue) and adult onset (red) GWASs, and betas and standard errors (gray) from the age of onset GWAS for the 61 asthma associated loci. Left panel: Childhood onset specific loci (top) and adult onset specific locus (bottom); right panel: Shared loci. Loci within each group are sorted by OR in the childhood onset GWAS (largest to smallest).

Finally, to directly test for loci associated with asthma age of onset, we conducted a third GWAS including all asthma cases in UKB who met our inclusion criteria (n=37,846). In this analysis, 19 loci were associated with age of onset (p<5×10^−8^) (appendix Figure 6 and Table 5). Age of onset loci overlapped with both the childhood and adult onset specific and shared loci, and asthma risk alleles at all but 2 loci were associated with earlier age of onset (11q12 in the *FADS2* gene and 12q13.11 near the *VDR* gene) (Table 2; Figure 2). SNPs at the 1q21.3 locus (*FLG*) had the largest effect on age of onset, with each copy of the asthma risk allele (rs61816761) associated on average with 4·57 (SE 0·43) years earlier onset compared to individuals without the risk allele (p=8·15×10^−27^). At the 17q12 locus (rs4795399) each copy of the risk allele was associated on average with 2·29 (SE 0·13) years earlier onset compared to individuals without the risk allele (p=6·76×10^−65^). Examples of significant age of onset effects at these and other loci are shown in Figure 3. Overall, both childhood onset specific and shared asthma risk loci were associated with younger ages of onset, and alleles at loci associated with younger ages of onset had larger effects compared to alleles at loci associated with later ages of onset. These results are consistent with a previous GWAS of age of asthma onset in European ancestry subjects that reported five genome-wide significant and three suggestive significant loci, all associated with earlier age of onset^37^. Six of those 8 loci are also associated with age of onset in the UKB GWAS (appendix Table 6).

**Figure 3.**
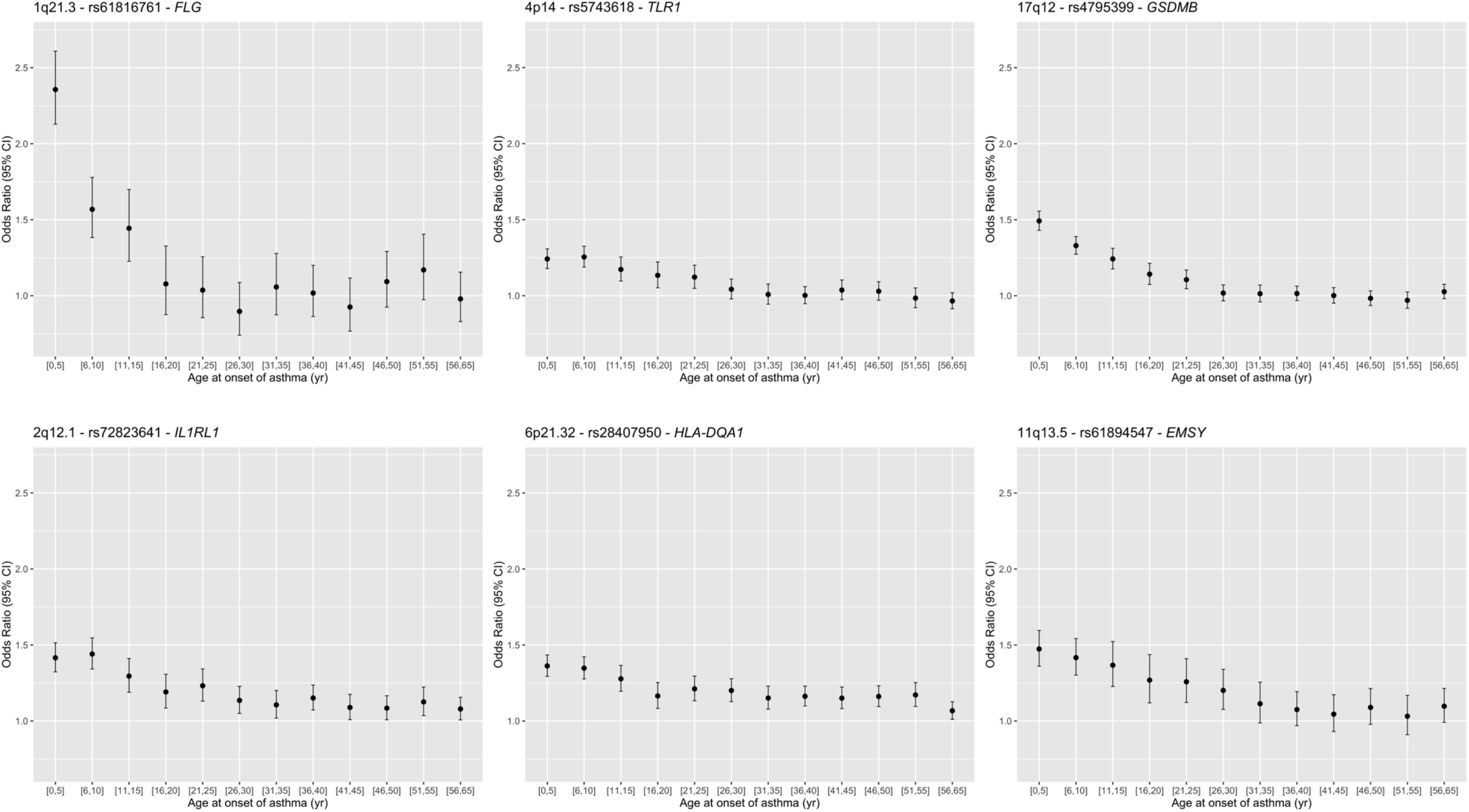
Age of onset effects (ORs) for lead SNPs at three genome-wide signficant childhood onset specific loci (1q21·3, 4p14 and 17q12), and three genome-wide signficant shared loci (2q12·1, 6p21·32, and 11q13·5). The sample sizes for the age of onset bins are [0,5]: n=4,637; [6,10]: n=4,255; [11,15]: n=2,684; [16,20]: n=2,128; [21,25]: n=2,578; [26,30]: n=2,923; [31,35]: n=2,599; [36,40]: n=3,571; [41,45]: n=2,967; [46,50]: n=3,266, [51,55]: n=2,544; [56,65]: n=3,694.

### SNP-based heritability (h^2^)

We used LD score regression to estimate the hertiabilities of childhood onset asthma, adult onset asthma, and age of asthma onset. Consistent with the number of associated SNPs and their effect sizes, estimated heritabilities were 0·33 for childhood onset asthma, 0·098 for adult onset asthma, and 0·14 for age of asthma onset. After excluding the significant SNPs (Table 2), estimates were reduced to 0·21, 0·082, and 0·087, respectively, indicating that the associated SNPs account for 0·11 of the variance in childhood onset asthma risk, 0·016 of the variance in adult onset risk, and 0·049 of the variance in age of asthma onset risk. These results reflect the more significant role for genetic variation in risk for childhood onset compared to adult onset asthma and, conversely, the larger role for environmental variation in risk for adult onset compared to childhood onset asthma.

### Tissue-specific expression of genes at associated loci

Using an unbiased approach, we asked whether the tissue-specific expression of genes that map to the 52 childhood onset loci differed from the tissue-specific expression of genes mapped to the 19 adult onset loci. Genes at childhood onset loci (Figure 1A) were most highly expressed in skin, whole blood, and small intestine (lower ileum) compared to all other tissues, whereas genes at adult onset loci (Figure 1B) were most highly expression in lung, whole blood, small intestine (lower ileum), and spleen (enrichment for higher expression, p<10×10^−3^) (appendix Figure 7 and Table 7). These patterns suggest both overlapping and distinct underlying mechanisms associated with asthma that begins in childhood and asthma with onset in adulthood.

### Predicted transcriptome-wide association test

To better understand molecular mechanisms and to narrow the list of candidate causal genes at associated loci, we focused on the five tissues that most highly expressed the genes at childhood onset or adult onset loci: skin, lung tissue, whole blood, small intestine, and spleen. We used PrediXcan^17^ to identify genes whose expression is predicted by variants associated with asthma in the childhood onset or adult onset GWAS and potentially mediate the effects of associated SNPs on asthma risk.

This analysis identified 113 unique, candidate causal genes at 22 of the 61 GWAS loci (p<1·4×10^−6^) (Figures 4–5) (appendix Figures 8-9 and Table 8). These included 39 genes associated with childhood onset asthma at eight of the childhood onset specific loci and 76 genes associated with childhood and/or adult onset asthma at 13 of the shared loci. Variants at the one adult onset specific locus at 2q22·3 did not predict the expression of any genes in the five tissues.

**Figure 4.**
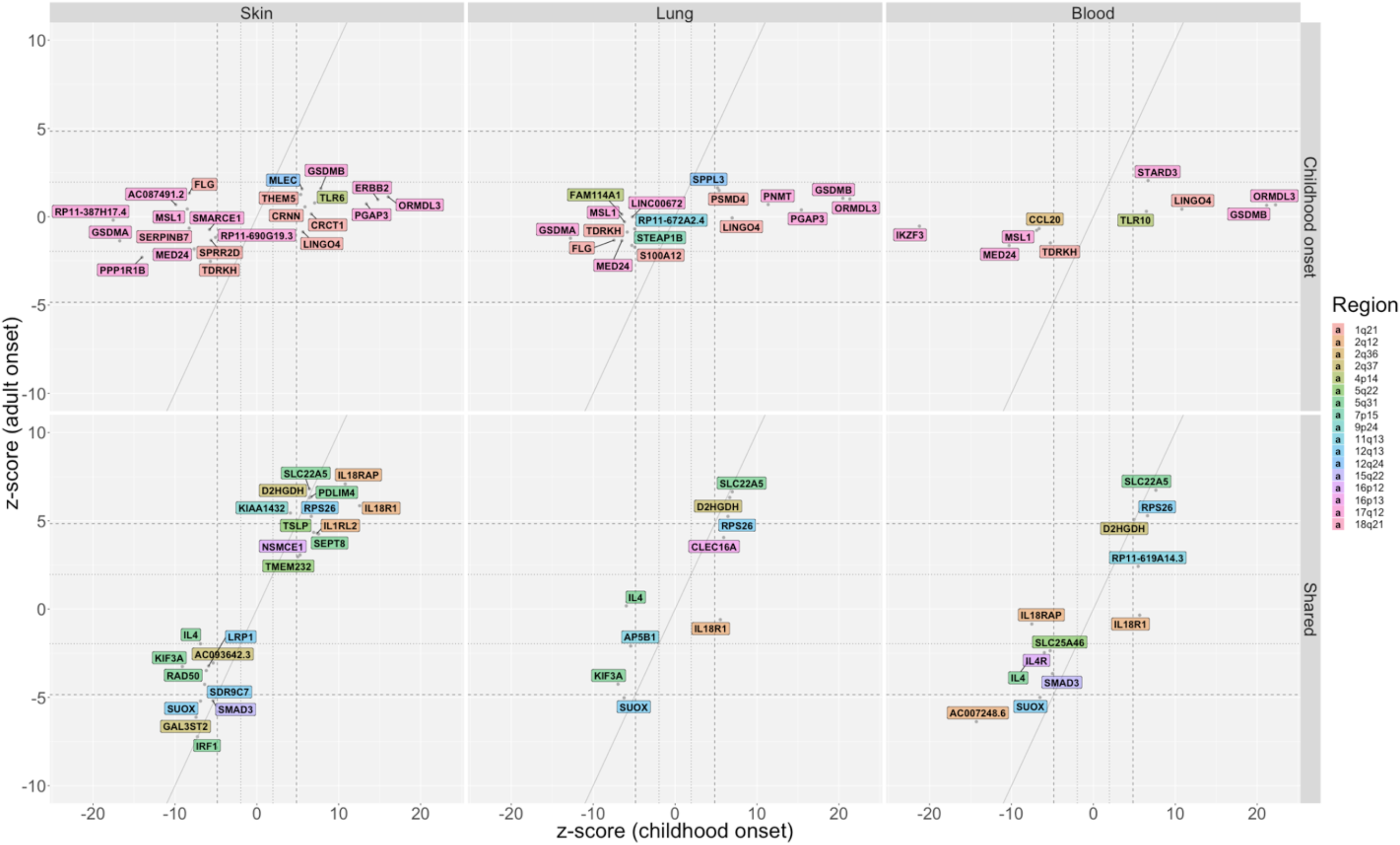
Results of PrediXcan studies at non-HLA region loci. Genes whose predicted expression was significantly associated with asthma in either the childhood onset cases or the adult onset cases are shown for skin (left panels), lung (middle panels) and whole blood (right panels). Values shown for skin combine both sun exposed and not sun exposed skin, showing the most significant statistic of the two. Results using gene expression in spleen and small intestine are shown in appendix Figure 8. The z-scores on the x- and y-axes are from transcriptome-wide tests of association with asthma using SNP sets that predict the expression of that gene. The diagonal dashed lines show the expected when associations are the same in the childhood onset and adult onset cases. The horizontal/vertical dashed lines correspond to z-scores ±4·84 (p=1·29×10^−6^); the horizontal/vertical dotted lines correspond to z-scores ±1·96 (p=0·05). Colored backgrounds correspond to the chromosome location of each gene (see Key). Upper panels: Genes that are associated with childhood onset asthma (no genes were associated with adult onset asthma). Lower panels: Shared genes associated with both childhood onset and adult onset asthma. HLA region genes (6p21·32 and 6p21·33) are shown in Figure 5.

**Figure 5.**
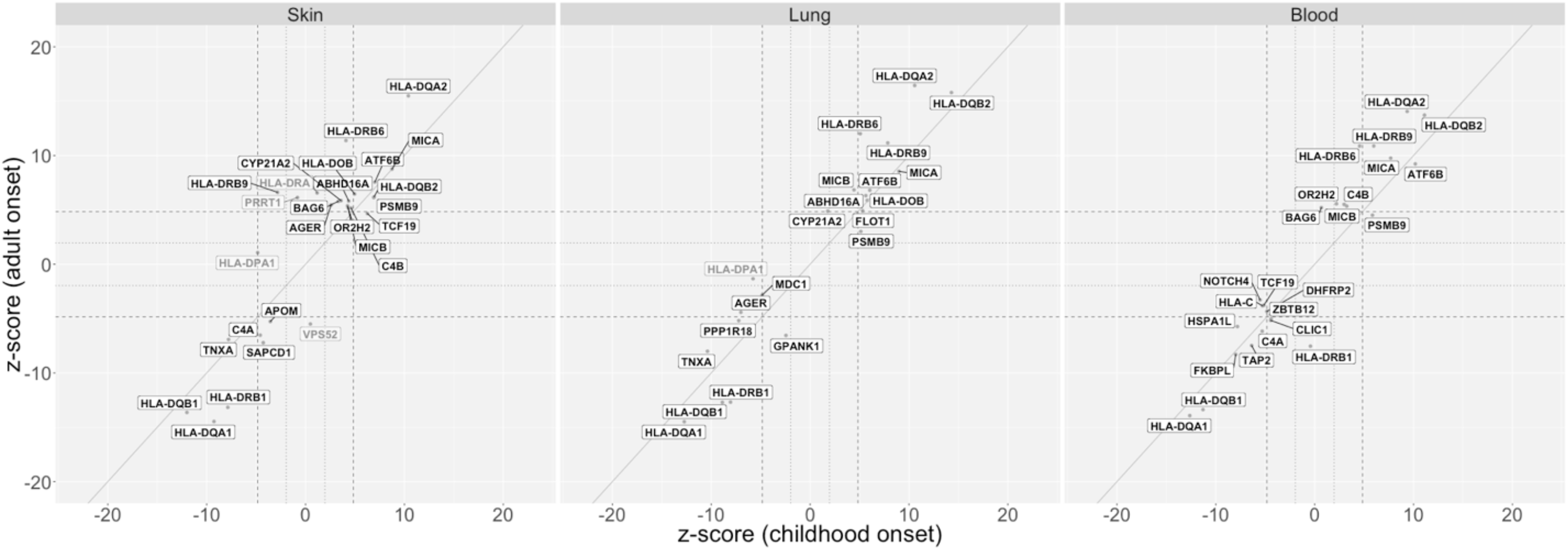
Results of PrediXcan studies of HLA region genes. Genes in the HLA region whose predicted expression was associated with childhood onset or adult onset asthma are shown for skin (left panel), lung tissue (middle panel) and whole blood (right panel). Results using gene expression in spleen and small intestine are shown in appendix Figure 9. The genes on darker shaded backgrounds correspond to shared genes; the four genes on lighter color backgrounds are age-specific. See Figure 4 for additional details.

The predicted genes most significantly associated with childhood onset asthma were at the 17q12 locus (Z score >10) in skin (*ORMDL3*, *ERBB2*, *PGAP3*, *GSDMA*, 2 long noncoding RNAs), lung (*ORMDL3*, *GSDMB*, *GSDMA*, *PGAP3*, *PNMT*), blood (*ORMDL3*, *GSDMB*, *IKZF3*, *MED24*), small intestine (*GSDMA*, *GSDMB*, *PGAP3*), and spleen (*ORMDL3*, *GSDMB*, *ZPBP2*, *MED24*). Some genes were predicted to be more highly expressed in individuals with asthma (e.g., *ORMDL3*, *GSDMB*, *PGAP3*, *ERBB2*), while others were predicted to be less expressed in individuals with asthma (e.g., *GSDMA*, *MED24*, *IKZF3*) (Figure 4A). This pattern of expression reflects the broad regulatory effects of SNPs and tissue specificifity of gene expression at this locus^27^. The childhood onset asthma locus at 1q21.3 includes genes essential for epidermal differentiation and maintaining essential barrier function. The predicted expression of nine genes at this locus were associated with childhood onset asthma. Higher predicted expression of *CRNN*, *CRCT1* and *THEM5* in skin, of *PSDM4* in lung and blood, and of *LINGO4* in skin, lung, and blood were associated with increased asthma risk. Lower predicted expression of *SPRR2D* in skin, of *S100A12* in lung and blood, of *FLG* in skin, lung and spleen, and of *TDRKH* in skin, lung, blood and spleen were associated with increased asthma risk. *S100A12* has been previously implicated in asthma^38^ and *FLG* variants have been associated with atopic dermatitis and food allergies, and asthma in the context of other allergic diseases^21, 28–34^. Other childhood onset specific genes previously implicated in asthma but not previously reported in asthma GWASs are *CCL20*^39^ at 2q36.3 and *TLR10*^40^ at 4p14 in whole blood, and *TLR6*^41, 42^ at 4p14, *AP5B1* at 11q13.1 and *SERPINB7*^43^ at 18q21·33 in skin.

The 5q31.1 region had independent loci that were both childhood onset specific and shared in the GWASs. Although the predicted expression of all eight asthma genes at this extended locus were shared, five genes were more significantly associated with childhood onset asthma: higher predicted expression of *RAD50* in skin but lower predicted expression of *SEPT8* in skin and lung, *IL4* in skin, lung and blood, and *AFF4* small intestine were associated with increased risk for asthma (Figure 4B). The remaining three genes had similar associations with childhood and adult onset asthma, with predicted lower expression of *IRF1* in skin and spleen and predicted higher expression of *PDLIM4* in skin and *SLC22A5* in all five tissues associated with increased asthma risk. *IL4, RAD50, SLC22A5* and *PDLIM4* have been highlighted in previous asthma GWAS^26, 44^, and *KIF3A* was identified in a GWAS of the atopic march^45^ and associated with childhood onset asthma in a candidate gene study^46^.

In contrast to all other loci, predicted expression of 44 genes at two independent shared loci in the HLA region (6p21.32 and 6p21.33; referred to as the HLA region from hereon in) were associated with childhood asthma only (n=1; in skin and lung), adult onset asthma only (n=3; in skin only), or both (n=39; in multiple tissues). (Figure 5). The sheer number of genes in this region with predicted expression associated with asthma, the generally broad tissue expression patterns, and three associated with asthma only in adult onset cases are consistent with this locus being among the two most significant loci in nearly all asthma GWASs, and the most significant locus in a previous small GWASs of adult onset asthma^7^ and in adults with asthma^44, 47^.

Among the remaining 37 shared loci (Figure 2B), SNPs at 12 predicted the expression of 23 unique genes, all of which were associated with both childhood onset and adult onset asthma. These include *IL18R1*, *IL18RAP*, and *ILRL2* at 2q12·1 in multiple tissues, *TSLP* at 5q22.1 in skin, *SMAD3* at 15q22·33 in skin and blood, *LRP1* at 12q13·3 in skin, *IL4R* at 16p12.1 in blood, and *CLEC16A* at 16p13.13 in lung. Loci associated with *IL18R1*, *IL18RAP*, *IL18RAP, ILRL2, TSLP, SMAD3*, *LRP1*, *IL4R* and *CLEC16A* were reported in previous asthma GWAS^21, 26, 36^.

## Discussion

We report here the first large GWAS of both childhood and adult onset cases. To both maximize our power to detect differences and minimize the likelihood of misclassification, we considered doctor diagnosed asthma before the age of 12 years as childhood onset asthma and doctor diagnosed asthma after the age of 25 years as adult onset asthma. These GWASs revealed 61 independent asthma loci, 23 specific to childhood onset, one specific to adult onset, and 37 shared; with overall larger effect sizes for childhood onset asthma at nearly all loci. Moreover, the predicted expression of 41 of the 113 implicated genes were associated specifically with childhood onset asthma, compared to the predicted expression of three genes associated specifically with adult onset asthma. Our findings of more childhood onset asthma loci and potentially causal genes, and the larger effect sizes of risk alleles in childhood onset cases are particularly striking given that there were nearly 2·5-times more adult onset than childhood onset cases in this study. Thus, despite having substantially less power to detect loci specific to childhood onset asthma, our analyses revealed many more childhood onset asthma loci. Similarly, the asthma risk alleles at 19 loci that were significant in the age of onset GWAS were all associated with younger age of onset. Finally, we showed that the SNP-based heritability of childhood onset asthma is over 3-times larger than the SNP-based heritability of adult onset asthma. These findings are consistent with previous studies showing decreased estimates of asthma heritability with increasing age of onset^48^, age of onset SNPs associated with earlier age of onset^37^, and an additive, unweighted genetic risk score comprised of 15 SNPs at eight asthma-associated loci associated with earlier age of onset^49^. Our study further shows that genetic risk for adult onset asthma is largely a subset of the genetic risk loci for childhood onset asthma, but with overall smaller effect sizes, consistent with a larger role for environmental risk factors in adult onset asthma.

Despite the overlap of adult onset and childhood onset loci, distinct mechanisms contributing to each were suggested by tissue enrichments: childhood onset loci were enriched for genes with highest expression in skin whereas adult onset loci were enriched for genes with highest expression in lung and spleen; both were enriched for genes highly expressed in whole blood and small intestine. The highlighting of skin as a target tissue for childhood onset asthma supports the widely held idea that asthma in childhood is due to impaired barrier function in the skin and other epithelial surfaces. This model proposes that compromised epithelial barriers promote sensitization to food and airway allergens and to wheezing illnesses in early life^35, 50^. In fact, childhood onset specific loci identified here have been associated with atopic dermatitis or food allergies, such as *FLG* on 1q21·3 with the atopic march^45^, atopic dermatitis^31–33^ and food allergies^28–30^, *KIF3A* on 5q31.1 and *AP5B1*/*OVOL1* on 11q13.1 with the the atopic march^45^ and atopic dermatitis^51^, *SERPINB7* on 18q21.33 with food allergies^43^, and *CRNN* (cornulin) on 1q21·3 with atopic dermatitis concomitant with asthma and reduced expression in atopic dermatitis-affected skin^52^. Variants at those loci were all associated with earlier age of asthma onset. We further show that these loci are associated with childhood onset asthma, even after exclusion of cases with a history of allergic diseases. In contrast, the enrichment for genes highly expressed in lung and spleen at adult onset loci suggests a more lung-centered, and potentially immune mediated, etiology for asthma with onset later in life. The prominant role of the HLA region in the adult onset asthma GWAS and the fact that predicted expression of three HLA region genes was associated only with adult onset asthma further highlights a central role for immune processes driving asthma pathogenesis in adults. The fact that both childhood onset and adult onset asthma loci were enriched for genes that are most highly expressed in whole blood cells and small intestine further indicate a shared immune etiology, as suggested from a large GWAS that included both children and adults^26^.

Combining GWAS with a transcriptome-wide association test that uses combinations of associated SNPs to predict gene expression in different tissues revealed significant complexity at the two most highly associated asthma loci. SNPs at the 17q12 locus predicted expression of 18 childhood onset asthma genes and SNPs at the HLA region predicted expression of 42 genes: three were associated with adult onset asthma and most were not HLA genes *per se*. In this regard, it is notable that the *HLA-DRB1*, *HLA-DQB1*, and *HLA-DQA1* genes, which are the most associated HLA loci with autoimmune diseases, are predicted to have reduced expression in both childhood onset and adult onset asthma. Instead, HLA genes with less clear functions have increased predicted expression in asthma (Figure 5). These results strengthen the argument that multiple genes contribute to asthma risk at the HLA and 17q12 loci and probably account for the highly significant GWAS p-values observed at these loci in nearly all studies. It is also likely that these genes have both tissue specific and broad effects in epithelium, lung, and immune tissues.

The new loci identified in our study include the first adult onset asthma specific association at 2q22.3. The lead SNP at 2q22.3 is intergenic between *TEX41* and *ACVR2A*. The predicted expression of *ACVR2A* was not associated with asthma in our study, despite it being expressed in lung, blood, small intestine and spleen. *TEX41* was not expressed in any of the five tissues investigated. Interestingly, a GWAS also performed in UKB subjects implicated variants near *TEX41* in heavy vs. never smoking behavior^53^. However, even after removing adult onset cases and controls with reported ‘ever smoking’, the p-value for this SNP remained significant and the OR slightly increased (OR 1·077 [95% CI 1·05, 1.1], p=2·26×10^−8^; n=12,132 cases and 176,704 controls). Variants in or near this gene, which encodes a lincRNA, have been associated with cardiovascular and immune mediated traits^54^, making this a potentially interesting candidate gene for adult onset asthma.

Our study had limitations. First, diagnoses of asthma and allergic disease in study subjects were from self reported doctor diagnosis and medical records (ICD10 codes). Thus, it is possible that diagnoses, age of onset, or both are misspecified in some subjects. On the one hand, the large sample size and our ability to replicate nearly all previously reported asthma loci (appendix Tables 2 and 9) suggest that our analyses were robust to any inaccuracies in the data. On the other hand, it is possible that subjects with adult onset asthma included cases with poor recall of childhood onset asthma in which symptoms remitted and then relapsed later in life^55^ or misclassified cases of COPD among the older age groups. Our sensitivity analysis suggested that if even as few as 5% of the adult onset cases were misclassified we should have observed some signal of association at childhood onset loci, which we did not. The fact that the odds ratios for asthma at shared loci are relatively similar from approximately age 25 to age 65 (Figure 3) and that we do not detect any association signal at the major COPD locus on chromosome 15q25.1 (appendix Figure 10), further suggests that there is negligible misclassification of cases in the older age groups. Second, although we used stringent criteria to classify loci as childhood or adult onset specific, we can’t exclude the possibility that in infinitely large sample sizes the effect sizes of some of these loci will have 95% CIs that overlap or the association p-value will become smaller than 0·05. Conversely, some of the shared loci with modest p-values in the adult onset cases may not be true risk loci for asthma with onset at older ages. Third, the gene expression data used to predict candidate target genes included heterogeneous tissues and were collected mostly from adults. As a result, our study may have missed relevant genes whose expression is developmentally regulated or environment specific. Our finding of candidate genes at only 22 of the 61 asthma loci may be due in part to the importance of both in asthma pathogenesis. Moreover, all inference based on gene expression is using imputed expression. It is possible, therefore, that some relevant genes were more difficult to impute and not included in our analysis, although a recent comparative study showed that PrediXcan is a more robust method for prediction of gene expression than other related methods^56^. Fourth, because of the ethnic composition of UKB, this study was limited to individuals of European ancestry only. As a result, we could not evaluate the genetic risk architecture or assess the effects of age of onset specific loci in other populations.

In the largest asthma GWAS to date, we show that genetic risk loci for adult onset asthma is largely a subset of the loci associated with childhood onset asthma, with overall smaller effect sizes for onset at later ages. These data suggest that childhood onset specific loci and those associated with age of onset play a role in disease initiation, whereas the other associated loci reflect shared mechanisms of disease progression. The differences in the target tissues that most highly express the genes at associated loci and the predicted expression of genes at age specific and shared loci provides additional genetic and molecular evidence for both shared and distinct pathogenic mechanisms in childhood onset and adult onset asthma. It is therefore possible that the most effective treatments will also differ between these two groups, and that strategies for precision medicine should be further personalized to account for age of asthma onset.

## Author’s Contributions

All authors were involved in the conception and design of the study and in writing the manuscript. MP and NS conducted all analyses and prepared figures and tables, under the overall supervision of DLN, CO, and HKI.

## URLs

UK Biobank, [https://www.ukbiobank.ac.uk/]

Hail: https://github.com/hail-is/hail

## Acknowledgements

This research has been conducted using UK Biobank Resource under Application Number 19526.

