## Supplementary material (text, tables, figures) for "Shared and Distinct Genetic Risk Factors for Childhood Onset and Adult Onset Asthma: Genome- and Transcriptome-wide Studies"

### Methods

#### Quality control of samples

We relied on quality control procedures performed centrally by the UK Biobank (UKB). We selected individuals from the White British Ancestry Set and filtered out those with poor-quality genotypes (high heterozygosity and missing rates)[1]. In addition, we excluded individuals with ambiguous sex assignments using X and Y chromosome SNPs; for pairs of individuals who were related as first or second-degree relatives based on genotypes, only the individual with the smallest number of missing genotypes was included. This resulted in 376,358 unrelated individuals with high-quality genotype for our study.

#### Phenotype and Group definitions

We defined five different groups for this study: childhood onset asthma cases, childhood onset asthma cases excluding individuals with allergic disease, adult onset asthma cases, and two sets of controls (with and without allergic disease).

Childhood onset asthma cases consisted of individuals with onset prior to 12 years of age (n=9,433), using the UKB data field 22147 (*Age asthma diagnosed by doctor*, Online follow-up) and data field 3786 (*Age asthma diagnosed*, Touchscreen). A similar group was defined excluding allergic rhinitis, atopic dermatitis and food allergy (n=6,228), using self-reported data in data fields 20002 (*Non-cancer illness code*, Verbal interview), 22126 (*Doctor diagnosed hayfever or allergic rhinitis*, Online follow-up) and 22146 (*Age hayfever or allergic rhinitis diagnosed by doctor*, Online follow-up), as well as hospital-level data 41202/41204 (main and secondary diagnoses using ICD10 codes L208, L209, J301, J302, J303 and J304). The adult onset asthma cases group was defined as individuals with onset after 25 and before 66 years of age (n=21,564), excluding individuals with chronic obstructive airways disease (COPD) either self-reported using UKB data fields 20002, 22130 (*Doctor diagnosed COPD*, Online follow-up), 22150 (*Age COPD diagnosed by doctor*, Online follow-up), 22128 (Doctor diagnosed emphysema, Online follow-up), 22129 (Doctor diagnosed chronic bronchitis, Online follow-up), 22148 (Age emphysema diagnosed by doctor, Online follow-up), 22149 (Age chronic bronchitis diagnosed by doctor, Online follow-up) and 3992 (Age emphysema/chronic bronchitis diagnosed, Online follow-up), or with COPD diagnosed by a doctor (hospital-level data) using data fields 41202/41204 (ICD10 codes J43, J430, J431, J432, J438, J439, J44, J440, J441, J448 and J449).

The control group was defined as individuals not having neither asthma nor COPD (n=318,237) using self-reported and hospital-level data (ICD10 codes used to exclude individuals with an asthma diagnosis are J45, J450, J451, J458 and J459). A second controls group with no allergy (n=284,429) was also defined with the same exclusions described above.

For the GWAS of asthma age of onset, all asthma cases with onset before 66 years of age were included (n=37,846).

#### Reproducible research – Phenotype query file

In the previous section we provide a description of our quality control procedures for samples and the phenotypes definitions. To further improve the reproducibility of our study, we also provide here the query file used to retrieve exactly the same data employed in our analysis. The UK Biobank, as other biobanks, is composed of large amounts of highly heterogeneous data types, different sources, and structure. To address to many challenges posed by these characteristics, we developed a tool called ukbREST (https://github.com/hakyimlab/ukbrest)[2] that processes the UK Biobank data, creating a private and secure server for easy and quick access for authorized users within a protected cloud (to be clear, the data is NOT made publicly available). This resource allows streamlined updating and programmatic, simple, and reproducible phenotype extraction within the private cloud the investigator is using. As a REST API implementation, it allows researchers to efficiently access subsets of SNPs and phenotypic traits, leveraging a BGEN indexer and an SQL database. We have used this tool to efficiently retrieve data to distribute jobs performing phenome-wide gene level association with PrediXcan/MetaXcan to thousands of phenotypes, leveraging a secure biomedical cloud called Bionimbus Protected Data Cloud (PDC)[3], which operates at FISMA moderate as IaaS with an NIH Trusted Partner status for analyzing protected datasets.

Our ukbREST tool allows us to write a descriptive query file in YAML that can be submitted to the private server for retrieval of the phenotype data requested. At the top of the file the user specifies the quality control criteria for samples (section "sample_filters" at the top), and then a series of sections that can be individually downloaded as separate files. The previous descriptions of the phenotypes are here shown as the precise SQL statements to get the exact set of data employed in this study.

**ukbREST phenotype query file.** This YAML file specifies the data to be pulled from the UK Biobank database stored in our in-house protected database server (Bionimbus)

```yaml
samples_filters:
- cin_white_british_ancestry_subset_0_0 = 1
- eid not in (select eid from bad_related_samples_2nd_higher_and_high_missrate)
- chet_missing_outliers_0_0 = 0
- cputative_sex_chromosome_aneuploidy_0_0 = 0
- cexcess_relatives_0_0 = 0
- eid not in (select eid from withdrawls)
- eid > 0

extras:
  sex: c31_0_0
  age_recruitment: c21022_0_0
  asthma_medication: c22167_0_0
  smoking_status: coalesce(nullifneg(c20116_2_0::int), nullifneg(c20116_1_0::int), nullifneg(c20116_0_0::int))
  fev1: array_avg(array[c3063_0_0, c3063_0_1, c3063_0_2])
  fev1_pred: c20153_0_0
  fev1_pred_perc: c20154_0_0
  fvc: array_avg(array[c3062_0_0, c3062_0_1, c3062_0_2])
  fev1_fvc_ratio: array_avg(array[c3063_0_0, c3063_0_1, c3063_0_2]) / array_avg(array[c3062_0_0, c3062_0_1, c3062_0_2])
  fev1pred_fvc_ratio: c20153_0_0 / array_avg(array[c3062_0_0, c3062_0_1, c3062_0_2])
  has_eczema:
    sql:
      1: |
        (
          eid in (
            select eid from events where field_id = 20002 and event in (values('1452'))
            union
            select eid from events where field_id in (values(41202), (41204)) and event in (values('L208'), ('L209'))
          )
        )
  has_hayfever:
    sql:
      1: |
        (
          eid in (
            select eid from events where field_id = 20002 and event in (values('1387'))
            union
            select eid from events where field_id in (values(41202), (41204)) and event in (values('J301'), ('J302'), ('J303'), ('J304'))
          )
          or
          (c22126_0_0 = '1')
          or
          (c22146_0_0 is not null)
        )
  has_food_allergy:
    sql:
      1: |
        eid in (select eid from events where field_id = 20002 and event in (values('1385')))

simple_covariates:
  sex: c31_0_0
  pc1: cpc1_0_0
  pc2: cpc2_0_0
  pc3: cpc3_0_0
  pc4: cpc4_0_0
  pc5: cpc5_0_0
  pc6: cpc6_0_0
  pc7: cpc7_0_0
  pc8: cpc8_0_0
  pc9: cpc9_0_0
  pc10: cpc10_0_0

data_aliases:
- &asthma_age_onset |
  case when
    -- children
    (
      coalesce(nullifneg(c22147_0_0), nullifneg(c3786_2_0), nullifneg(c3786_1_0), nullifneg(c3786_0_0)) < 12
    )
    or
    (
      coalesce(nullifneg(c22147_0_0), nullifneg(c3786_2_0), nullifneg(c3786_1_0), nullifneg(c3786_0_0)) BETWEEN 12 and 25
    )
    or
    (
      coalesce(nullifneg(c22147_0_0), nullifneg(c3786_2_0), nullifneg(c3786_1_0), nullifneg(c3786_0_0)) BETWEEN 26 and 65

      and

      -- exclude copd
      eid not in (select eid from events where field_id = 20002 and event in (values('1112')))
      and
      (c22130_0_0 is null or c22130_0_0::int = 0)
      and
      nullifneg(c22150_0_0) is null
      and
      eid not in (select eid from events where field_id in (values(41202), (41204)) and event in (values('J44'), ('J440'), ('J441'), ('J448'), ('J449')))

      and

      -- emphysema/chronic bronchitis
      eid not in (select eid from events where field_id = 20002 and event in (values('1113'), ('1412'), ('1472')))
      and
      (c22128_0_0 is null or c22128_0_0::int = 0) and (c22129_0_0 is null or c22129_0_0::int = 0)
      and
      nullifneg(c22148_0_0) is null and nullifneg(c22149_0_0) is null
        and
        coalesce(nullifneg(c3992_0_0), nullifneg(c3992_1_0), nullifneg(c3992_2_0)) is null
      and
      eid not in (select eid from events where field_id in (values(41202), (41204)) and event in (values('J43'), ('J430'), ('J431'), ('J432'), ('J438'), ('J439')))
    )
  then
    coalesce(nullifneg(c22147_0_0), nullifneg(c3786_2_0), nullifneg(c3786_1_0), nullifneg(c3786_0_0))
  else
    NULL
```

```
  end

- &asthma_children |
  coalesce(nullifneg(c22147_0_0), nullifneg(c3786_2_0), nullifneg(c3786_1_0), nullifneg(c3786_0_0)) < 12

- &asthma_children_no_allergic |
  (
    coalesce(nullifneg(c22147_0_0), nullifneg(c3786_2_0), nullifneg(c3786_1_0), nullifneg(c3786_0_0)) < 12
  )
  and
  (
    eid not in (select eid from events where field_id = 20002 and event in (values('1452'), ('1387'), ('1385')))
    and
    eid not in (select eid from events where field_id in (values(41202), (41204)) and event in (
      values('L208'), ('L209'),
      ('J301'), ('J302'), ('J303'), ('J304')
    ))
    and
    (c22126_0_0 = '0' or c22126_0_0 is null)
    and
    c22146_0_0 is null
  )

- &asthma_adults |
  (
    coalesce(nullifneg(c22147_0_0), nullifneg(c3786_2_0), nullifneg(c3786_1_0), nullifneg(c3786_0_0)) BETWEEN 26 and 65
  )
  and
  (
    -- exclude copd
    eid not in (select eid from events where field_id = 20002 and event in (values('1112')))
    and
    (c22130_0_0 is null or c22130_0_0::int = 0)
    and
    nullifneg(c22150_0_0) is null
    and
    eid not in (select eid from events where field_id in (values(41202), (41204)) and event in (values('J44'), ('J440'), ('J441'), ('J448'), ('J449')))

    and

    -- emphysema/chronic bronchitis
    eid not in (select eid from events where field_id = 20002 and event in (values('1113'), ('1412'), ('1472')))
    and
    (c22128_0_0 is null or c22128_0_0::int = 0) and (c22129_0_0 is null or c22129_0_0::int = 0)
    and
    nullifneg(c22148_0_0) is null and nullifneg(c22149_0_0) is null
      and
      coalesce(nullifneg(c3992_0_0), nullifneg(c3992_1_0), nullifneg(c3992_2_0)) is null
    and
    eid not in (select eid from events where field_id in (values(41202), (41204)) and event in (values('J43'), ('J430'), ('J431'), ('J432'), ('J438'), ('J439')))
  )

- &controls |
  (
    coalesce(nullifneg(c22147_0_0), nullifneg(c3786_2_0), nullifneg(c3786_1_0), nullifneg(c3786_0_0)) is null
    and
    eid not in (select eid from events where field_id in (values(41202), (41204)) and event in (values('J45'), ('J450'), ('J451'), ('J458'), ('J459')))
  )
  and
  (
    -- copd
    eid not in (select eid from events where field_id = 20002 and event in (values('1112')))
    and
    (c22130_0_0 is null or c22130_0_0::int = 0)
    and
    nullifneg(c22150_0_0) is null
    and
    eid not in (select eid from events where field_id in (values(41202), (41204)) and event in (values('J44'), ('J440'), ('J441'), ('J448'), ('J449')))

    and

    -- emphysema/chronic bronchitis
    eid not in (select eid from events where field_id = 20002 and event in (values('1113'), ('1412'), ('1472')))
    and
    (c22128_0_0 is null or c22128_0_0::int = 0) and (c22129_0_0 is null or c22129_0_0::int = 0)
    and
    nullifneg(c22148_0_0) is null and nullifneg(c22149_0_0) is null
      and
      coalesce(nullifneg(c3992_0_0), nullifneg(c3992_1_0), nullifneg(c3992_2_0)) is null
    and
    eid not in (select eid from events where field_id in (values(41202), (41204)) and event in (values('J43'), ('J430'), ('J431'), ('J432'), ('J438'), ('J439')))
  )

- &controls_no_allergic |
  (
    coalesce(nullifneg(c22147_0_0), nullifneg(c3786_2_0), nullifneg(c3786_1_0), nullifneg(c3786_0_0)) is null
    and
    eid not in (select eid from events where field_id in (values(41202), (41204)) and event in (values('J45'), ('J450'), ('J451'), ('J458'), ('J459')))
  )
  and
  (
    -- copd
    eid not in (select eid from events where field_id = 20002 and event in (values('1112')))
    and
    (c22130_0_0 is null or c22130_0_0::int = 0)
    and
    nullifneg(c22150_0_0) is null
    and
    eid not in (select eid from events where field_id in (values(41202), (41204)) and event in (values('J44'), ('J440'), ('J441'), ('J448'), ('J449')))

    and

    -- emphysema/chronic bronchitis
    eid not in (select eid from events where field_id = 20002 and event in (values('1113'), ('1412'), ('1472')))
    and
    (c22128_0_0 is null or c22128_0_0::int = 0) and (c22129_0_0 is null or c22129_0_0::int = 0)
    and
    nullifneg(c22148_0_0) is null and nullifneg(c22149_0_0) is null
      and
      coalesce(nullifneg(c3992_0_0), nullifneg(c3992_1_0), nullifneg(c3992_2_0)) is null
    and
    eid not in (select eid from events where field_id in (values(41202), (41204)) and event in (values('J43'), ('J430'), ('J431'), ('J432'), ('J438'), ('J439')))
```

```
)
and
(
 eid not in (select eid from events where field_id = 20002 and event in (values('1452'), ('1387'), ('1385')))
 and
 eid not in (select eid from events where field_id in (values(41202), (41204)) and event in (
   values('L208'), ('L209'),
   ('J301'), ('J302'), ('J303'), ('J304')
 ))
 and
 (c22126_0_0 = '0' or c22126_0_0 is null)
 and
 c22146_0_0 is null
)

data:
 asthma_age_onset: *asthma_age_onset

 asthma_children:
  sql:
   1: *asthma_children
   0: *controls

 asthma_children_no_allergic:
  sql:
   1: *asthma_children_no_allergic
   0: *controls_no_allergic

 asthma_adults:
  sql:
   1: *asthma_adults
   0: *controls
```

#### Quality control of genotype data

Details on genotyping, imputation and QC of UK Biobank genotype are described in Bycroft[1]. Briefly, after sample and marker QC, a genotype dataset containing 488,377 samples and 803,426 markers from two arrays (UK BiLEVE Axiom Array and UK Biobank Axiom Array) was imputed using the Haplotype Reference Consortium (HRC) and UK10K haplotype resource increasing the total number of variants to 96 million.

We performed additional QC to ensure high quality genotypes by requiring variants to have an imputation quality score (INFO) > 80%, a minor allele frequency (MAF) > 0·1%, call rate > 95%, and a Hardy-Weinberg equilibrium P-value greater than $1x10^{-10}$. This resulted in a final set of 10,894,596 variants for analyses.

#### Sensitivity analysis to adult onset misclassification

To assess the proportion of childhood onset cases among the adult onset cases needed to detect associations with SNPs at the childhood onset loci, we replaced adult onset cases with childhood onset cases in varying proportions and repeated analyses of association with the lead SNP at the most associated locus (rs4795399 in *GSDMB* on 17q12-21; left panel of figure below) and the lead SNP at the locus with the largest effect size (rs61816761 in *FLG* on 1q21·3; right panel). We considered proportions from 0% to 35% replacements with randomly sampled childhood onset cases (x-axis). For each proportion, we repeated the procedure 50 times to quantify the sampling variability; the median and 25th and 75th percentiles of the $-\log_{10}$ p-values are shown. The black horizontal lines show the genome-wide significance threshold ($5x10^{-8}$) and the nominally significant threshold (0·05). The p-value for each SNP in the UK Biobank adult onset asthma GWAS is shown as 0% contamination. At the 17q locus (left panel), the p- values (y-axis) become genome-wide significant when 15% or more of the sample is composed of childhood onset cases; at 5% contamination the median p-value for the 17q SNP in the "adult onset" sample is 0·039 (interquartile range [IQR] = 0·031), which is considerably more significant than the p-value in the adult onset cases in our study (0·628). At the 1q locus (right panel), the p-values (y-axis) become genome-wide significant when 20% or more of the sample is composed of childhood onset cases; at 5% contamination the median p-value for the 1q SNP in the "adult onset" sample is 0·094 (IQR = 0·065), which is considerably more significant than the p-value in the adult onset cases as defined in our study (0·656). These data indicate that relatively few (~5%) childhood onset cases would be sufficient to generate some signal of association at these two childhood onset loci in the adult onset cases. The fact that we did not suggests that the adult onset cases have little "contamination" with childhood asthma cases.

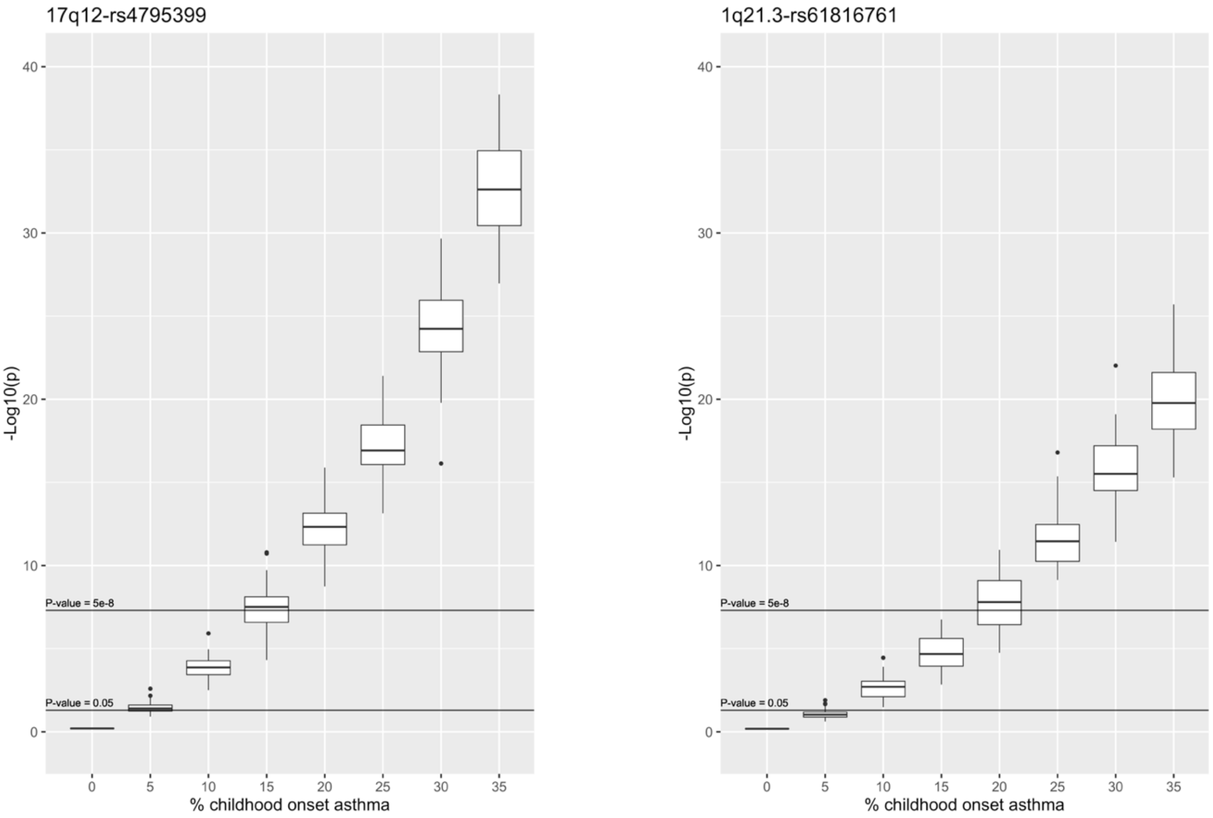

#### Permutation testing of shared risk loci

To compare the odds ratios (OR) between childhood onset and adult onset asthma GWAS we simulated shared variants under the null hypothesis of no difference in effect size between childhood and adult onset cases. To simulate effect sizes under the null hypothesis, we reshuffled (permuted) the childhood and adult onset cases.

We sampled 9,433 individuals from the combined pool of childhood and adult onset cases and labelled them as childhood onset cases. Similarly, we sampled 21,564 individuals and labelled them as adult onset cases. By sampling from the combined pool, we ensured the null hypothesis of no difference between the two classes. The sampling was done with replacement. We used the same 318,237 controls as for the main GWASs.

For the genotypes, we used the 39 shared SNPs from our main analysis (Table 2C in manuscript). We performed logistic regression of childhood onset cases vs controls against each of the 39 SNPs separately. The effect sizes were implicitly established by the true underlying effect sizes. We repeated this sampling 100 times resulting in 3900 representative SNPs. 739 SNPs were genome-wide significant ($P<5x10^{-8}$) in both the childhood onset and the adult onset permuted "cases" and therefore considered to be shared.

To generate the null distribution of median differences between childhood and adult onset loci, we further resampled 10,000 sets of 39 SNPs and computed the median difference of their log odds ratios. Finally, we compared the observed median difference between childhood onset and adult onset log odds ratios to this null distribution as plotted in the following figure. The *P*-value was calculated as the number of times the observed median exceeded the simulated median differences. The observed median value (0·074) is shown by the vertical black line on the figure below (permutation test $P<10^{-4}$).

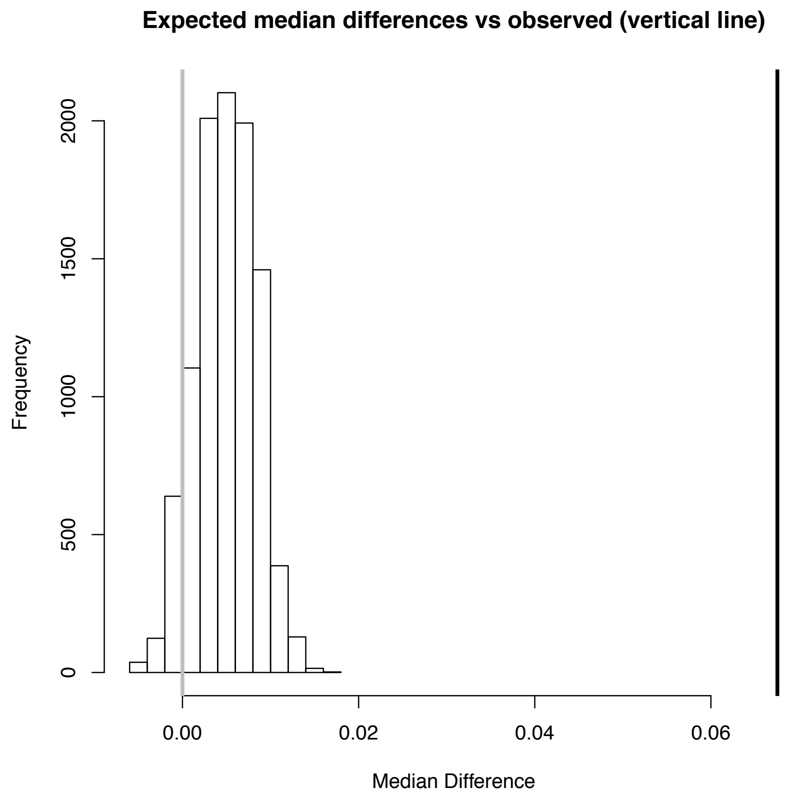

#### PrediXcan and S-PrediXcan

PrediXcan correlates predicted expression levels with phenotypes to find genes that may mediate the genotype-phenotype association. Here we use the summary statistic version, S-PrediXcan, which uses GWAS results and prediction weights to directly infer the gene-level association statistics. The weights were downloaded from our repository http://predictdb.org, which serves prediction models from GTEx and other reference transcriptome datasets. These are based on elastic net models with mixing parameter 0·5, which we have shown to produce sparse and robust prediction[4].

#### Sensitivity analysis of COPD and asthma age of onset

To further the potential misclassification of COPD as adult onset asthma, we performed two sensitivity analyses (GWAS): 1) we removed adults with ages of asthma onset between 46 to 65 years, and 2) we removed adult-onset cases and controls with FEV1/FVC <0·70. The first GWAS included 12,060 cases with onset between ages 25 and 45 years, who were compared to all controls. The second GWAS included 14,409 cases and 252,199 controls with FEV1/FVC ≥0·70 from among the 18,930 adult onset asthma cases and 294,043 controls with spirometry data (FEV1 and FVC). We then compared the results of these two GWASs with the GWAS using the full set of adult onset cases and controls. As shown in the figures below, the ORs and p-values from each GWAS are highly correlated ($r = 0{\cdot}76$ for both ORs and p-values in sensitivity analysis #1, and $r = 0{\cdot}82$ for both ORs and p-values in sensitivity analysis #2). However, both yielded ORs that were slightly larger in the subset analyses. This suggests some dilution of the genetic effect by both the older cases and those with FEV1/FVC <0·70 (who have a slightly later onset). This could be due to the decreasing heritability with age on onset or to contamination with COPD cases. However, the fact that significance increases overall when the excluded individuals are included, reassures us that most of these cases are likely to have asthma. Moreover, the fact that we do not see any evidence for association in the adult onset GWAS at the prominent COPD locus on chromosome 15q25.1 (see appendix Figure 9) further suggests that contamination with COPD cases is not a major issue in our analysis and that the lower ORs in the adult onset compared GWAS to childhood onset GWAS (see Figure 2) is not due to misdiagnosis in the adult onset cases. In fact, these differences would be even greater if we focused only on younger adults or excluded those with poor lung function.

Comparisons of the GWAS odds ratios (left) and p-values (right) from the complete adult onset group (x-axis) vs adult onset cases with age of onset from 26 to 45 years (y-axis). The correlations for both panels are 0·76.

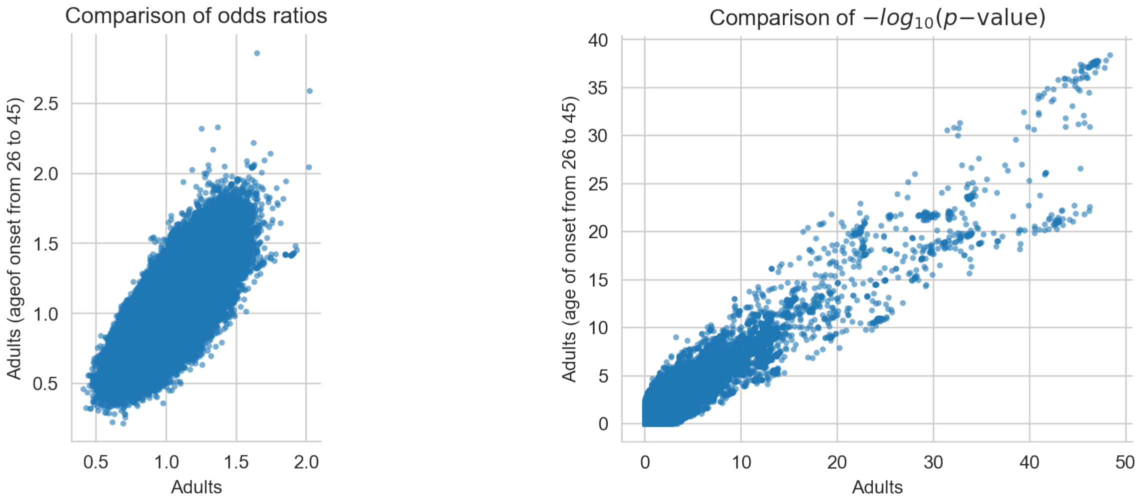

Comparisons of GWAS odds ratios (left) and p-values (right) from the complete adult onset group (x-axis) vs adult onset cases with FEV1/FVC < 0·70 (y-axis). The correlations for both panels are 0·82.

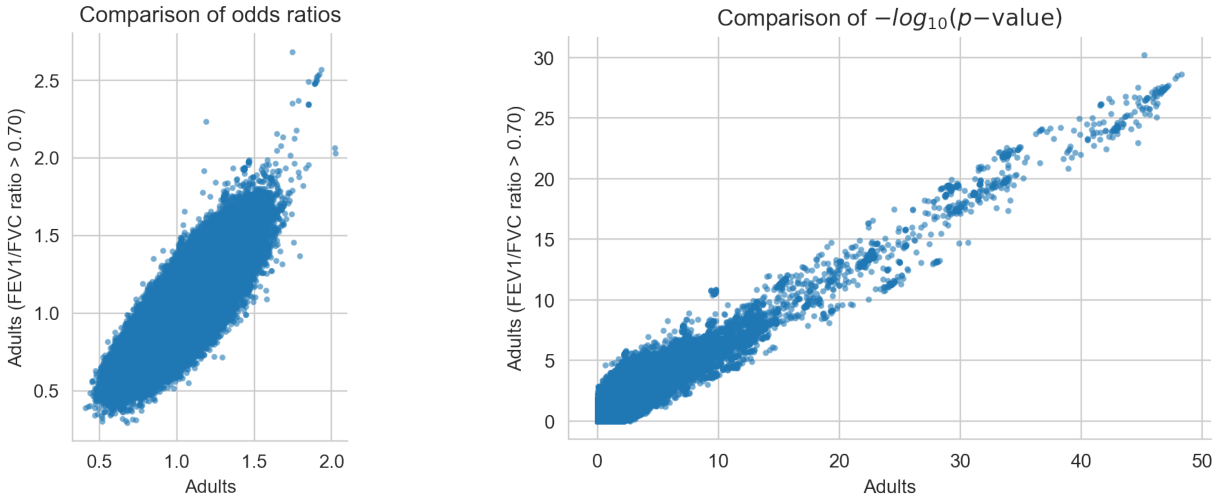

### Figures

**Figure 1. Comparisons of odds ratios and p-values from childhood onset (top) and adult onset (bottom) GWAS using the top 10 principal components (x-axis) and the top 14 principal components (y-axis).**

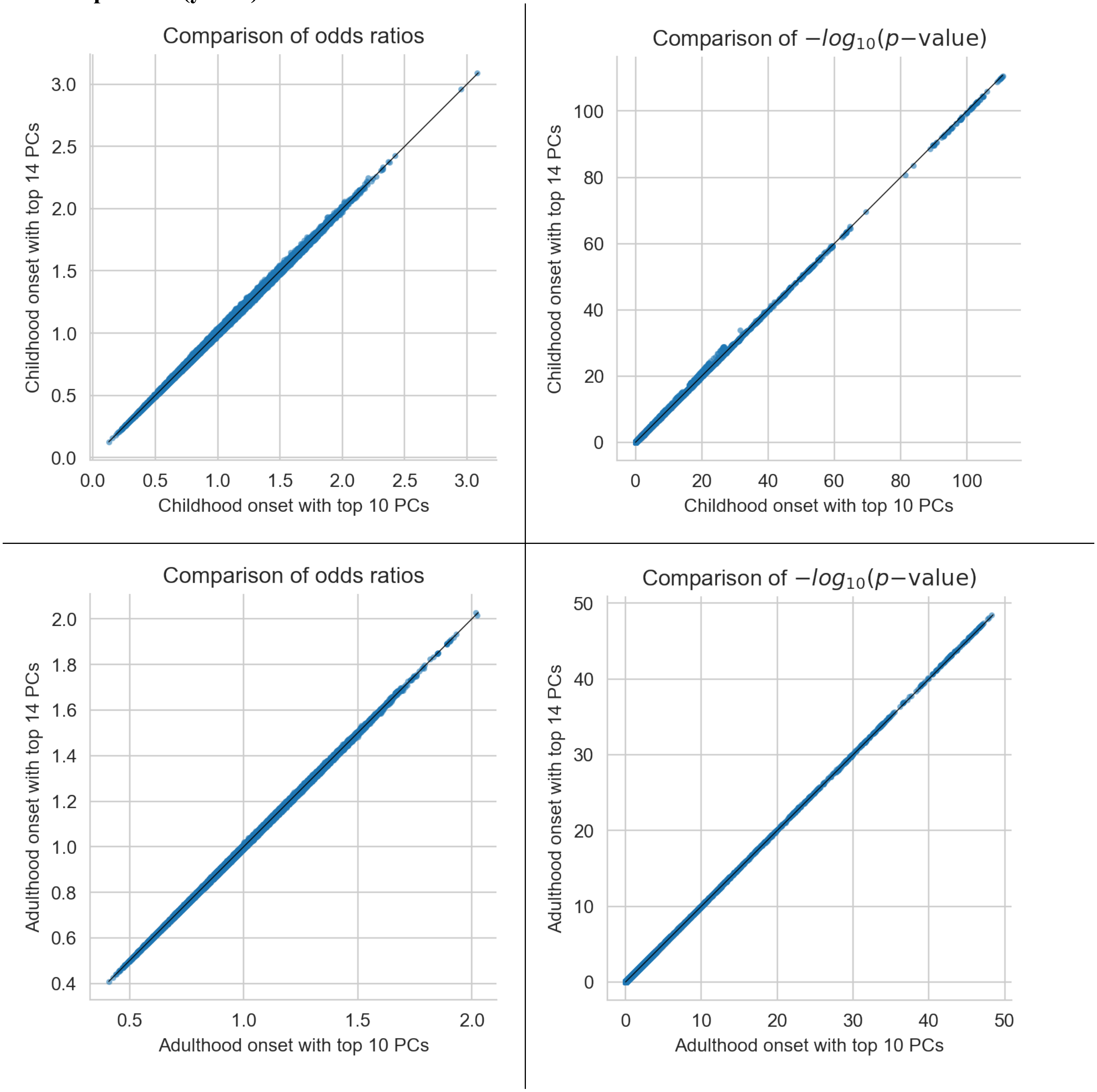

**Figure 2. Manhattan plots showing GWAS results excluding self-reported asthma.**

**A. The GWAS results of childhood onset (full set including self-report + ICD10, 9,433 cases) at the top and the childhood onset (ICD10 only, 3,462 cases) at the bottom.**

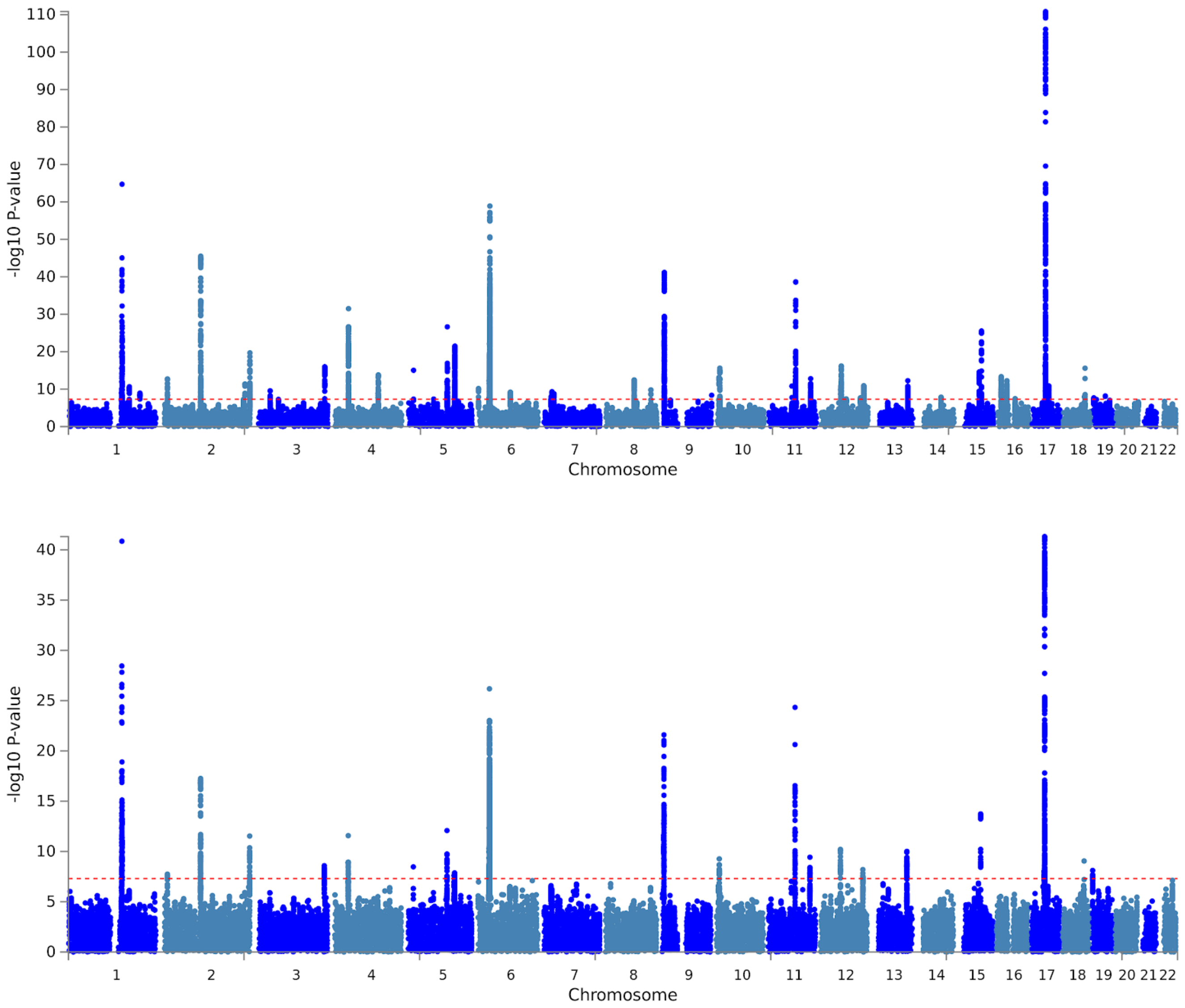

**B. The GWAS results of adult onset (full set including self-report + ICD10, 21,564 cases) at the top and the adult onset (ICD10 only, 9,260 cases) at the bottom.**

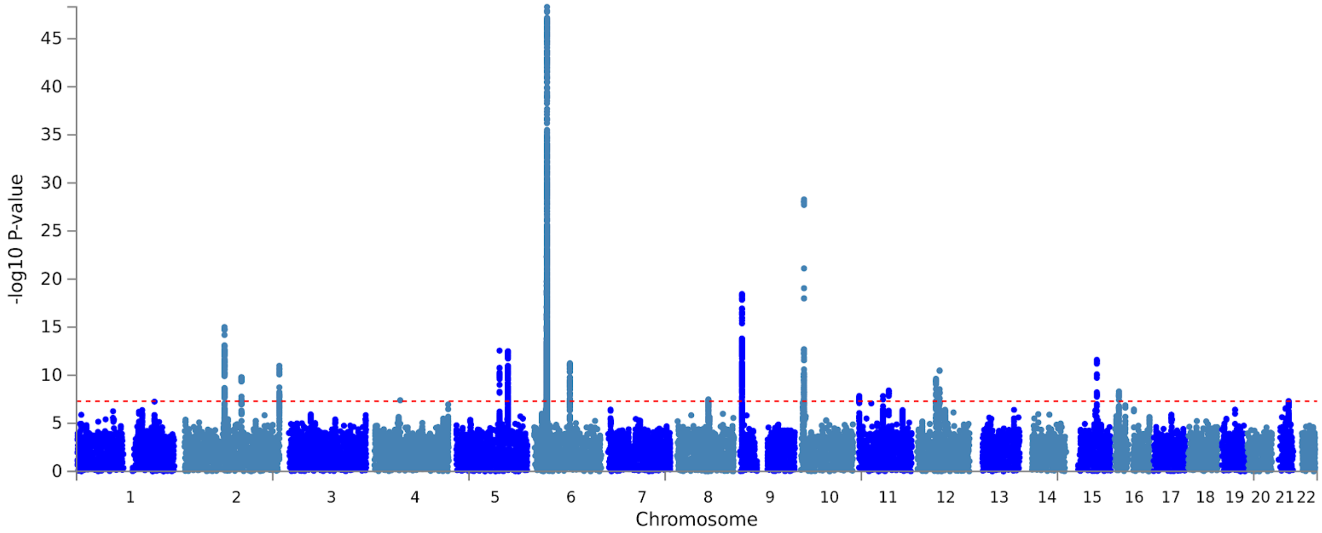

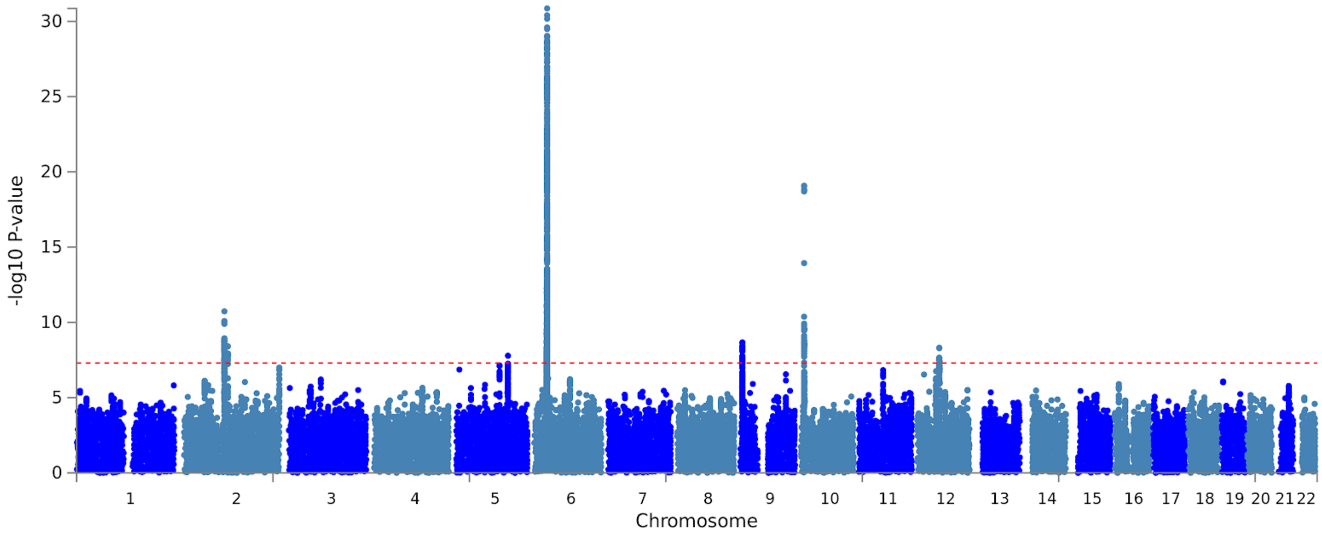

**Figure 3. QQ-plot of p-values from a GWAS comparing patients with asthma with ICD10 + self-report to those with self-report only.**

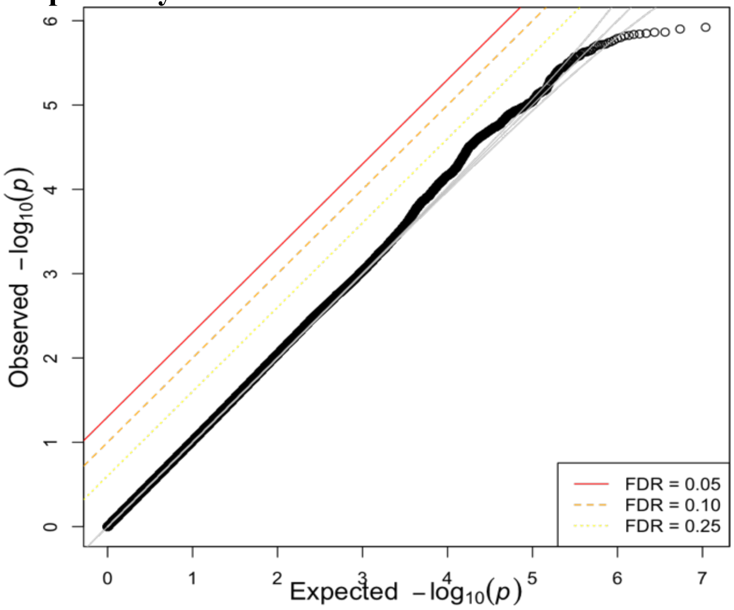

**Figure 4. Forest plot comparing ORs from the GWASs for all childhood onset cases and childhood onset cases excluding those with allergy.** ORs and 95% confidence intervals are shown for the 61 loci in Table 2 (blue, childhood onset asthma; green, childhood onset asthma excluding allergy), ordered by the OR in the childhood onset GWAS (largest to smallest).

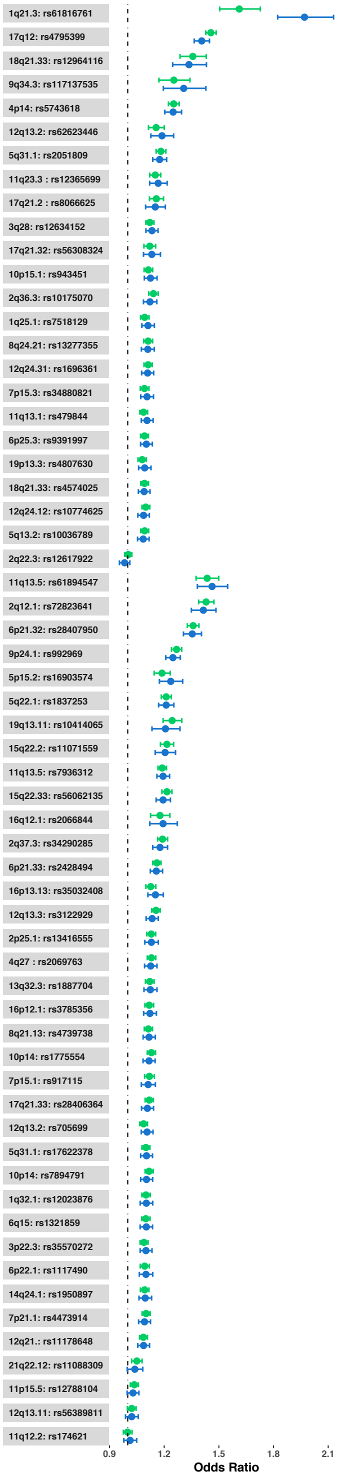

**Figure 5. Regional association plots for 23 childhood and one adult onset asthma specific loci.**

**Loci Associated with Childhood Onset Asthma Only**

**1q21·3 – rs61816761**

Childhood Onset

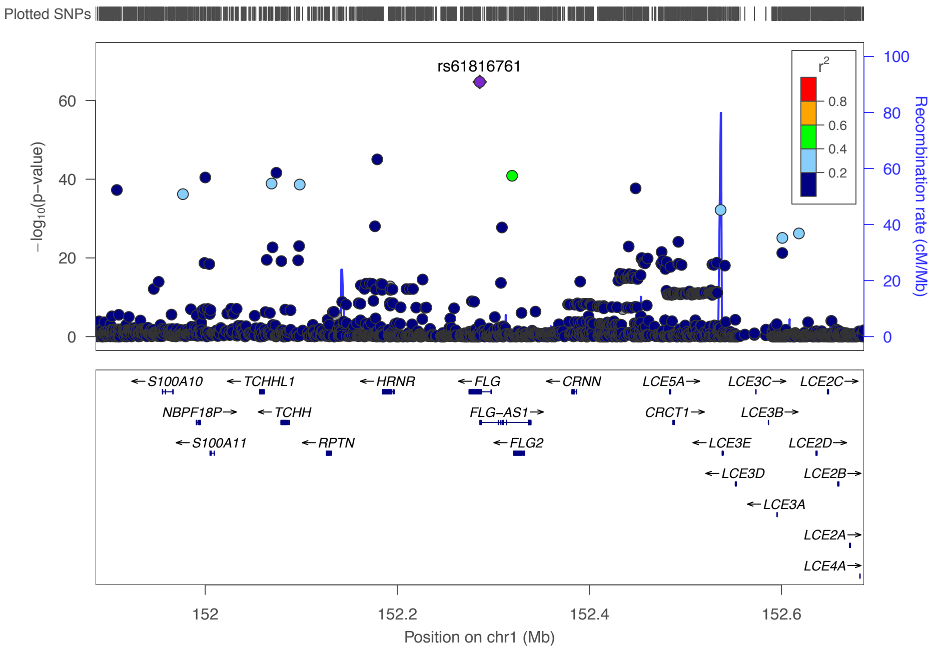

Adult Onset

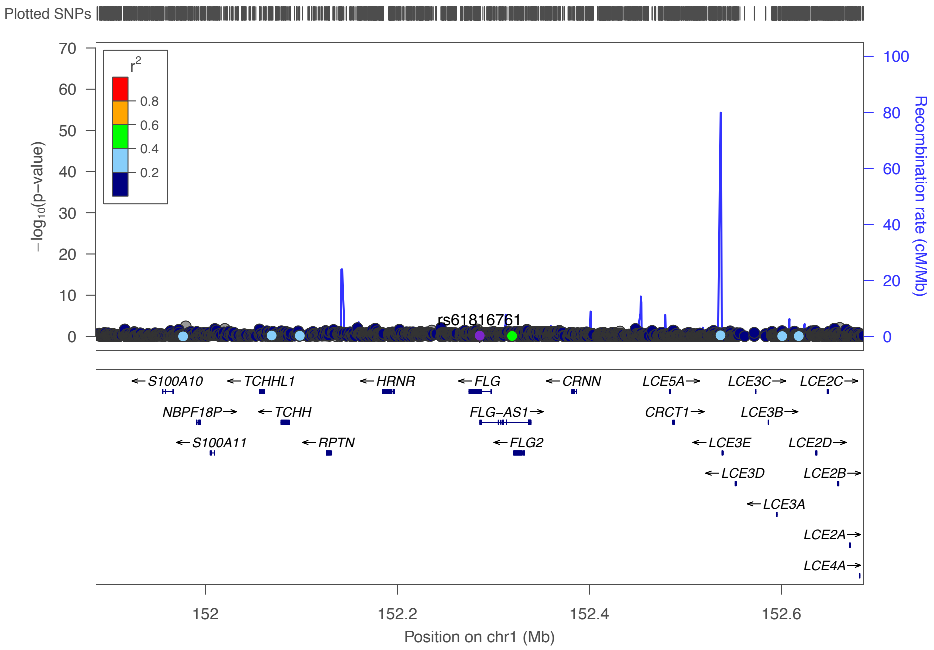

**1q21·5 – rs7518129**

Childhood Onset

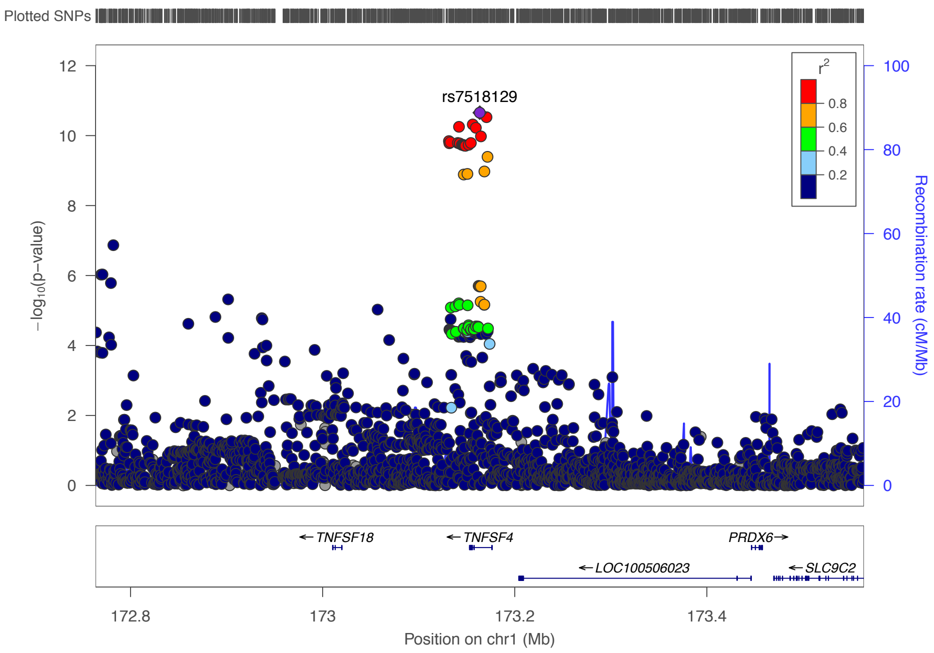

Adult Onset

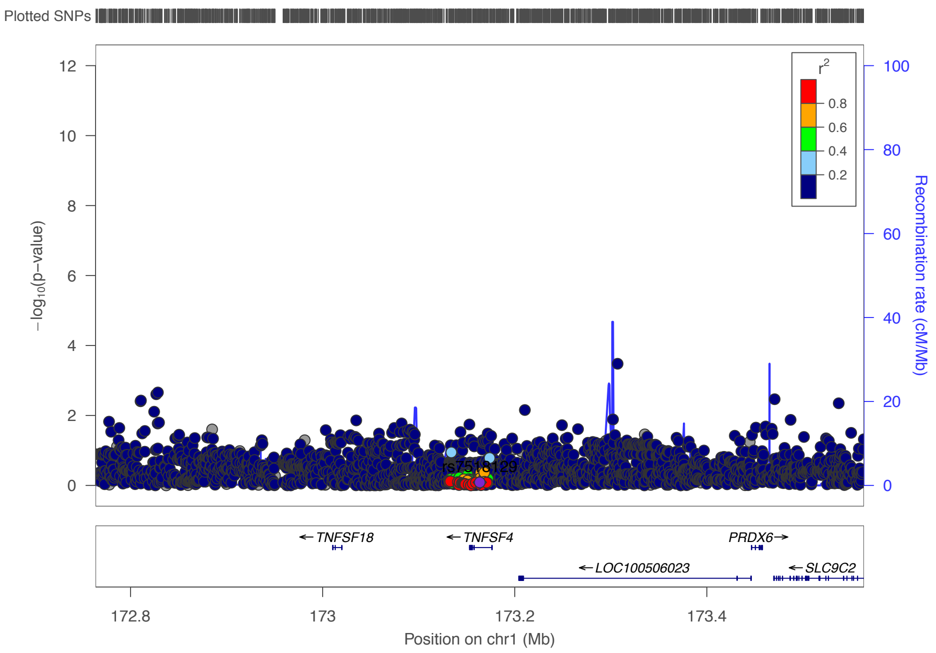

**2q36·3 – rs10175070**

Childhood Onset

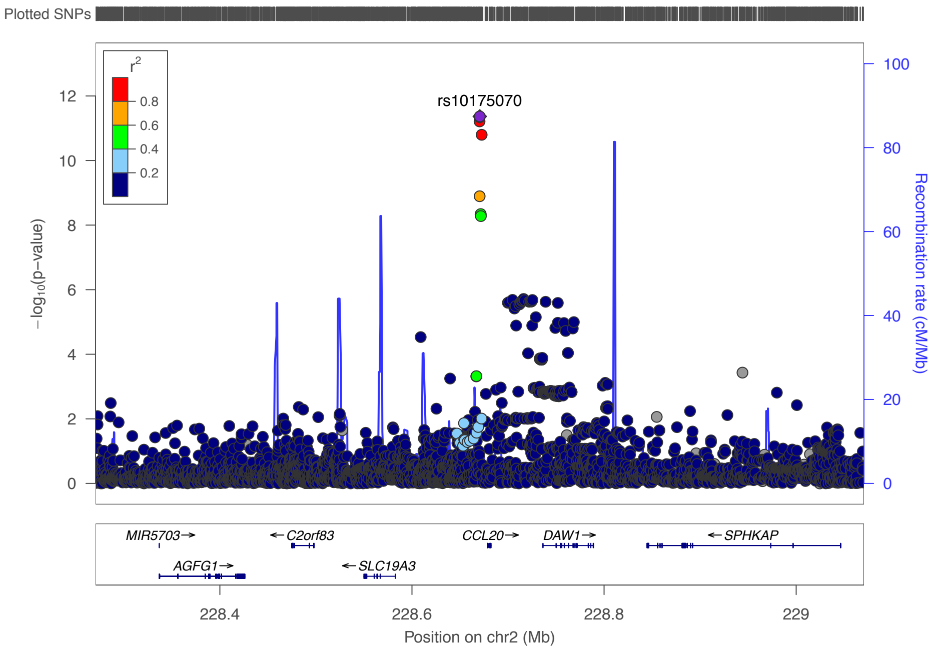

Adult Onset

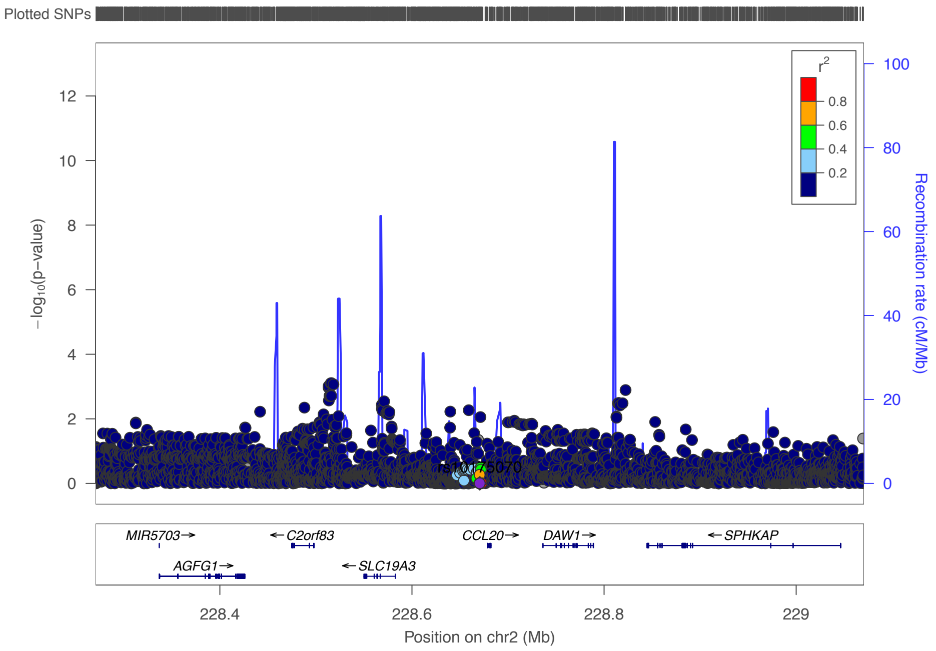

# 3q28 – rs12634152

Childhood Onset

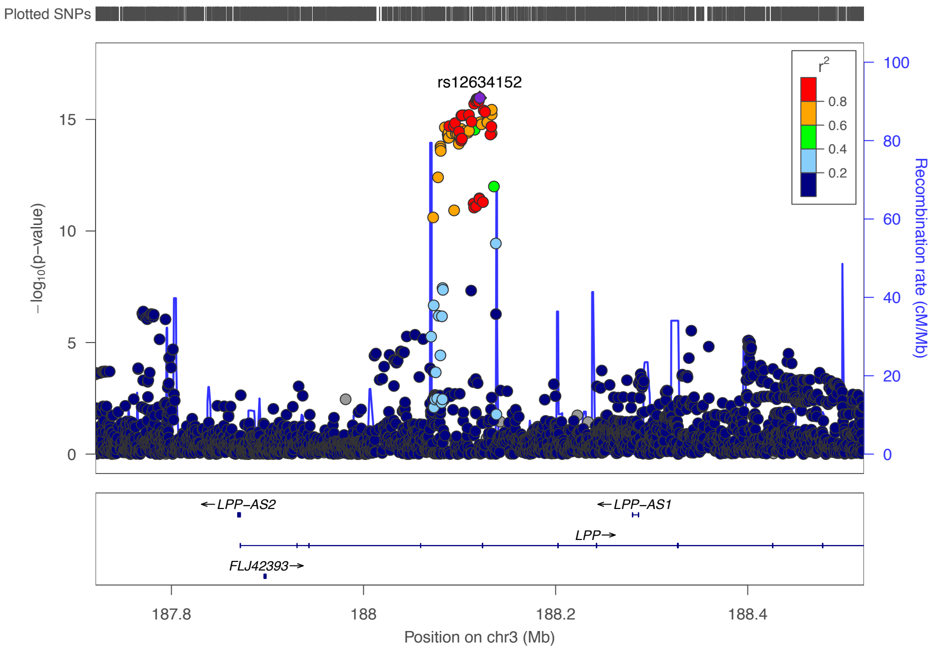

Adult Onset

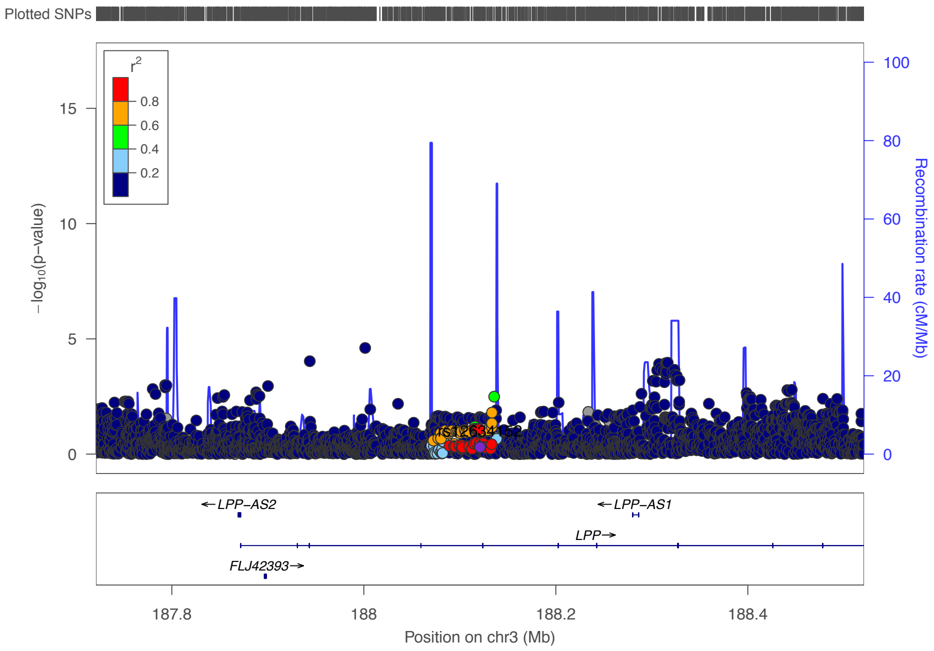

**4p14 – rs5743618**

Childhood Onset

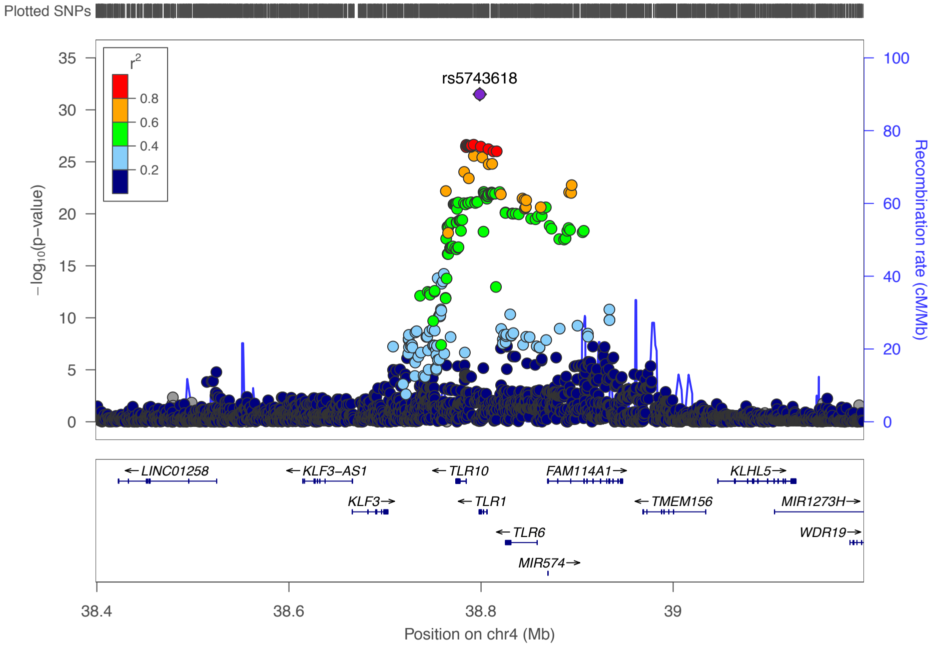

Adult Onset

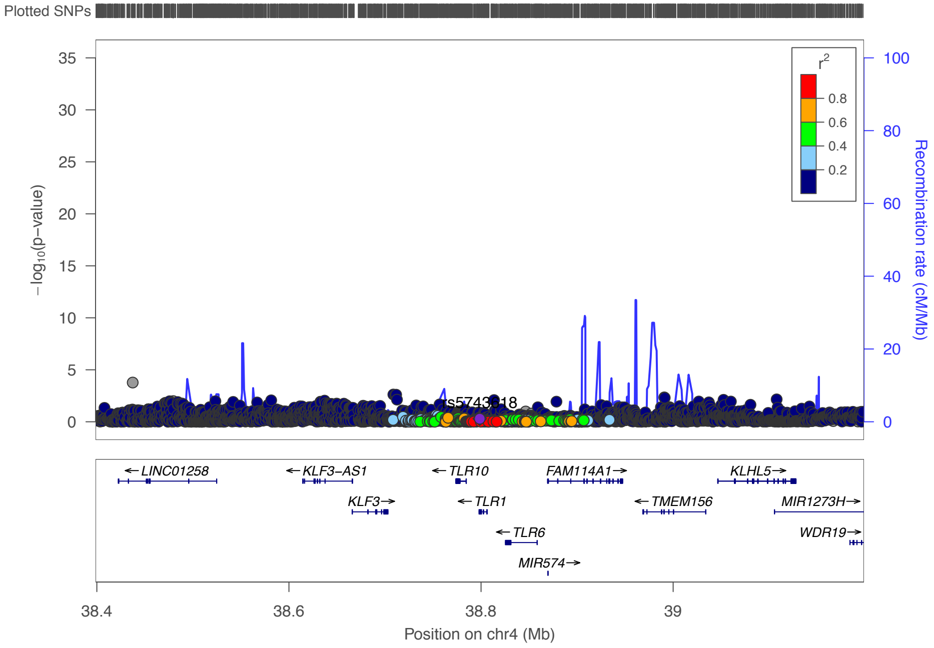

**5q31·2 – rs10036789**

Childhood Onset | Adult Onset

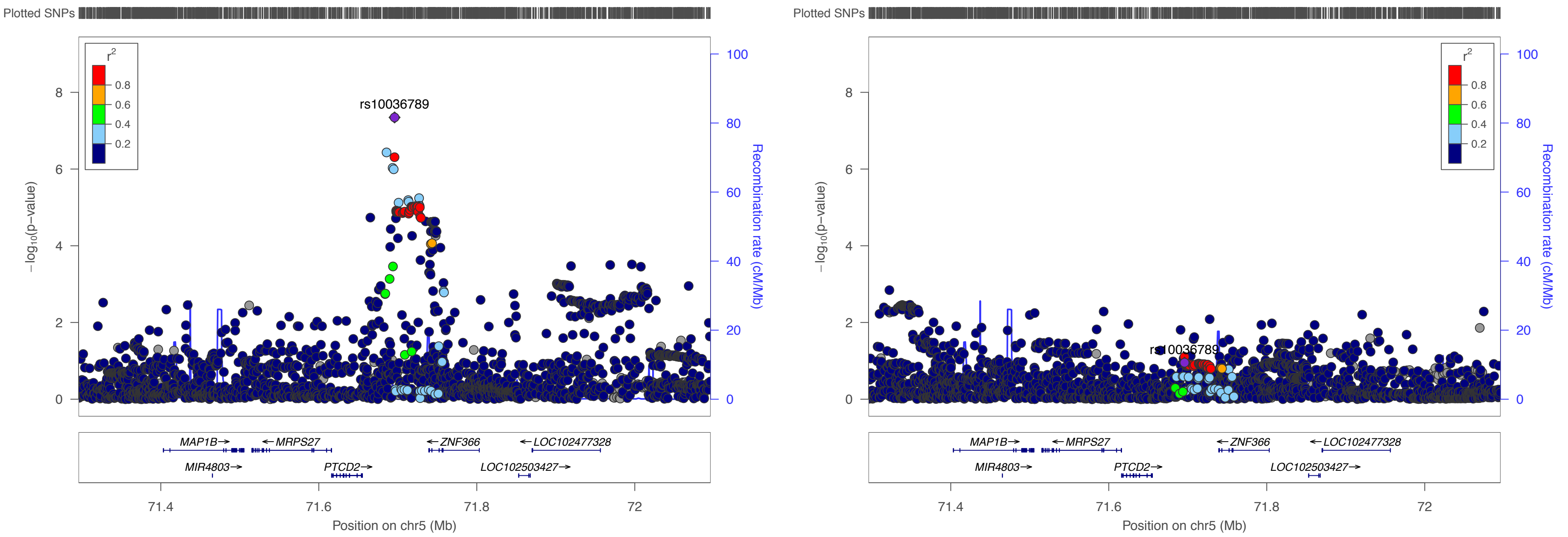

**5q31·1 – rs2051809**

Childhood Onset

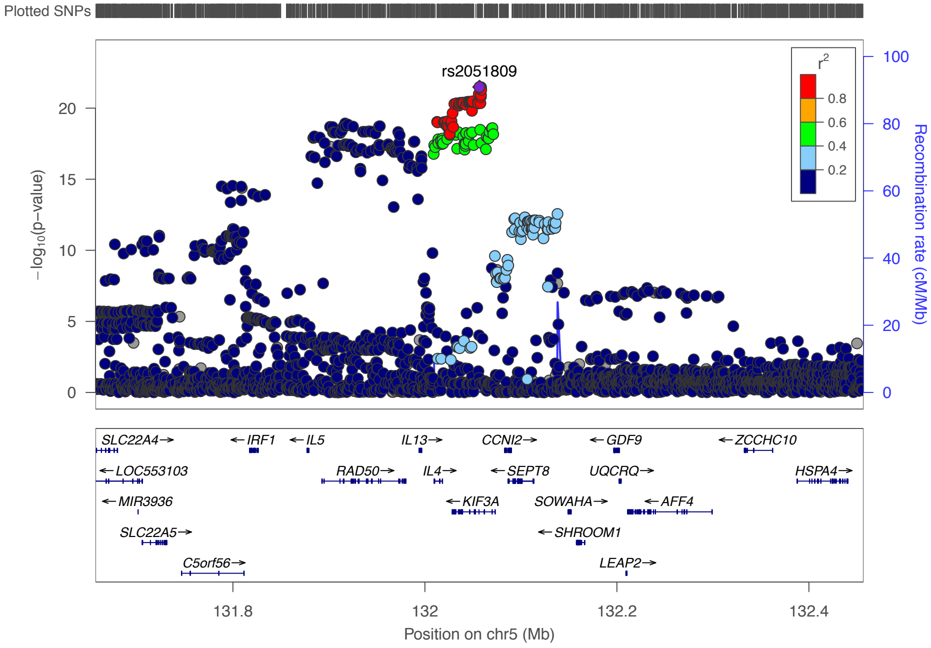

Adult Onset

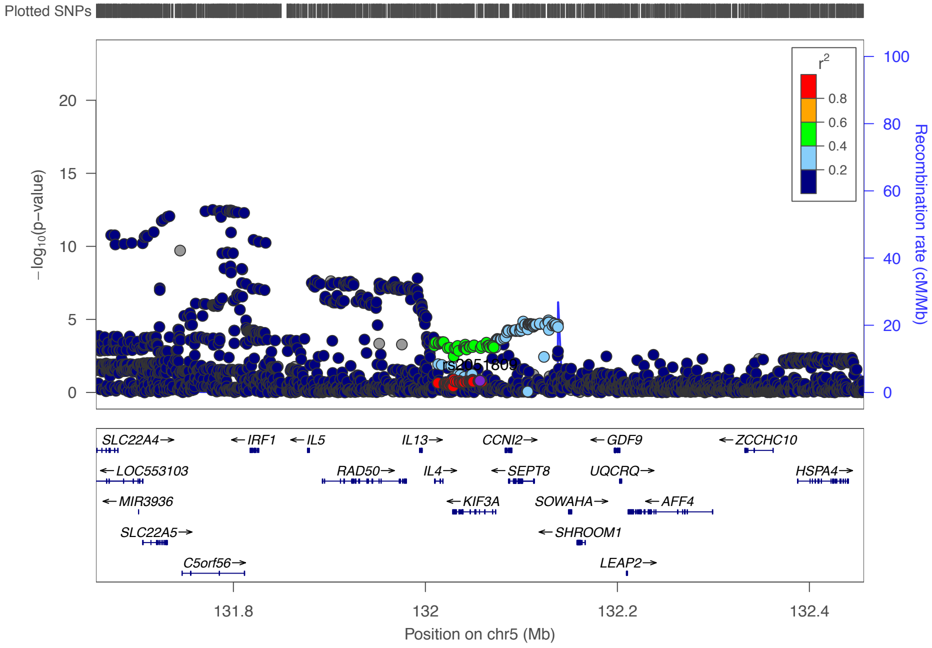

**6p25·3 – rs9391997**

Childhood Onset

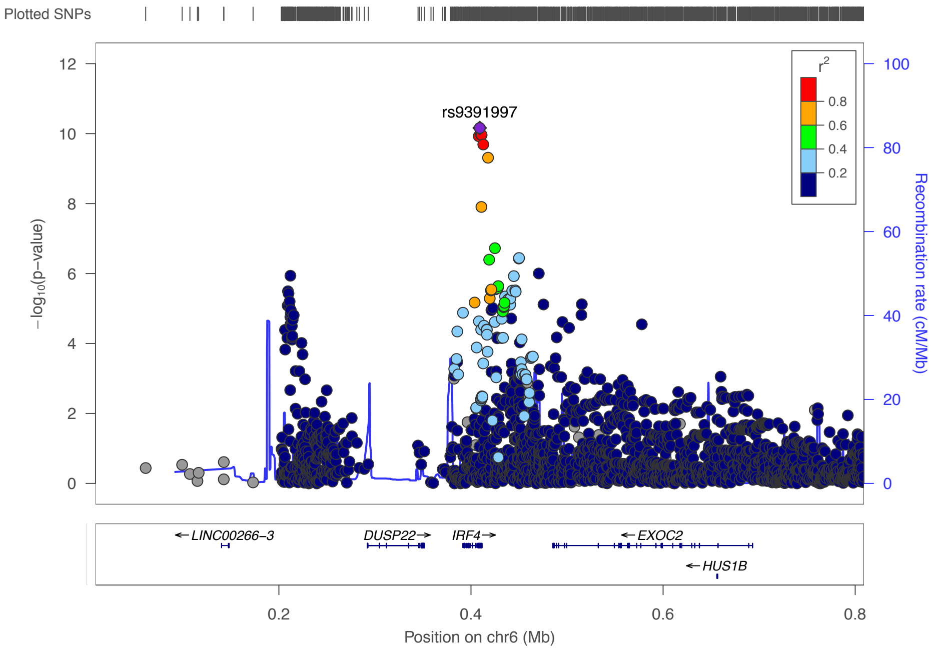

Adult Onset

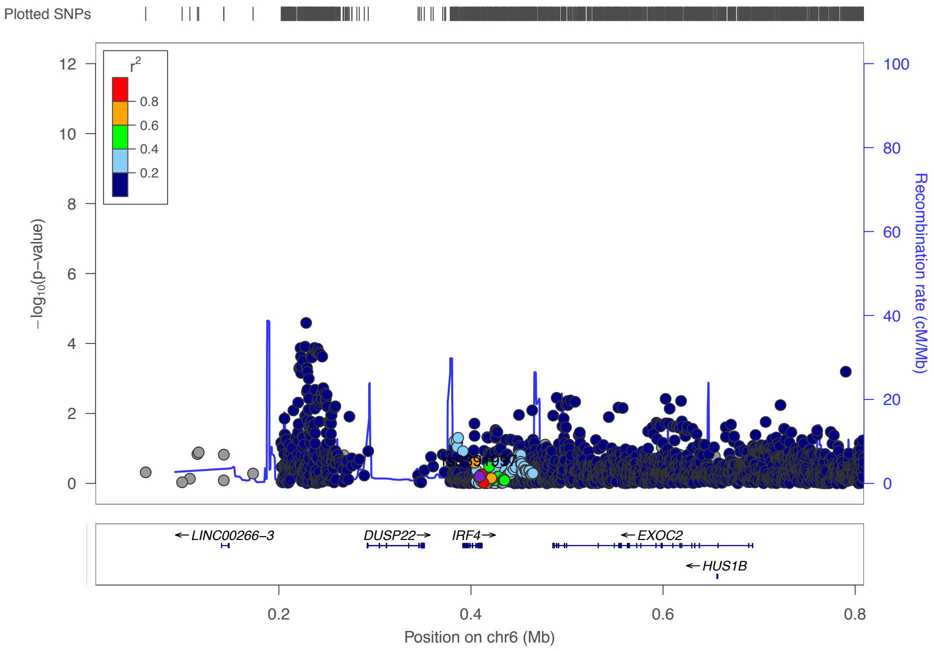

**7p15·3 – rs34880821**

Childhood Onset

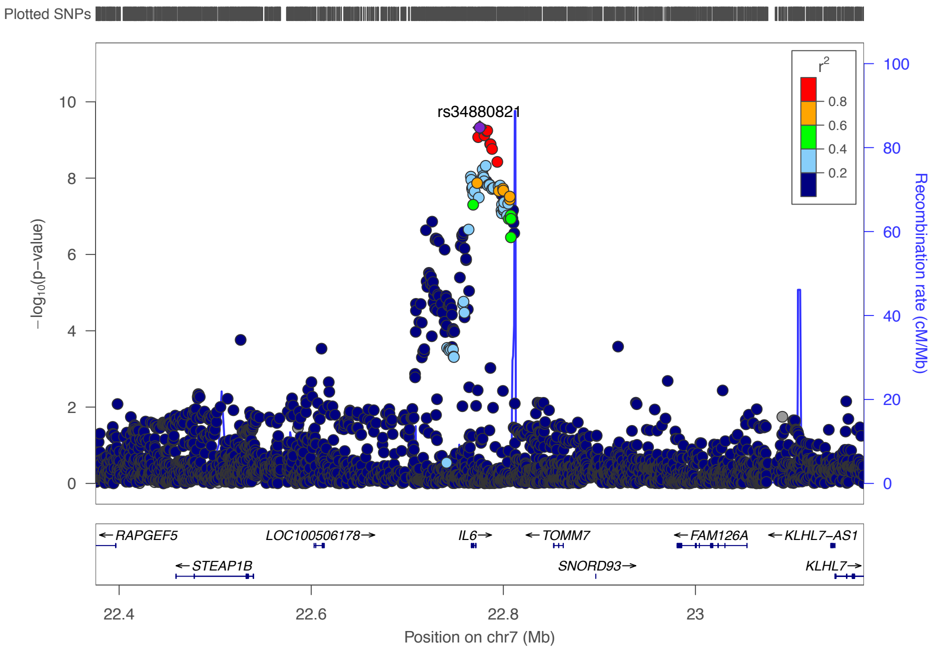

Adult Onset

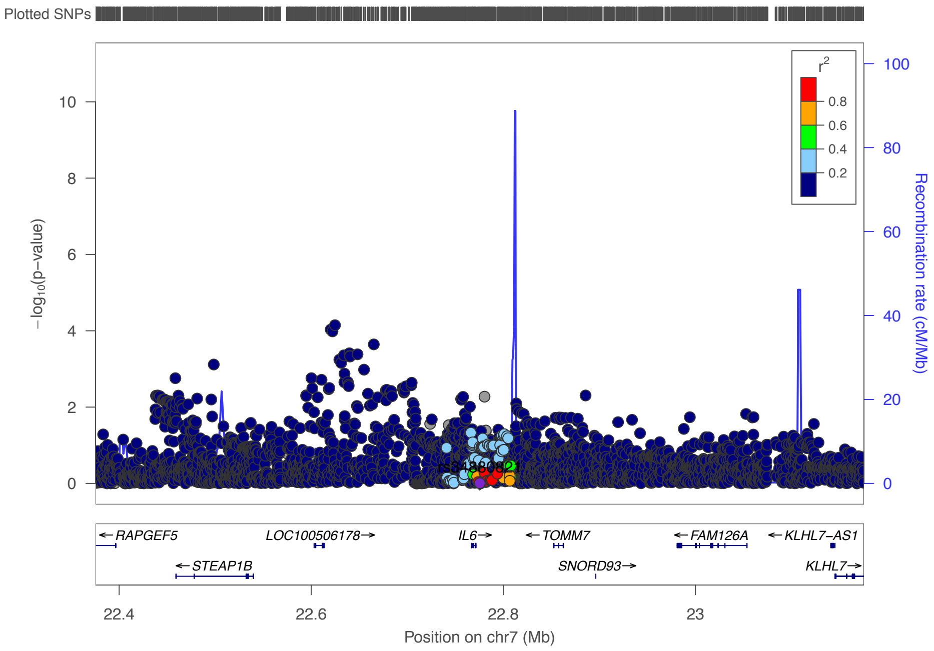

## 8q24·21 – rs13277355

##### Childhood Onset

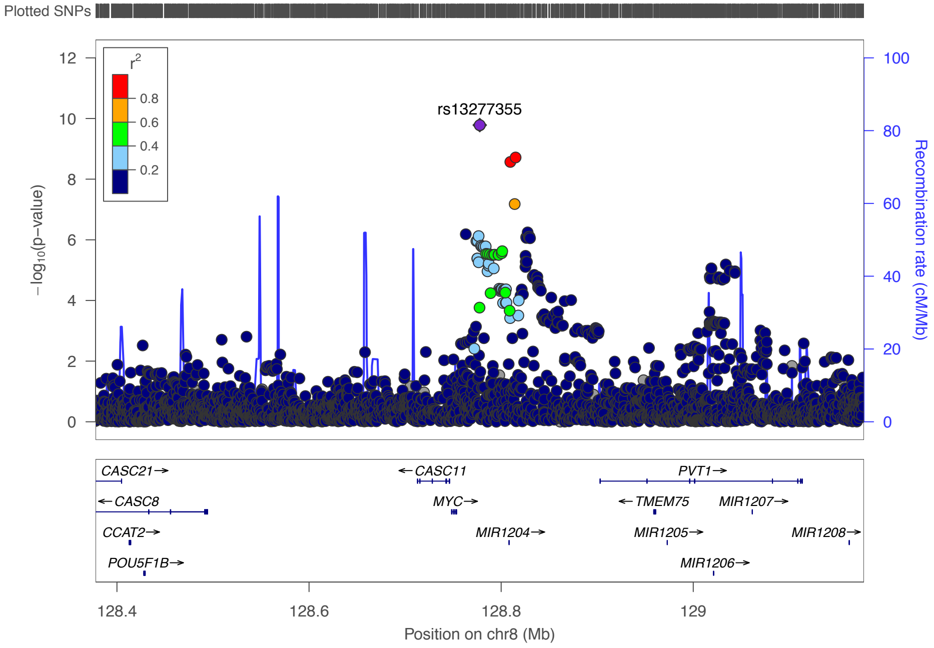

##### Adult Onset

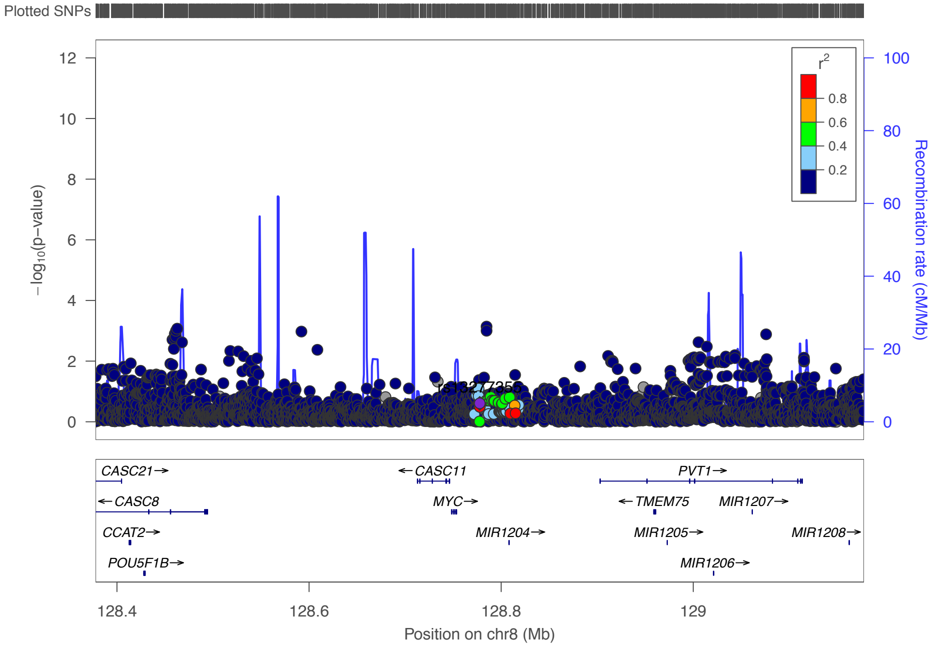

### 9q34·3 – rs117137535

Childhood Onset

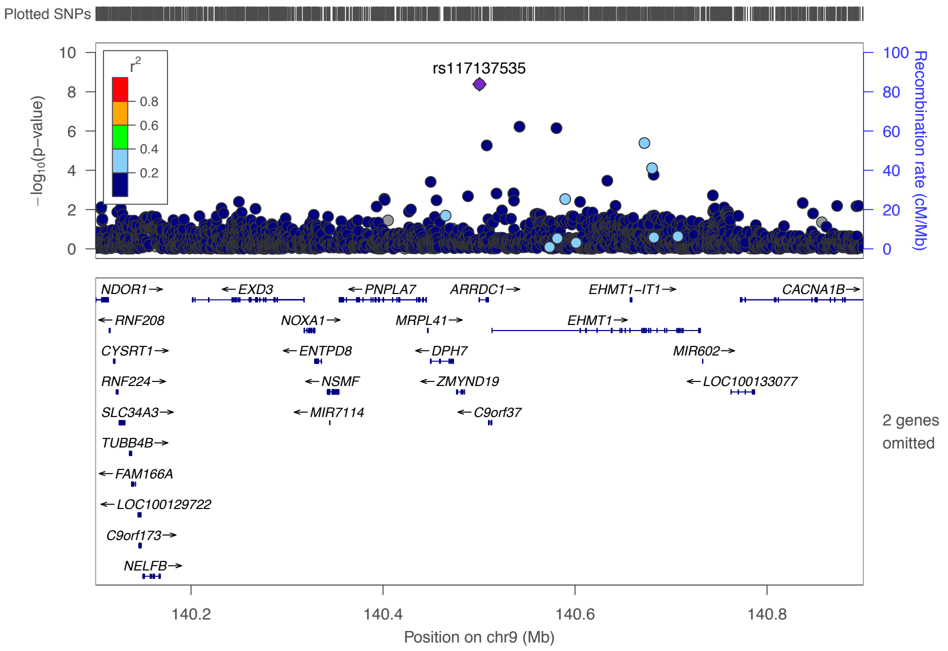

Adult Onset

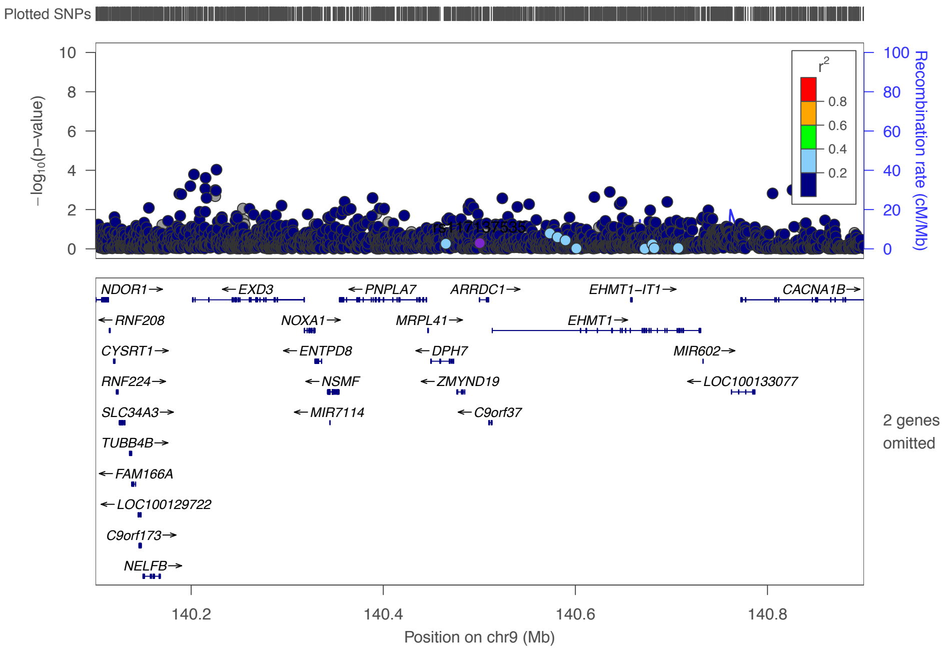

**10p15·1 – rs943451**

Childhood Onset

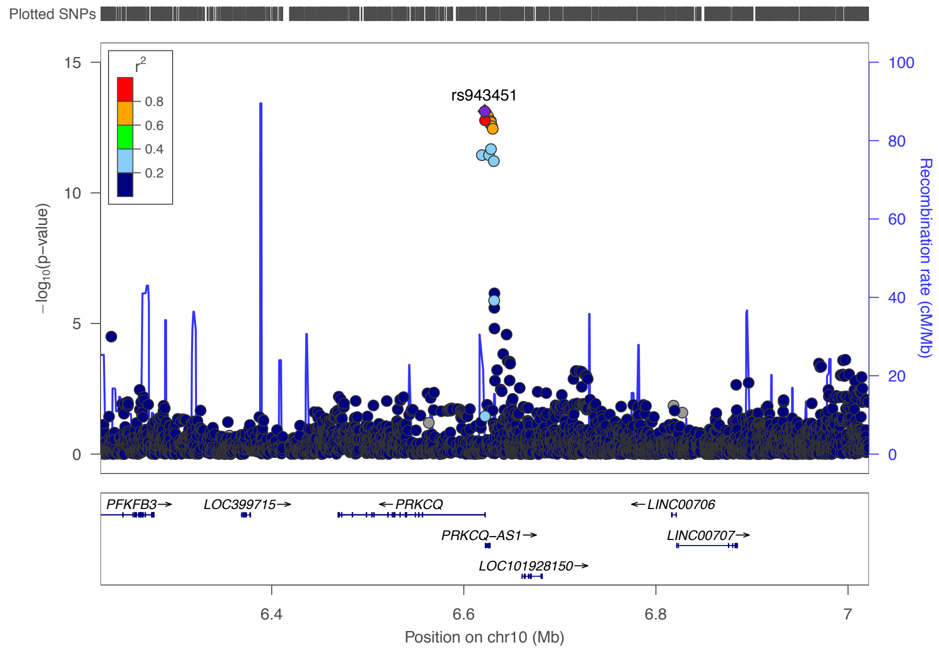

Adult Onset

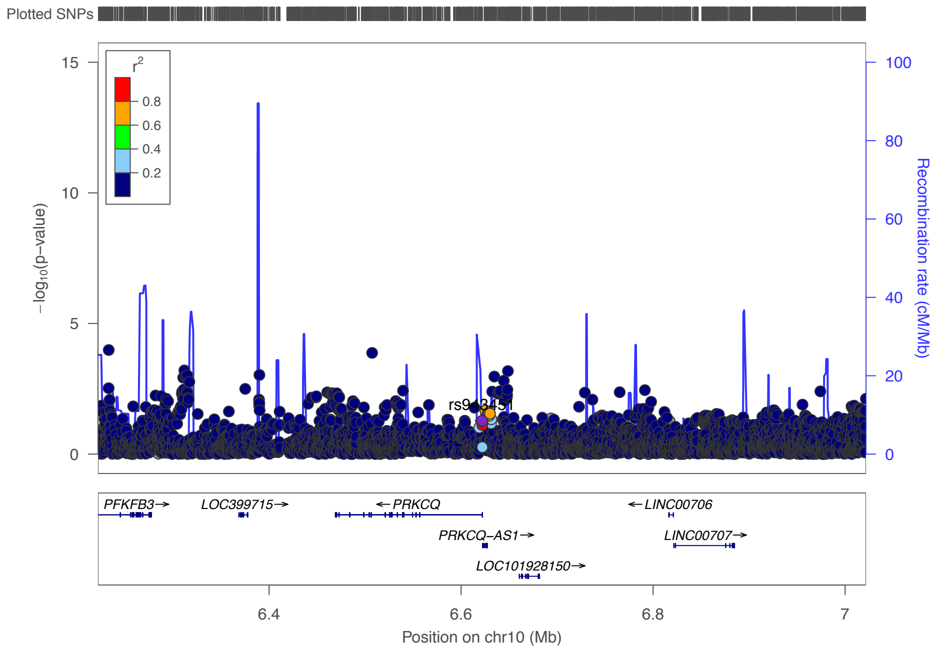

**11q23·3 – rs12365699**

Childhood Onset

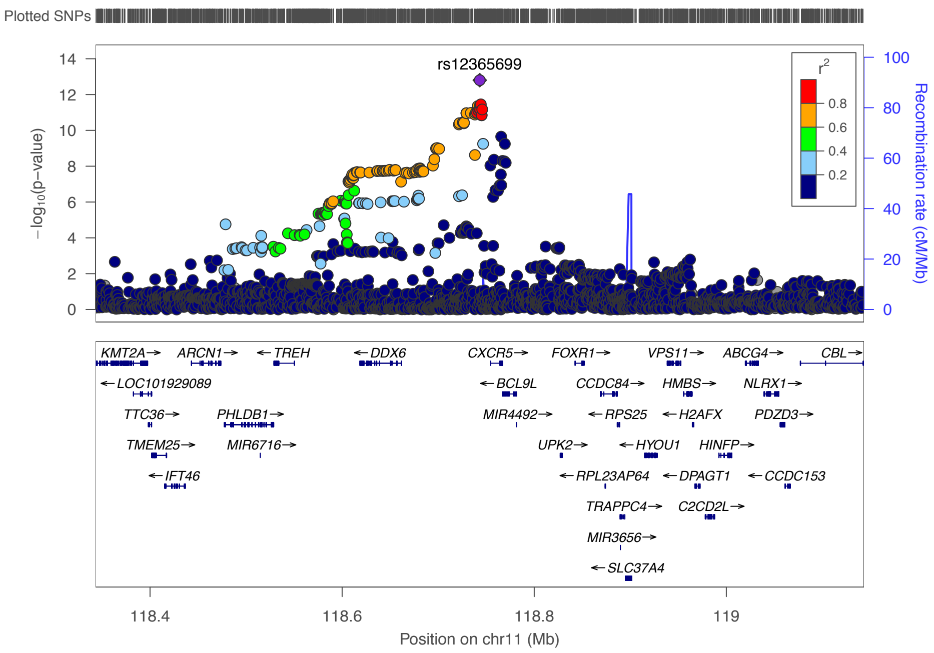

Adult Onset

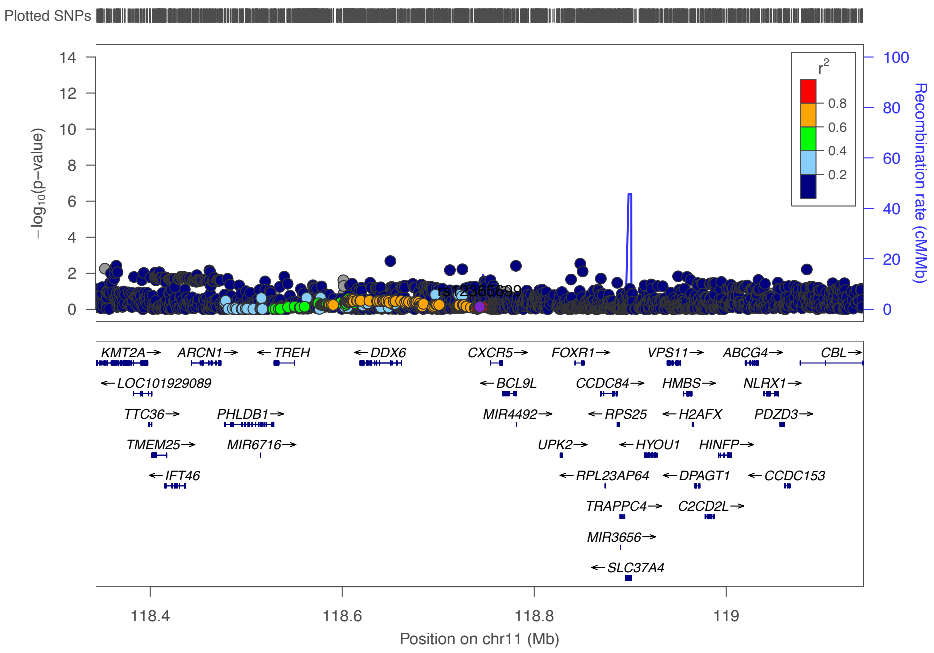

**12q13·2 – rs62623446**

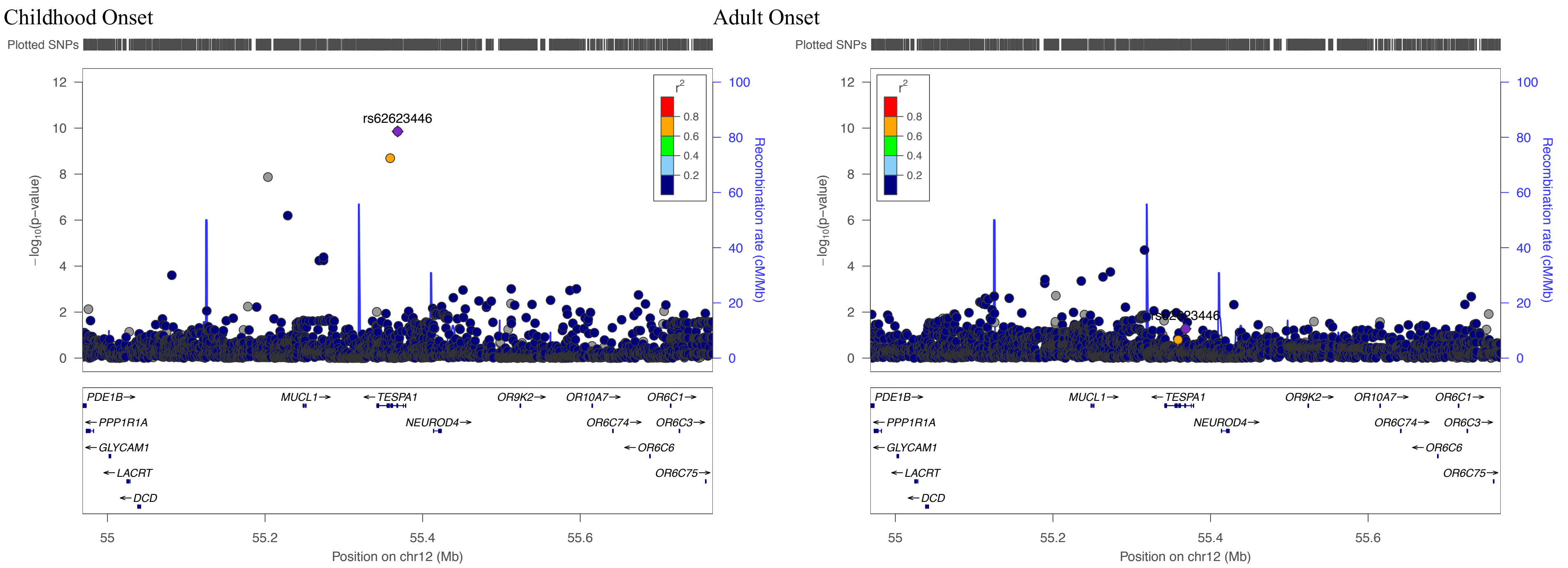

**12q24·31 – rs1696361**

Childhood Onset

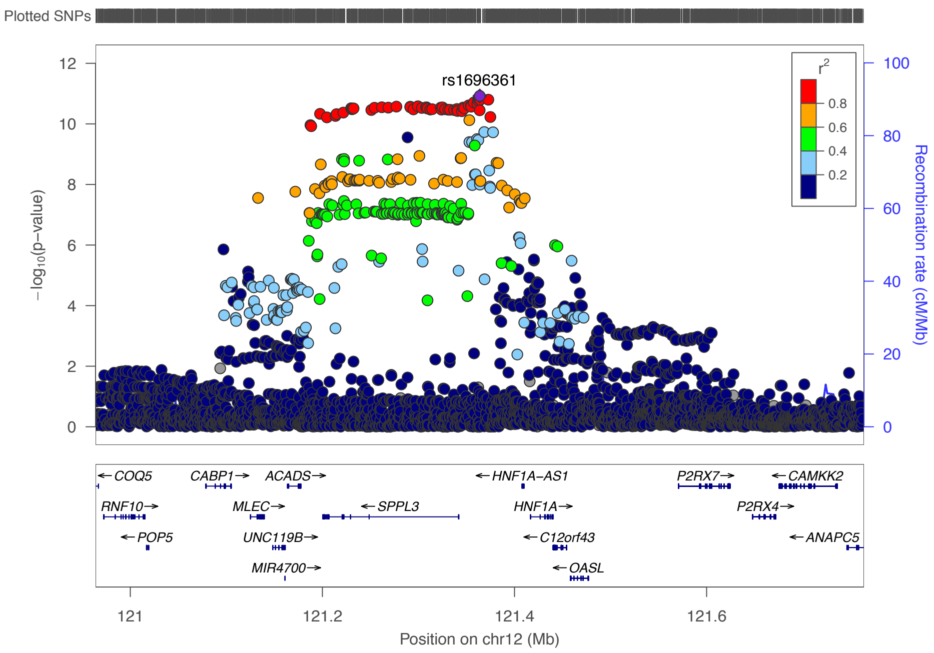

Adult Onset

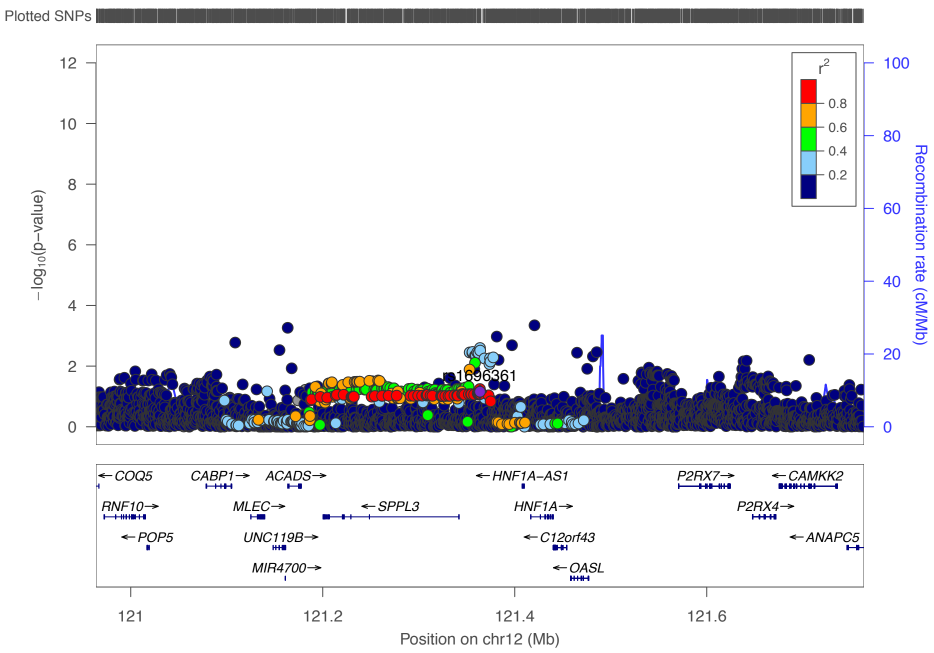

**17q12 – rs4795399**

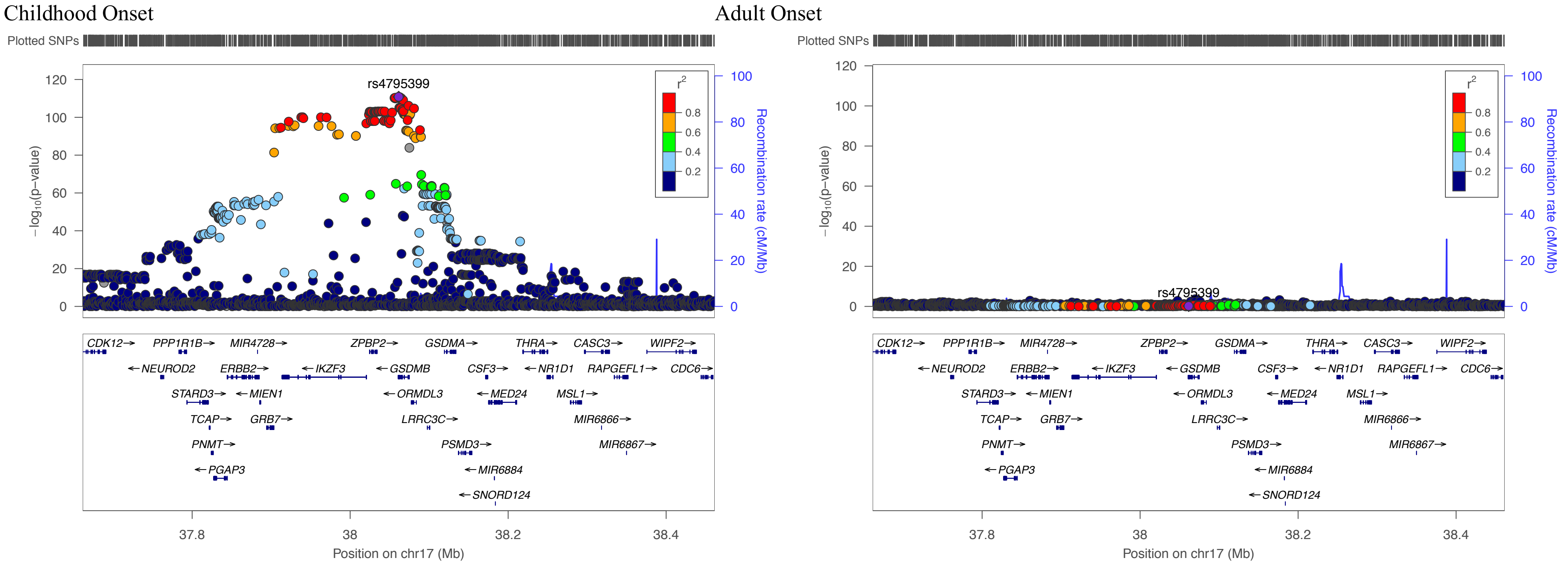

**17q21·2 – rs8066625**

Childhood Onset

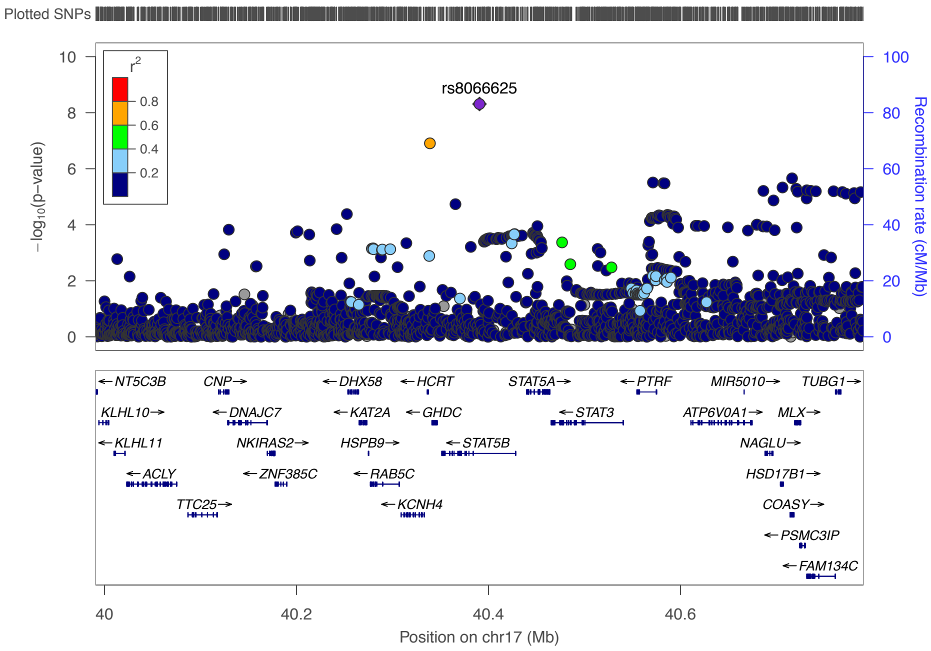

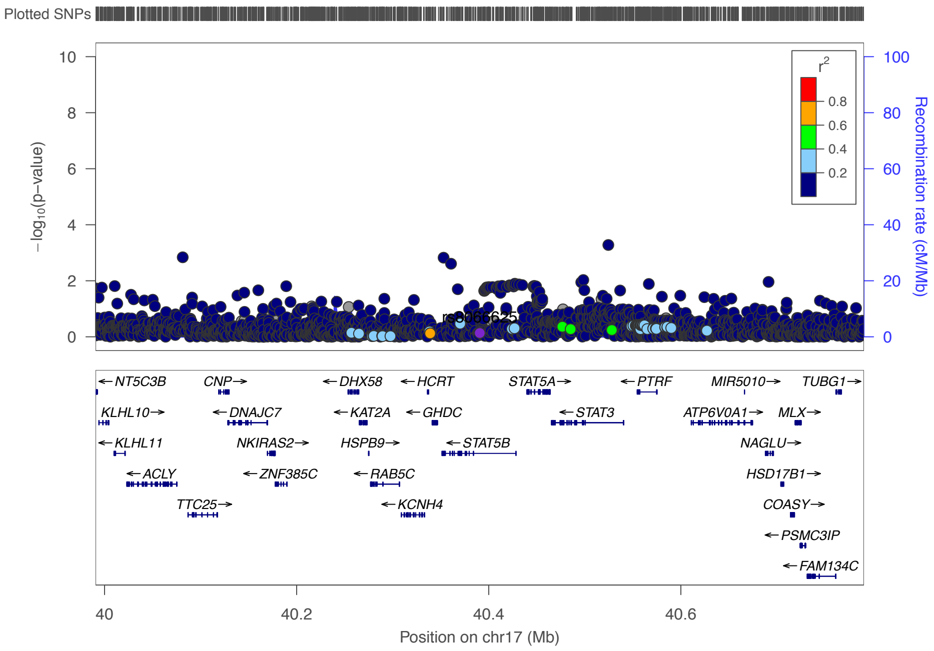

**17q21·32 – rs56308324**

Childhood Onset

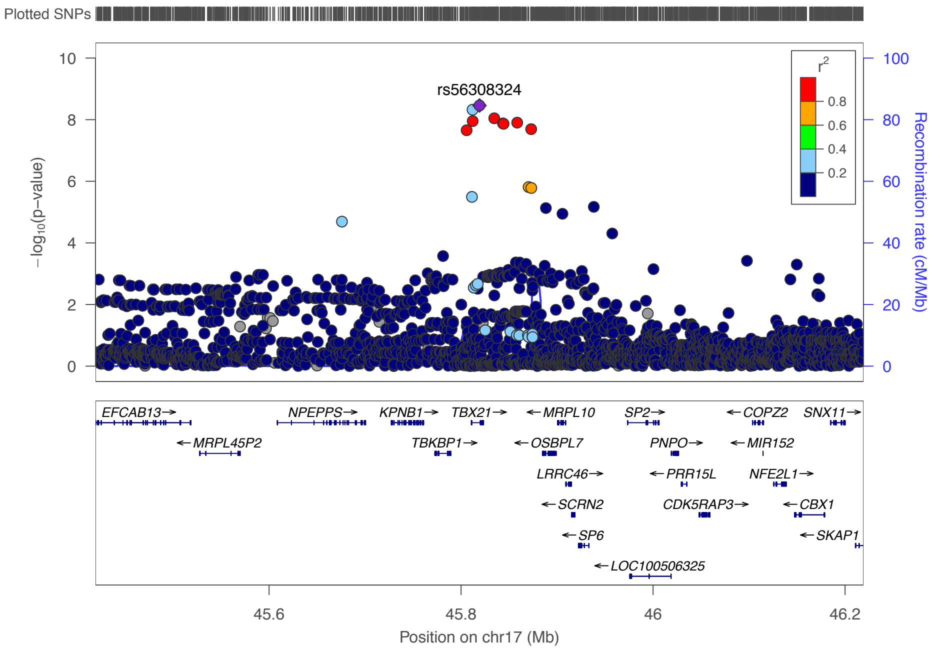

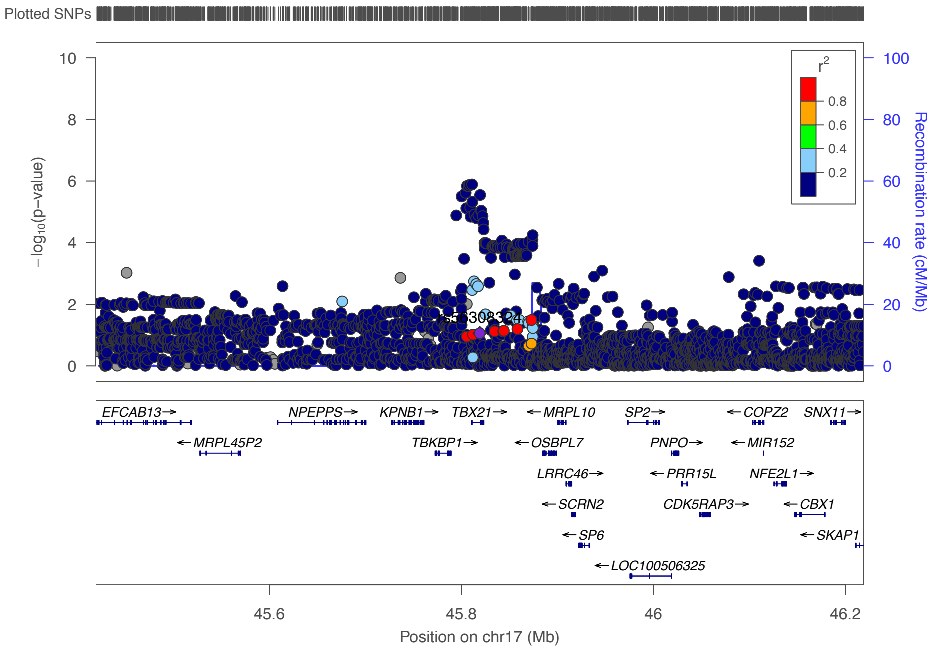

**18q21·33 – rs4574025**

Childhood Onset

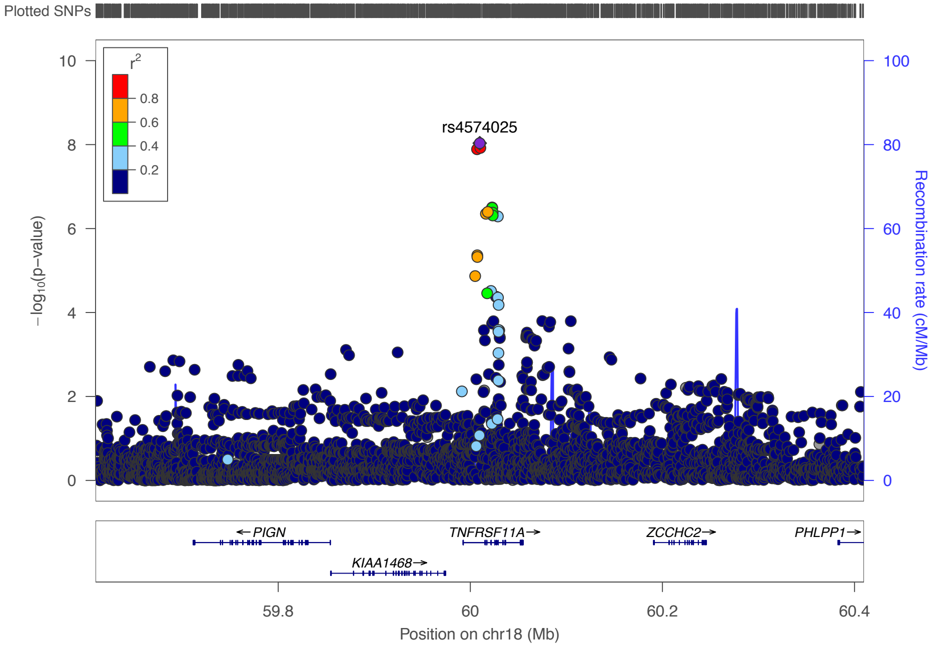

Adult Onset

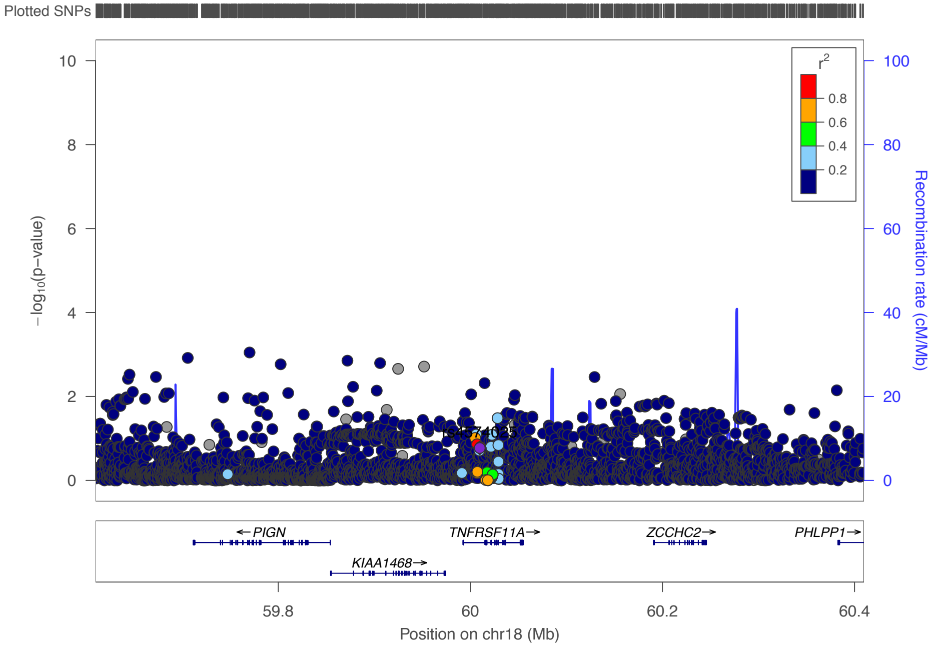

**18q21·33 – rs12964116**

Childhood Onset

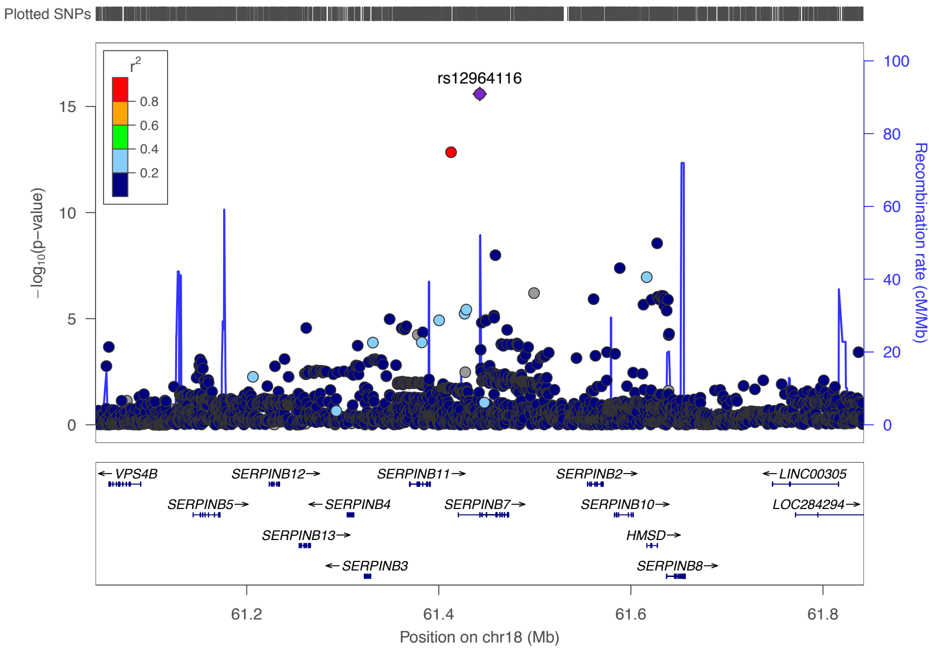

Adult Onset

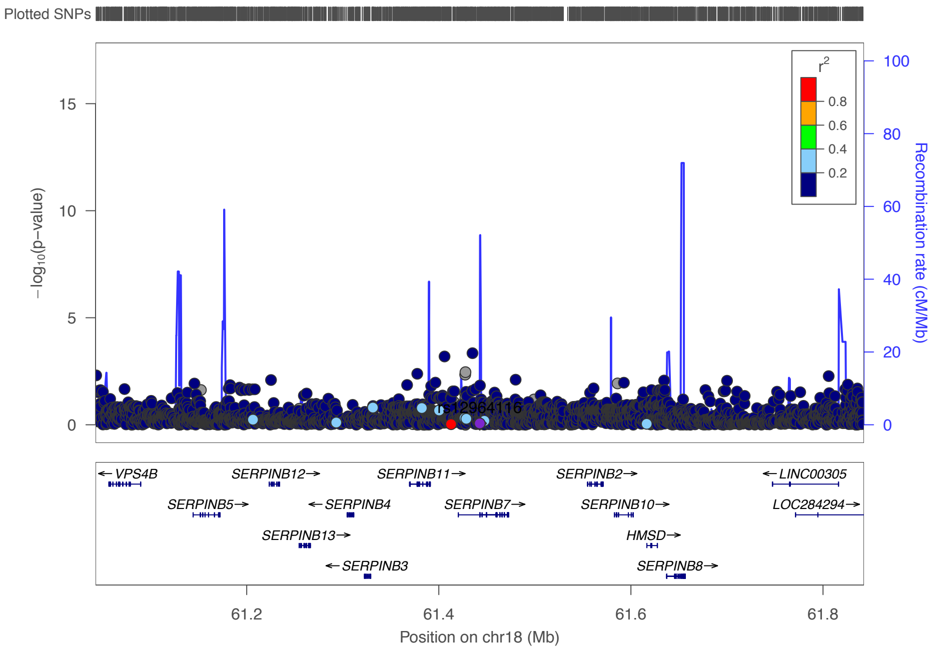

**19p13·3 – rs4807630**

Childhood Onset

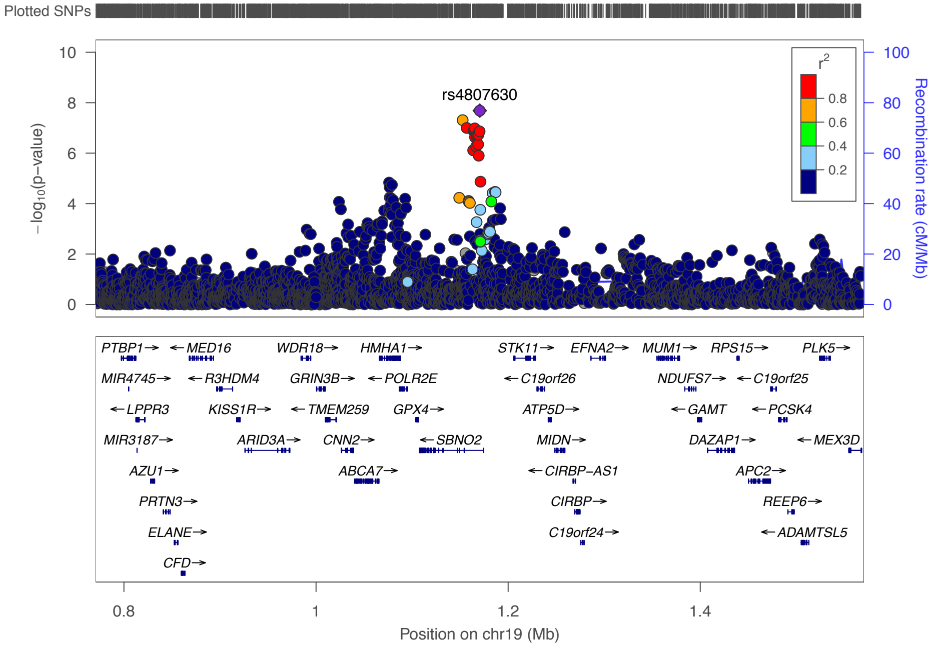

Adult Onset

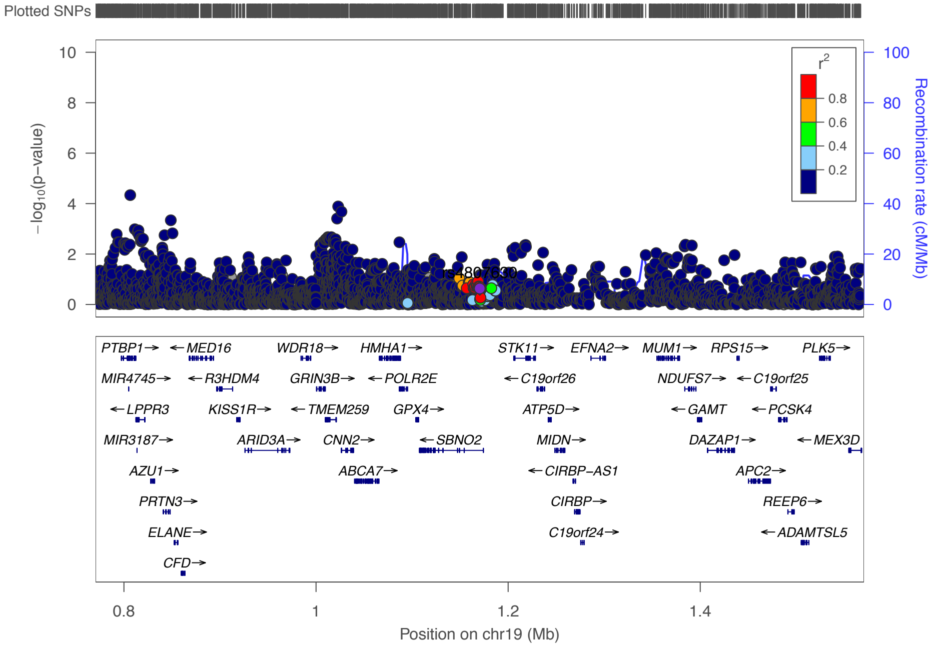

**Locus Associated with Adult Onset Asthma Only**

2q22·3 – rs12617922

Childhood Onset

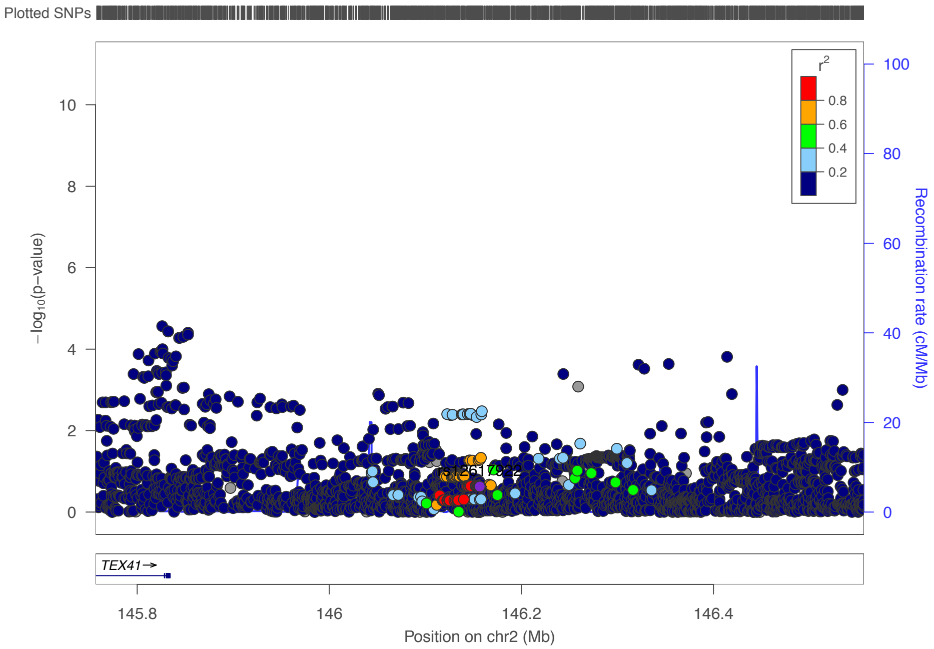

Adult Onset

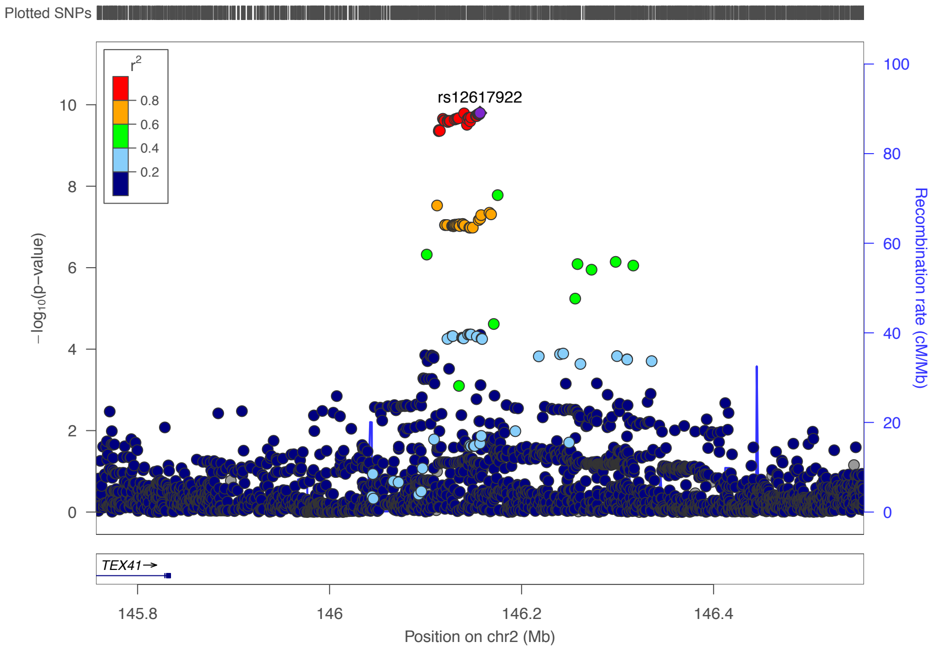

**Figure 6**. **Manhattan plot showing results for the asthma age of onset GWAS.** Each point corresponds to a SNP; the y-axis shows the $-\log_{10}$ p-value; the x-axis shows the position of each SNP along the 22 autosomes. The dashed red line shows genome-wide significance ($p=5x10^{-8}$).

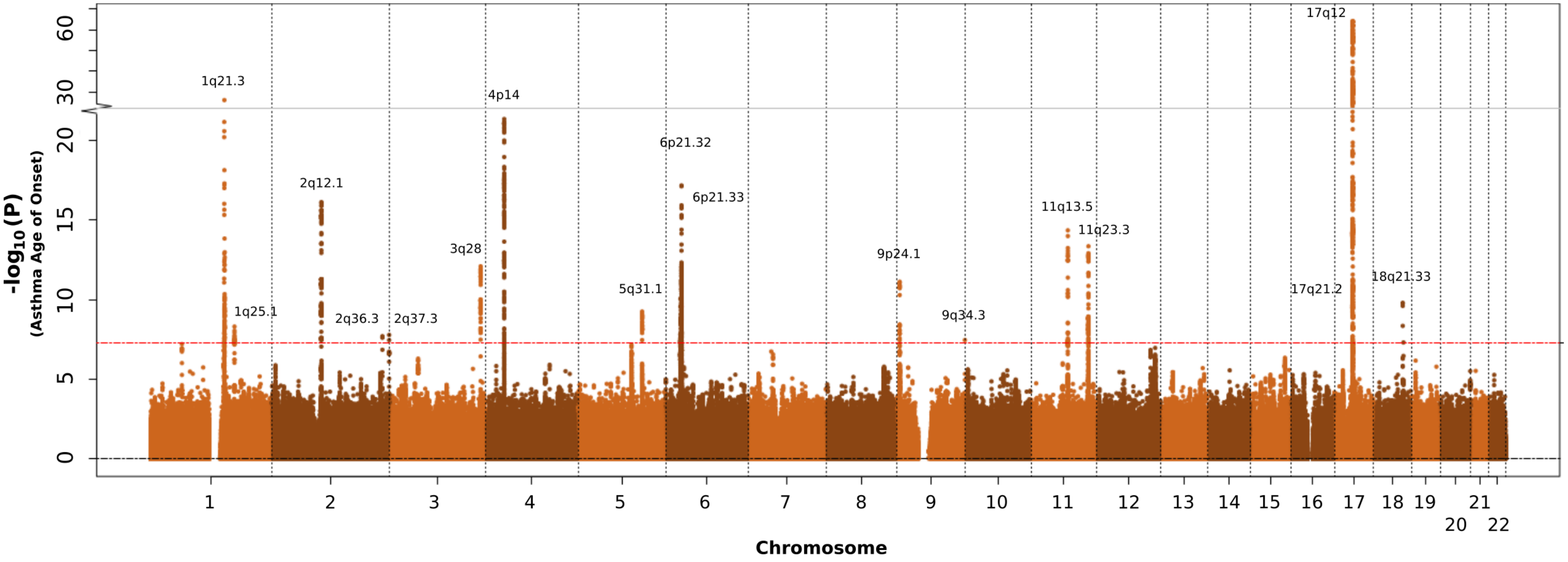

**Figure 7. Tissue-specific enrichment of genes mapped to GWAS significant loci (from FUMA[5]).** Asthma associated locus enrichment for differentially expressed genes in specific tissues (only more highly expressed genes are shown). A, childhood onset enrichment (at 54 loci); B, adult onset enrichment (19 loci). Tissues with $P_{Bon} < 0{\cdot}05$ are shown in red. See Table 4 below for p-values.

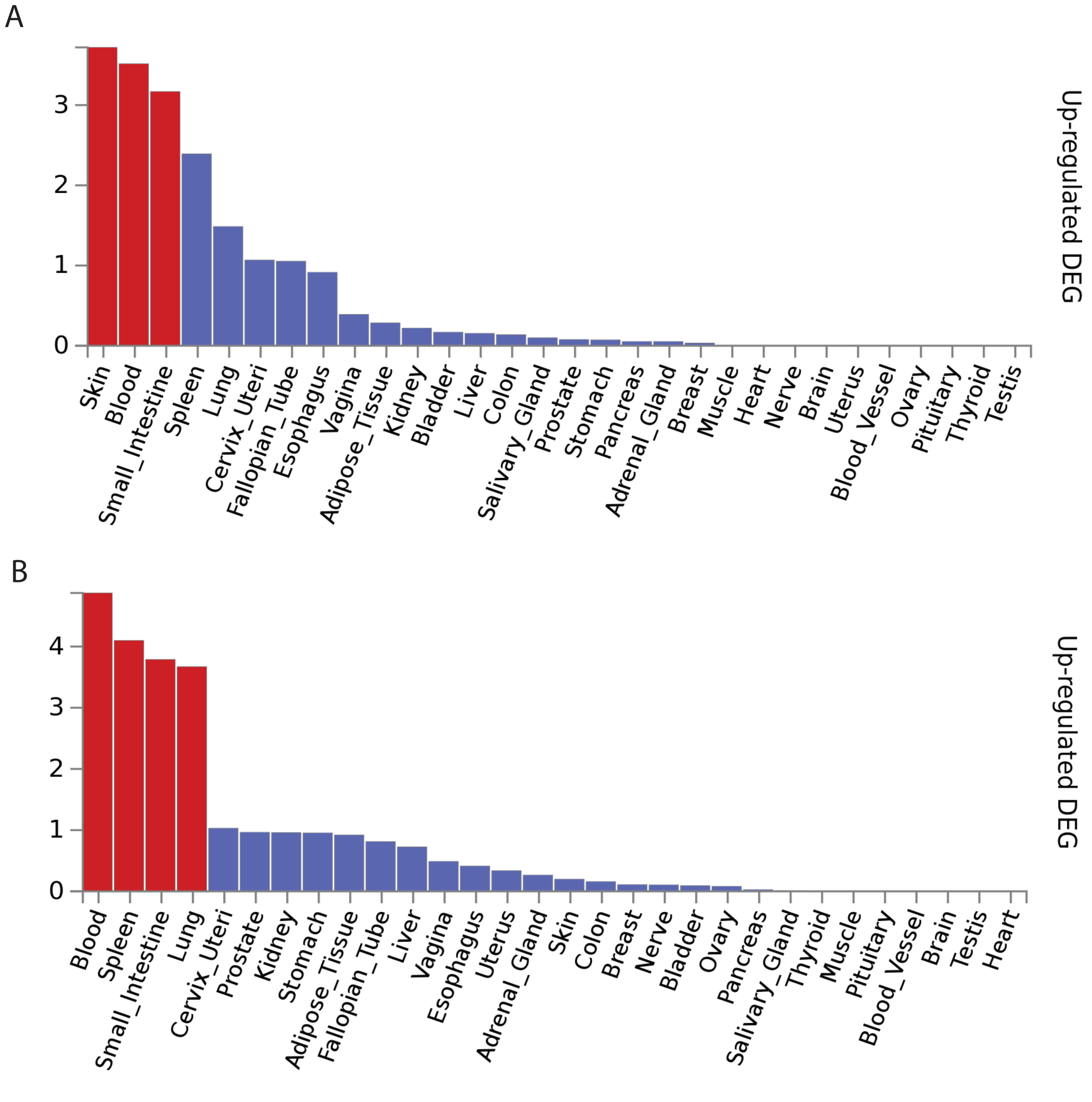

**Figure 8. PrediXcan genes in non-HLA regions associated with childhood onset and/or adult onset asthma in spleen and small intestine.** See Figure 4 in manuscript for additional details.

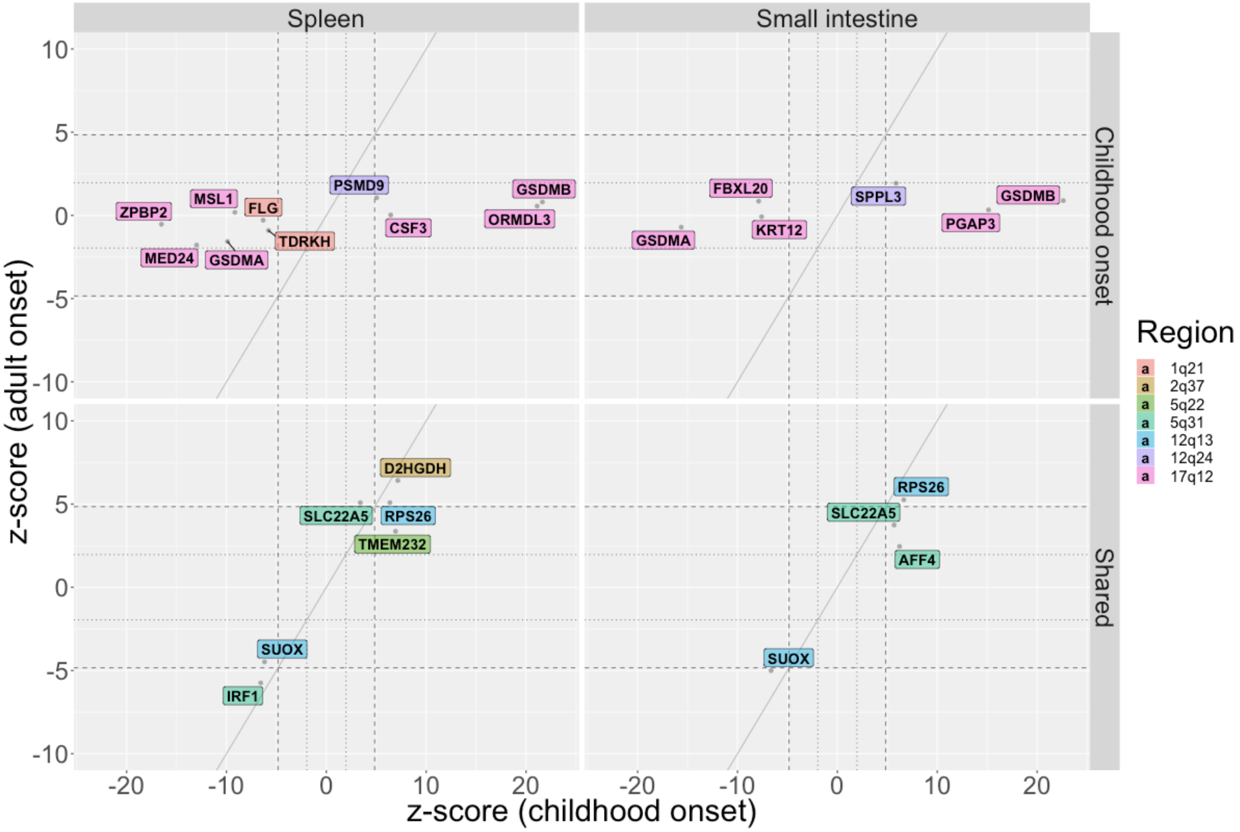

**Figure 9. PrediXcan genes in the HLA region associated in childhood onset and adult onset asthma in spleen and small intestine.** See Figure 5 in manuscript for additional details.

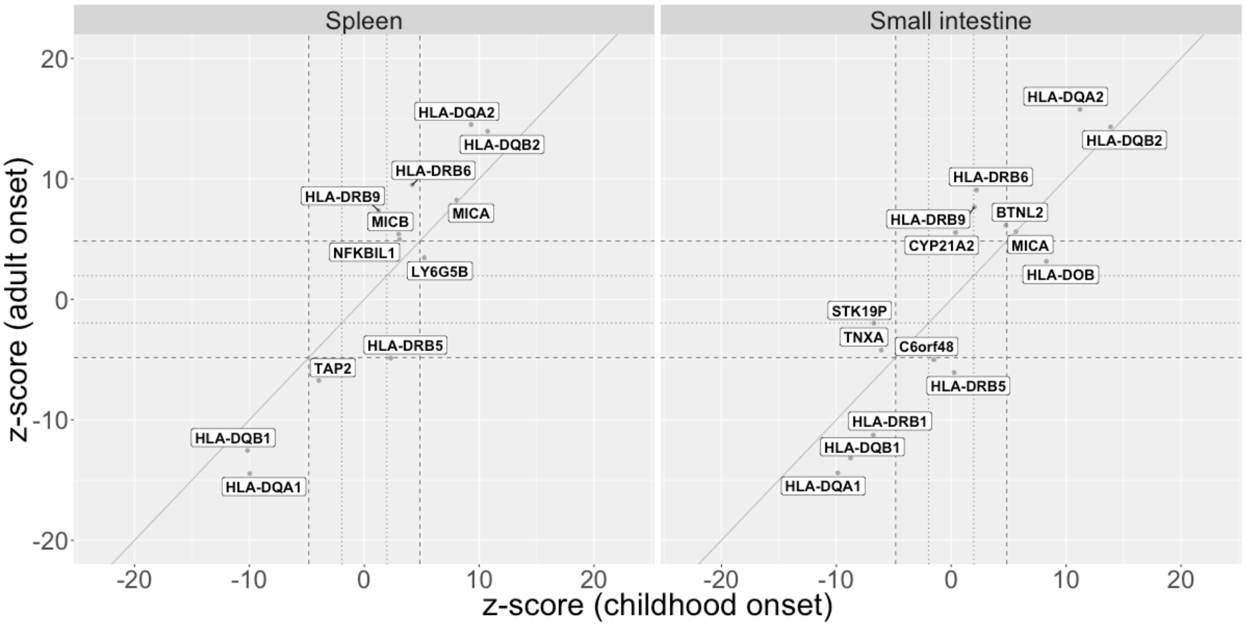

**Figure 10. Manhattan plot of COPD GWAS in the UKB (9,876 cases and 318,237 controls).**

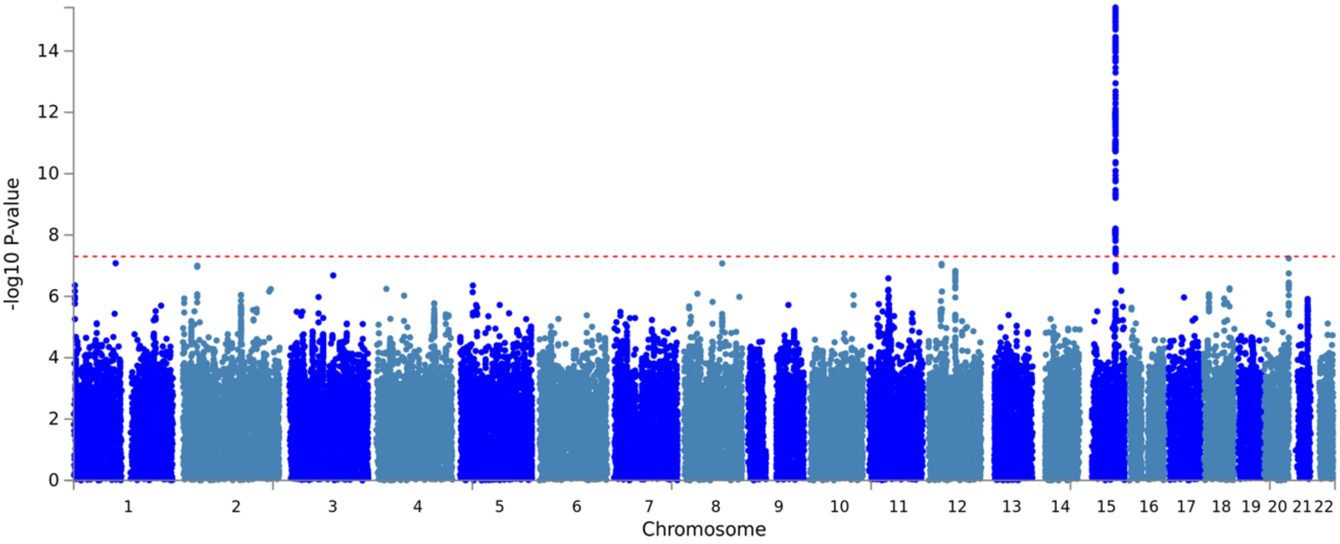

### Tables

**Table 1. PrediXcan prediction performance p-values.**

| Tissue | Gene Ensembl ID | Gene Symbol | Prediction performance | | Number of SNPs used |
|---|---|---|---|---|---|
| | | | R2 | P-value | |
| Lung | ENSG00000172057·5 | *ORMDL3* | 0·143 | 2·539E-13 | 8 |
| Lung | ENSG00000166949·11 | *SMAD3* | 0·024 | 5·450E-03 | 20 |
| Lung | ENSG00000169442·4 | *CD52* | 0·136 | 1·938E-12 | 64 |
| Lung | ENSG00000154252·11 | *GAL3ST2* | 0·096 | 9·324E-09 | 22 |
| Lung | ENSG00000073584·14 | *SMARCE1* | 0·165 | 1·117E-14 | 39 |
| Lung | ENSG00000204650·9 | *CRHR1-IT1* | 0·650 | 1·648E-88 | 142 |
| Lung | ENSG00000204516·5 | *MICB* | 0·329 | 1·391E-31 | 56 |
| Lung | ENSG00000115607·5 | *IL18RAP* | 0·090 | 2·061E-08 | 21 |
| Lung | ENSG00000204525·10 | *HLA-C* | 0·423 | 1·224E-44 | 44 |
| Lung | ENSG00000163221·7 | *S100A12* | 0·014 | 3·155E-02 | 40 |
| Lung | ENSG00000137337·10 | *MDC1* | 0·012 | 4·301E-02 | 57 |
| Lung | ENSG00000100412·11 | *ACO2* | 0·085 | 3·947E-08 | 19 |
| Lung | ENSG00000254810·1 | *RP11-672A2·4* | 0·084 | 5·383E-08 | 26 |
| Lung | ENSG00000132394·6 | *EEFSEC* | 0·046 | 8·542E-05 | 43 |
| Lung | ENSG00000229391·3 | *HLA-DRB6* | 0·671 | 2·219E-94 | 51 |
| Lung | ENSG00000240065·3 | *PSMB9* | 0·236 | 1·404E-21 | 71 |
| Lung | ENSG00000240053·8 | *LY6G5B* | 0·261 | 2·999E-24 | 54 |
| Lung | ENSG00000264070·1 | *DND1P1* | 0·537 | 5·558E-63 | 51 |
| Lung | ENSG00000137310·7 | *TCF19* | 0·064 | 3·562E-06 | 23 |
| Lung | ENSG00000244731·3 | *C4A* | 0·328 | 6·631E-32 | 51 |
| Lung | ENSG00000224389·4 | *C4B* | 0·112 | 2·231E-10 | 50 |
| Lung | ENSG00000204463·8 | *BAG6* | 0·049 | 4·966E-05 | 18 |
| Lung | ENSG00000164209·12 | *SLC25A46* | 0·012 | 4·585E-02 | 16 |
| Lung | ENSG00000134824·9 | *FADS2* | 0·093 | 1·332E-08 | 5 |
| Lung | ENSG00000204267·9 | *TAP2* | 0·130 | 3·480E-12 | 20 |
| Lung | ENSG00000213760·6 | *ATP6V1G2* | 0·131 | 6·280E-12 | 11 |
| Lung | ENSG00000108342·8 | *CSF3* | 0·017 | 2·039E-02 | 8 |
| Lung | ENSG00000204301·5 | *NOTCH4* | 0·235 | 1·234E-21 | 62 |
| Lung | ENSG00000237940·3 | *AC093642·3* | 0·093 | 1·563E-08 | 20 |

| | | | | | |
|---|---|---|---|---|---|
| Lung | ENSG00000198502·5 | *HLA-DRB5* | 0·627 | 3·574E-82 | 44 |
| Lung | ENSG00000186952·10 | *TMEM232* | 0·055 | 1·890E-05 | 51 |
| Lung | ENSG00000223501·4 | *VPS52* | 0·037 | 5·237E-04 | 4 |
| Lung | ENSG00000231852·2 | *CYP21A2* | 0·067 | 1·627E-06 | 28 |
| Lung | ENSG00000204438·6 | *GPANK1* | 0·033 | 9·513E-04 | 18 |
| Lung | ENSG00000113522·9 | *RAD50* | 0·176 | 7·418E-16 | 30 |
| Lung | ENSG00000077238·9 | *IL4R* | 0·107 | 8·294E-10 | 40 |
| Lung | ENSG00000263874·1 | *LINC00672* | 0·017 | 1·699E-02 | 13 |
| Lung | ENSG00000204444·6 | *APOM* | 0·012 | 4·893E-02 | 3 |
| Lung | ENSG00000105889·10 | *STEAP1B* | 0·130 | 1·283E-11 | 9 |
| Lung | ENSG00000197375·8 | *SLC22A5* | 0·126 | 2·378E-11 | 11 |
| Lung | ENSG00000146112·7 | *PPP1R18* | 0·056 | 1·534E-05 | 17 |
| Lung | ENSG00000143631·10 | *FLG* | 0·220 | 9·150E-20 | 17 |
| Lung | ENSG00000196301·3 | *HLA-DRB9* | 0·294 | 1·773E-27 | 134 |
| Lung | ENSG00000196126·6 | *HLA-DRB1* | 0·360 | 2·730E-35 | 29 |
| Lung | ENSG00000157837·11 | *SPPL3* | 0·124 | 3·508E-11 | 12 |
| Lung | ENSG00000204520·8 | *MICA* | 0·505 | 1·171E-57 | 50 |
| Lung | ENSG00000248290·1 | *TNXA* | 0·150 | 2·142E-13 | 131 |
| Lung | ENSG00000237541·3 | *HLA-DQA2* | 0·648 | 1·746E-88 | 46 |
| Lung | ENSG00000141744·3 | *PNMT* | 0·058 | 9·917E-06 | 15 |
| Lung | ENSG00000167914·6 | *GSDMA* | 0·375 | 6·415E-38 | 41 |
| Lung | ENSG00000196735·7 | *HLA-DQA1* | 0·519 | 8·282E-60 | 29 |
| Lung | ENSG00000232629·4 | *HLA-DQB2* | 0·524 | 1·731E-60 | 37 |
| Lung | ENSG00000161395·8 | *PGAP3* | 0·101 | 2·722E-09 | 9 |
| Lung | ENSG00000073605·14 | *GSDMB* | 0·131 | 9·445E-12 | 26 |
| Lung | ENSG00000204305·9 | *AGER* | 0·063 | 3·371E-06 | 18 |
| Lung | ENSG00000131437·11 | *KIF3A* | 0·016 | 2·055E-02 | 8 |
| Lung | ENSG00000179344·12 | *HLA-DQB1* | 0·634 | 2·591E-86 | 35 |
| Lung | ENSG00000180902·12 | *D2HGDH* | 0·366 | 1·267E-36 | 8 |
| Lung | ENSG00000137312·10 | *FLOT1* | 0·145 | 3·666E-13 | 34 |
| Lung | ENSG00000213171·2 | *LINGO4* | 0·080 | 1·363E-07 | 43 |
| Lung | ENSG00000159352·11 | *PSMD4* | 0·014 | 2·954E-02 | 48 |
| Lung | ENSG00000254470·2 | *AP5B1* | 0·112 | 3·379E-10 | 22 |

| | | | | | |
|---|---|---|---|---|---|
| Lung | ENSG00000115604·6 | *IL18R1* | 0·182 | 2·262E-16 | 123 |
| Lung | ENSG00000241106·2 | *HLA-DOB* | 0·527 | 1·356E-60 | 44 |
| Lung | ENSG00000231389·3 | *HLA-DPA1* | 0·050 | 4·911E-05 | 36 |
| Lung | ENSG00000182134·11 | *TDRKH* | 0·104 | 1·686E-09 | 17 |
| Lung | ENSG00000204427·7 | *ABHD16A* | 0·031 | 1·433E-03 | 53 |
| Lung | ENSG00000038532·10 | *CLEC16A* | 0·050 | 4·444E-05 | 8 |
| Lung | ENSG00000213676·6 | *ATF6B* | 0·153 | 1·521E-13 | 26 |
| Lung | ENSG00000188895·7 | *MSL1* | 0·113 | 3·577E-10 | 29 |
| Lung | ENSG00000139531·8 | *SUOX* | 0·286 | 1·589E-26 | 36 |
| Lung | ENSG00000197728·5 | *RPS26* | 0·675 | 7·725E-96 | 29 |
| Lung | ENSG00000008838·13 | *MED24* | 0·093 | 1·627E-08 | 25 |
| Lung | ENSG00000197712·7 | *FAM114A1* | 0·031 | 1·413E-03 | 55 |
| Lung | ENSG00000113520·6 | *IL4* | 0·021 | 8·424E-03 | 18 |
| Skin_Not_Sun_Exposed_Suprapubic | ENSG00000204657·2 | *OR2H2* | 0·033 | 2·372E-03 | 6 |
| Skin_Not_Sun_Exposed_Suprapubic | ENSG00000213171·2 | *LINGO4* | 0·263 | 9·100E-21 | 24 |
| Skin_Not_Sun_Exposed_Suprapubic | ENSG00000226644·1 | *RP11-128M1·1* | 0·126 | 7·065E-10 | 44 |
| Skin_Not_Sun_Exposed_Suprapubic | ENSG00000240053·8 | *LY6G5B* | 0·145 | 1·077E-11 | 17 |
| Skin_Not_Sun_Exposed_Suprapubic | ENSG00000113520·6 | *IL4* | 0·045 | 2·834E-04 | 40 |
| Skin_Not_Sun_Exposed_Suprapubic | ENSG00000231389·3 | *HLA-DPA1* | 0·131 | 2·989E-10 | 22 |
| Skin_Not_Sun_Exposed_Suprapubic | ENSG00000204650·9 | *CRHR1-IT1* | 0·707 | 1·301E-90 | 83 |
| Skin_Not_Sun_Exposed_Suprapubic | ENSG00000172057·5 | *ORMDL3* | 0·015 | 3·829E-02 | 37 |
| Skin_Not_Sun_Exposed_Suprapubic | ENSG00000131771·9 | *PPP1R1B* | 0·036 | 1·197E-03 | 9 |
| Skin_Not_Sun_Exposed_Suprapubic | ENSG00000161395·8 | *PGAP3* | 0·096 | 9·753E-08 | 15 |
| Skin_Not_Sun_Exposed_Suprapubic | ENSG00000115604·6 | *IL18R1* | 0·077 | 1·581E-06 | 58 |
| Skin_Not_Sun_Exposed_Suprapubic | ENSG00000264968·1 | *RP11-387H17·4* | 0·027 | 6·630E-03 | 3 |
| Skin_Not_Sun_Exposed_Suprapubic | ENSG00000179344·12 | *HLA-DQB1* | 0·709 | 3·446E-91 | 29 |
| Skin_Not_Sun_Exposed_Suprapubic | ENSG00000237541·3 | *HLA-DQA2* | 0·522 | 1·853E-52 | 41 |
| Skin_Not_Sun_Exposed_Suprapubic | ENSG00000141736·9 | *ERBB2* | 0·044 | 3·935E-04 | 33 |
| Skin_Not_Sun_Exposed_Suprapubic | ENSG00000229391·3 | *HLA-DRB6* | 0·595 | 4·320E-65 | 54 |
| Skin_Not_Sun_Exposed_Suprapubic | ENSG00000204267·9 | *TAP2* | 0·121 | 1·404E-09 | 58 |
| Skin_Not_Sun_Exposed_Suprapubic | ENSG00000231852·2 | *CYP21A2* | 0·150 | 1·561E-11 | 39 |
| Skin_Not_Sun_Exposed_Suprapubic | ENSG00000214546·3 | *AC087491·2* | 0·077 | 1·950E-06 | 21 |
| Skin_Not_Sun_Exposed_Suprapubic | ENSG00000213760·6 | *ATP6V1G2* | 0·162 | 2·174E-12 | 34 |

| | | | | | |
|---|---|---|---|---|---|
| Skin_Not_Sun_Exposed_Suprapubic | ENSG00000073605·14 | *GSDMB* | 0·028 | 5·239E-03 | 6 |
| Skin_Not_Sun_Exposed_Suprapubic | ENSG00000134824·9 | *FADS2* | 0·033 | 2·634E-03 | 4 |
| Skin_Not_Sun_Exposed_Suprapubic | ENSG00000232629·4 | *HLA-DQB2* | 0·038 | 1·111E-03 | 95 |
| Skin_Not_Sun_Exposed_Suprapubic | ENSG00000166949·11 | *SMAD3* | 0·034 | 2·005E-03 | 36 |
| Skin_Not_Sun_Exposed_Suprapubic | ENSG00000197712·7 | *FAM114A1* | 0·244 | 2·350E-19 | 46 |
| Skin_Not_Sun_Exposed_Suprapubic | ENSG00000198502·5 | *HLA-DRB5* | 0·698 | 1·943E-88 | 52 |
| Skin_Not_Sun_Exposed_Suprapubic | ENSG00000204438·6 | *GPANK1* | 0·136 | 1·686E-10 | 76 |
| Skin_Not_Sun_Exposed_Suprapubic | ENSG00000132394·6 | *EEFSEC* | 0·041 | 6·077E-04 | 13 |
| Skin_Not_Sun_Exposed_Suprapubic | ENSG00000137310·7 | *TCF19* | 0·081 | 8·387E-07 | 16 |
| Skin_Not_Sun_Exposed_Suprapubic | ENSG00000196301·3 | *HLA-DRB9* | 0·191 | 9·649E-15 | 65 |
| Skin_Not_Sun_Exposed_Suprapubic | ENSG00000204315·3 | *FKBPL* | 0·014 | 4·951E-02 | 7 |
| Skin_Not_Sun_Exposed_Suprapubic | ENSG00000204525·10 | *HLA-C* | 0·469 | 2·620E-43 | 40 |
| Skin_Not_Sun_Exposed_Suprapubic | ENSG00000244731·3 | *C4A* | 0·391 | 1·092E-34 | 55 |
| Skin_Not_Sun_Exposed_Suprapubic | ENSG00000224389·4 | *C4B* | 0·194 | 3·474E-15 | 34 |
| Skin_Not_Sun_Exposed_Suprapubic | ENSG00000196735·7 | *HLA-DQA1* | 0·409 | 8·522E-36 | 41 |
| Skin_Not_Sun_Exposed_Suprapubic | ENSG00000170426·1 | *SDR9C7* | 0·061 | 1·985E-05 | 7 |
| Skin_Not_Sun_Exposed_Suprapubic | ENSG00000188895·7 | *MSL1* | 0·100 | 4·158E-08 | 25 |
| Skin_Not_Sun_Exposed_Suprapubic | ENSG00000143536·7 | *CRNN* | 0·077 | 1·062E-06 | 30 |
| Skin_Not_Sun_Exposed_Suprapubic | ENSG00000073584·14 | *SMARCE1* | 0·163 | 6·187E-13 | 10 |
| Skin_Not_Sun_Exposed_Suprapubic | ENSG00000182134·11 | *TDRKH* | 0·085 | 7·084E-07 | 15 |
| Skin_Not_Sun_Exposed_Suprapubic | ENSG00000163216·6 | *SPRR2D* | 0·028 | 4·100E-03 | 12 |
| Skin_Not_Sun_Exposed_Suprapubic | ENSG00000196407·7 | *THEM5* | 0·021 | 1·616E-02 | 14 |
| Skin_Not_Sun_Exposed_Suprapubic | ENSG00000237940·3 | *AC093642·3* | 0·169 | 2·887E-13 | 22 |
| Skin_Not_Sun_Exposed_Suprapubic | ENSG00000204520·8 | *MICA* | 0·482 | 1·465E-45 | 62 |
| Skin_Not_Sun_Exposed_Suprapubic | ENSG00000266753·2 | *FBXL20* | 0·042 | 5·887E-04 | 7 |
| Skin_Not_Sun_Exposed_Suprapubic | ENSG00000254852·4 | *NPIPA2* | 0·088 | 2·067E-07 | 14 |
| Skin_Not_Sun_Exposed_Suprapubic | ENSG00000110917·3 | *MLEC* | 0·116 | 2·992E-09 | 70 |
| Skin_Not_Sun_Exposed_Suprapubic | ENSG00000186952·10 | *TMEM232* | 0·094 | 1·279E-07 | 16 |
| Skin_Not_Sun_Exposed_Suprapubic | ENSG00000264070·1 | *DND1P1* | 0·650 | 3·194E-76 | 28 |
| Skin_Not_Sun_Exposed_Suprapubic | ENSG00000204516·5 | *MICB* | 0·502 | 1·471E-49 | 92 |
| Skin_Not_Sun_Exposed_Suprapubic | ENSG00000204314·6 | *PRRT1* | 0·217 | 8·503E-17 | 70 |
| Skin_Not_Sun_Exposed_Suprapubic | ENSG00000241782·1 | *RP11-91P24·1* | 0·041 | 6·355E-04 | 7 |
| Skin_Not_Sun_Exposed_Suprapubic | ENSG00000180902·12 | *D2HGDH* | 0·439 | 7·217E-40 | 34 |

| | | | | | |
|---|---|---|---|---|---|
| Skin_Not_Sun_Exposed_Suprapubic | ENSG00000240065·3 | *PSMB9* | 0·088 | 3·054E-07 | 70 |
| Skin_Not_Sun_Exposed_Suprapubic | ENSG00000145777·10 | *TSLP* | 0·261 | 1·366E-20 | 20 |
| Skin_Not_Sun_Exposed_Suprapubic | ENSG00000166396·8 | *SERPINB7* | 0·132 | 1·146E-10 | 9 |
| Skin_Not_Sun_Exposed_Suprapubic | ENSG00000143631·10 | *FLG* | 0·029 | 3·948E-03 | 22 |
| Skin_Not_Sun_Exposed_Suprapubic | ENSG00000196126·6 | *HLA-DRB1* | 0·296 | 5·786E-24 | 29 |
| Skin_Not_Sun_Exposed_Suprapubic | ENSG00000248290·1 | *TNXA* | 0·088 | 3·842E-07 | 75 |
| Skin_Not_Sun_Exposed_Suprapubic | ENSG00000008838·13 | *MED24* | 0·136 | 8·432E-11 | 18 |
| Skin_Not_Sun_Exposed_Suprapubic | ENSG00000164402·9 | *SEPT8* | 0·173 | 1·255E-13 | 16 |
| Skin_Not_Sun_Exposed_Suprapubic | ENSG00000154252·11 | *GAL3ST2* | 0·053 | 9·932E-05 | 5 |
| Skin_Not_Sun_Exposed_Suprapubic | ENSG00000115598·5 | *IL1RL2* | 0·085 | 6·599E-07 | 46 |
| Skin_Not_Sun_Exposed_Suprapubic | ENSG00000174130·8 | *TLR6* | 0·210 | 3·392E-16 | 21 |
| Skin_Not_Sun_Exposed_Suprapubic | ENSG00000213676·6 | *ATF6B* | 0·138 | 1·241E-10 | 54 |
| Skin_Not_Sun_Exposed_Suprapubic | ENSG00000131435·8 | *PDLIM4* | 0·066 | 1·452E-05 | 14 |
| Skin_Not_Sun_Exposed_Suprapubic | ENSG00000197728·5 | *RPS26* | 0·759 | 1·092E-109 | 30 |
| Skin_Not_Sun_Exposed_Suprapubic | ENSG00000169509·5 | *CRCT1* | 0·060 | 3·740E-05 | 29 |
| Skin_Not_Sun_Exposed_Suprapubic | ENSG00000139531·8 | *SUOX* | 0·233 | 6·339E-19 | 11 |
| Skin_Not_Sun_Exposed_Suprapubic | ENSG00000197375·8 | *SLC22A5* | 0·287 | 2·716E-23 | 14 |
| Skin_Not_Sun_Exposed_Suprapubic | ENSG00000115607·5 | *IL18RAP* | 0·066 | 1·399E-05 | 42 |
| Skin_Not_Sun_Exposed_Suprapubic | ENSG00000204301·5 | *NOTCH4* | 0·283 | 2·760E-23 | 30 |
| Skin_Not_Sun_Exposed_Suprapubic | ENSG00000204427·7 | *ABHD16A* | 0·040 | 7·840E-04 | 11 |
| Skin_Not_Sun_Exposed_Suprapubic | ENSG00000241106·2 | *HLA-DOB* | 0·495 | 2·183E-48 | 50 |
| Skin_Sun_Exposed_Lower_leg | ENSG00000204444·6 | *APOM* | 0·021 | 6·705E-03 | 25 |
| Skin_Sun_Exposed_Lower_leg | ENSG00000163216·6 | *SPRR2D* | 0·099 | 1·369E-09 | 49 |
| Skin_Sun_Exposed_Lower_leg | ENSG00000204427·7 | *ABHD16A* | 0·012 | 4·062E-02 | 27 |
| Skin_Sun_Exposed_Lower_leg | ENSG00000204525·10 | *HLA-C* | 0·577 | 7·147E-78 | 98 |
| Skin_Sun_Exposed_Lower_leg | ENSG00000105889·10 | *STEAP1B* | 0·016 | 1·775E-02 | 16 |
| Skin_Sun_Exposed_Lower_leg | ENSG00000254852·4 | *NPIPA2* | 0·017 | 1·574E-02 | 7 |
| Skin_Sun_Exposed_Lower_leg | ENSG00000204315·3 | *FKBPL* | 0·028 | 1·635E-03 | 15 |
| Skin_Sun_Exposed_Lower_leg | ENSG00000229391·3 | *HLA-DRB6* | 0·617 | 1·272E-86 | 43 |
| Skin_Sun_Exposed_Lower_leg | ENSG00000231852·2 | *CYP21A2* | 0·193 | 1·526E-18 | 32 |
| Skin_Sun_Exposed_Lower_leg | ENSG00000248290·1 | *TNXA* | 0·071 | 3·603E-07 | 30 |
| Skin_Sun_Exposed_Lower_leg | ENSG00000240053·8 | *LY6G5B* | 0·225 | 6·757E-22 | 28 |
| Skin_Sun_Exposed_Lower_leg | ENSG00000228727·4 | *SAPCD1* | 0·042 | 8·167E-05 | 31 |

| | | | | | |
|---|---|---|---|---|---|
| Skin_Sun_Exposed_Lower_leg | ENSG00000204516·5 | *MICB* | 0·445 | 8·909E-52 | 43 |
| Skin_Sun_Exposed_Lower_leg | ENSG00000264070·1 | *DND1P1* | 0·623 | 5·574E-90 | 29 |
| Skin_Sun_Exposed_Lower_leg | ENSG00000132394·6 | *EEFSEC* | 0·080 | 5·593E-08 | 16 |
| Skin_Sun_Exposed_Lower_leg | ENSG00000244731·3 | *C4A* | 0·282 | 9·421E-29 | 64 |
| Skin_Sun_Exposed_Lower_leg | ENSG00000164209·12 | *SLC25A46* | 0·066 | 8·972E-07 | 34 |
| Skin_Sun_Exposed_Lower_leg | ENSG00000224389·4 | *C4B* | 0·198 | 6·450E-19 | 51 |
| Skin_Sun_Exposed_Lower_leg | ENSG00000204650·9 | *CRHR1-IT1* | 0·721 | 4·585E-120 | 222 |
| Skin_Sun_Exposed_Lower_leg | ENSG00000204463·8 | *BAG6* | 0·099 | 5·211E-10 | 37 |
| Skin_Sun_Exposed_Lower_leg | ENSG00000073605·14 | *GSDMB* | 0·061 | 2·487E-06 | 26 |
| Skin_Sun_Exposed_Lower_leg | ENSG00000263874·1 | *LINC00672* | 0·071 | 3·581E-07 | 39 |
| Skin_Sun_Exposed_Lower_leg | ENSG00000204314·6 | *PRRT1* | 0·284 | 1·660E-29 | 36 |
| Skin_Sun_Exposed_Lower_leg | ENSG00000134824·9 | *FADS2* | 0·032 | 7·800E-04 | 13 |
| Skin_Sun_Exposed_Lower_leg | ENSG00000254470·2 | *AP5B1* | 0·022 | 5·141E-03 | 36 |
| Skin_Sun_Exposed_Lower_leg | ENSG00000223501·4 | *VPS52* | 0·013 | 3·579E-02 | 27 |
| Skin_Sun_Exposed_Lower_leg | ENSG00000213760·6 | *ATP6V1G2* | 0·114 | 4·153E-11 | 18 |
| Skin_Sun_Exposed_Lower_leg | ENSG00000110801·9 | *PSMD9* | 0·043 | 9·339E-05 | 11 |
| Skin_Sun_Exposed_Lower_leg | ENSG00000196407·7 | *THEM5* | 0·028 | 1·715E-03 | 21 |
| Skin_Sun_Exposed_Lower_leg | ENSG00000204267·9 | *TAP2* | 0·215 | 6·009E-21 | 80 |
| Skin_Sun_Exposed_Lower_leg | ENSG00000204287·9 | *HLA-DRA* | 0·090 | 5·285E-09 | 27 |
| Skin_Sun_Exposed_Lower_leg | ENSG00000204420·4 | *C6orf25* | 0·032 | 7·022E-04 | 7 |
| Skin_Sun_Exposed_Lower_leg | ENSG00000204301·5 | *NOTCH4* | 0·225 | 9·104E-22 | 45 |
| Skin_Sun_Exposed_Lower_leg | ENSG00000143631·10 | *FLG* | 0·071 | 2·348E-07 | 33 |
| Skin_Sun_Exposed_Lower_leg | ENSG00000204657·2 | *OR2H2* | 0·014 | 2·583E-02 | 39 |
| Skin_Sun_Exposed_Lower_leg | ENSG00000137312·10 | *FLOT1* | 0·061 | 1·587E-06 | 16 |
| Skin_Sun_Exposed_Lower_leg | ENSG00000197712·7 | *FAM114A1* | 0·402 | 1·334E-45 | 23 |
| Skin_Sun_Exposed_Lower_leg | ENSG00000169442·4 | *CD52* | 0·051 | 1·314E-05 | 29 |
| Skin_Sun_Exposed_Lower_leg | ENSG00000198502·5 | *HLA-DRB5* | 0·701 | 3·765E-113 | 46 |
| Skin_Sun_Exposed_Lower_leg | ENSG00000204305·9 | *AGER* | 0·057 | 5·332E-06 | 22 |
| Skin_Sun_Exposed_Lower_leg | ENSG00000174130·8 | *TLR6* | 0·214 | 9·039E-21 | 19 |
| Skin_Sun_Exposed_Lower_leg | ENSG00000237940·3 | *AC093642·3* | 0·123 | 7·301E-12 | 14 |
| Skin_Sun_Exposed_Lower_leg | ENSG00000143536·7 | *CRNN* | 0·219 | 3·501E-21 | 63 |
| Skin_Sun_Exposed_Lower_leg | ENSG00000231389·3 | *HLA-DPA1* | 0·266 | 1·940E-26 | 16 |
| Skin_Sun_Exposed_Lower_leg | ENSG00000241106·2 | *HLA-DOB* | 0·523 | 3·981E-65 | 63 |

| | | | | | |
|---|---|---|---|---|---|
| Skin_Sun_Exposed_Lower_leg | ENSG00000145777·10 | *TSLP* | 0·247 | 6·762E-25 | 28 |
| Skin_Sun_Exposed_Lower_leg | ENSG00000232629·4 | *HLA-DQB2* | 0·045 | 5·052E-05 | 66 |
| Skin_Sun_Exposed_Lower_leg | ENSG00000008838·13 | *MED24* | 0·212 | 7·541E-21 | 40 |
| Skin_Sun_Exposed_Lower_leg | ENSG00000125347·9 | *IRF1* | 0·013 | 2·967E-02 | 8 |
| Skin_Sun_Exposed_Lower_leg | ENSG00000264968·1 | *RP11-387H17·4* | 0·071 | 3·630E-07 | 18 |
| Skin_Sun_Exposed_Lower_leg | ENSG00000167914·6 | *GSDMA* | 0·056 | 7·190E-06 | 9 |
| Skin_Sun_Exposed_Lower_leg | ENSG00000172057·5 | *ORMDL3* | 0·018 | 1·111E-02 | 21 |
| Skin_Sun_Exposed_Lower_leg | ENSG00000141736·9 | *ERBB2* | 0·043 | 7·548E-05 | 20 |
| Skin_Sun_Exposed_Lower_leg | ENSG00000240065·3 | *PSMB9* | 0·100 | 1·360E-09 | 45 |
| Skin_Sun_Exposed_Lower_leg | ENSG00000131771·9 | *PPP1R1B* | 0·041 | 1·418E-04 | 17 |
| Skin_Sun_Exposed_Lower_leg | ENSG00000179344·12 | *HLA-DQB1* | 0·668 | 1·291E-101 | 29 |
| Skin_Sun_Exposed_Lower_leg | ENSG00000115604·6 | *IL18R1* | 0·086 | 1·491E-08 | 41 |
| Skin_Sun_Exposed_Lower_leg | ENSG00000115607·5 | *IL18RAP* | 0·044 | 5·443E-05 | 41 |
| Skin_Sun_Exposed_Lower_leg | ENSG00000237541·3 | *HLA-DQA2* | 0·660 | 9·621E-100 | 56 |
| Skin_Sun_Exposed_Lower_leg | ENSG00000196735·7 | *HLA-DQA1* | 0·382 | 1·226E-41 | 49 |
| Skin_Sun_Exposed_Lower_leg | ENSG00000131437·11 | *KIF3A* | 0·037 | 2·450E-04 | 13 |
| Skin_Sun_Exposed_Lower_leg | ENSG00000204520·8 | *MICA* | 0·461 | 1·002E-53 | 88 |
| Skin_Sun_Exposed_Lower_leg | ENSG00000188895·7 | *MSL1* | 0·143 | 1·197E-13 | 41 |
| Skin_Sun_Exposed_Lower_leg | ENSG00000161395·8 | *PGAP3* | 0·091 | 6·666E-09 | 41 |
| Skin_Sun_Exposed_Lower_leg | ENSG00000113520·6 | *IL4* | 0·039 | 1·481E-04 | 22 |
| Skin_Sun_Exposed_Lower_leg | ENSG00000139531·8 | *SUOX* | 0·369 | 2·817E-40 | 14 |
| Skin_Sun_Exposed_Lower_leg | ENSG00000196126·6 | *HLA-DRB1* | 0·321 | 2·123E-33 | 46 |
| Skin_Sun_Exposed_Lower_leg | ENSG00000186952·10 | *TMEM232* | 0·059 | 3·198E-06 | 17 |
| Skin_Sun_Exposed_Lower_leg | ENSG00000119408·12 | *NEK6* | 0·034 | 4·919E-04 | 31 |
| Skin_Sun_Exposed_Lower_leg | ENSG00000169189·12 | *NSMCE1* | 0·025 | 2·689E-03 | 6 |
| Skin_Sun_Exposed_Lower_leg | ENSG00000182134·11 | *TDRKH* | 0·084 | 2·944E-08 | 15 |
| Skin_Sun_Exposed_Lower_leg | ENSG00000226644·1 | *RP11-128M1·1* | 0·029 | 1·086E-03 | 19 |
| Skin_Sun_Exposed_Lower_leg | ENSG00000166949·11 | *SMAD3* | 0·054 | 6·257E-06 | 16 |
| Skin_Sun_Exposed_Lower_leg | ENSG00000170426·1 | *SDR9C7* | 0·027 | 1·919E-03 | 17 |
| Skin_Sun_Exposed_Lower_leg | ENSG00000110917·3 | *MLEC* | 0·143 | 5·962E-14 | 35 |
| Skin_Sun_Exposed_Lower_leg | ENSG00000213171·2 | *LINGO4* | 0·335 | 1·283E-34 | 70 |
| Skin_Sun_Exposed_Lower_leg | ENSG00000123384·9 | *LRP1* | 0·046 | 4·747E-05 | 17 |
| Skin_Sun_Exposed_Lower_leg | ENSG00000166396·8 | *SERPINB7* | 0·028 | 1·675E-03 | 24 |

| | | | | | |
|---|---|---|---|---|---|
| Skin_Sun_Exposed_Lower_leg | ENSG00000131435·8 | *PDLIM4* | 0·187 | 5·720E-18 | 19 |
| Skin_Sun_Exposed_Lower_leg | ENSG00000113522·9 | *RAD50* | 0·018 | 1·156E-02 | 44 |
| Skin_Sun_Exposed_Lower_leg | ENSG00000137310·7 | *TCF19* | 0·150 | 1·529E-14 | 38 |
| Skin_Sun_Exposed_Lower_leg | ENSG00000115598·5 | *IL1RL2* | 0·137 | 5·314E-13 | 66 |
| Skin_Sun_Exposed_Lower_leg | ENSG00000197375·8 | *SLC22A5* | 0·303 | 1·258E-30 | 28 |
| Skin_Sun_Exposed_Lower_leg | ENSG00000197728·5 | *RPS26* | 0·698 | 2·508E-113 | 43 |
| Skin_Sun_Exposed_Lower_leg | ENSG00000180902·12 | *D2HGDH* | 0·459 | 2·611E-53 | 18 |
| Skin_Sun_Exposed_Lower_leg | ENSG00000154252·11 | *GAL3ST2* | 0·058 | 3·330E-06 | 4 |
| Skin_Sun_Exposed_Lower_leg | ENSG00000164402·9 | *SEPT8* | 0·342 | 6·760E-36 | 22 |
| Skin_Sun_Exposed_Lower_leg | ENSG00000213676·6 | *ATF6B* | 0·153 | 1·580E-14 | 108 |
| Skin_Sun_Exposed_Lower_leg | ENSG00000073584·14 | *SMARCE1* | 0·237 | 3·980E-23 | 113 |
| Skin_Sun_Exposed_Lower_leg | ENSG00000196301·3 | *HLA-DRB9* | 0·033 | 5·479E-04 | 66 |
| Spleen | ENSG00000240053·8 | *LY6G5B* | 0·052 | 1·387E-02 | 10 |
| Spleen | ENSG00000182134·11 | *TDRKH* | 0·242 | 1·911E-08 | 15 |
| Spleen | ENSG00000139531·8 | *SUOX* | 0·389 | 6·252E-16 | 12 |
| Spleen | ENSG00000143631·10 | *FLG* | 0·126 | 1·290E-04 | 16 |
| Spleen | ENSG00000197728·5 | *RPS26* | 0·725 | 5·830E-41 | 18 |
| Spleen | ENSG00000108342·8 | *CSF3* | 0·153 | 2·322E-05 | 36 |
| Spleen | ENSG00000125347·9 | *IRF1* | 0·059 | 7·257E-03 | 4 |
| Spleen | ENSG00000186952·10 | *TMEM232* | 0·062 | 6·144E-03 | 49 |
| Spleen | ENSG00000180902·12 | *D2HGDH* | 0·347 | 1·006E-12 | 8 |
| Spleen | ENSG00000204520·8 | *MICA* | 0·686 | 1·035E-34 | 29 |
| Spleen | ENSG00000221946·3 | *FXYD7* | 0·072 | 2·165E-03 | 5 |
| Spleen | ENSG00000137337·10 | *MDC1* | 0·099 | 5·640E-04 | 67 |
| Spleen | ENSG00000237541·3 | *HLA-DQA2* | 0·632 | 3·549E-30 | 27 |
| Spleen | ENSG00000167914·6 | *GSDMA* | 0·213 | 1·482E-07 | 4 |
| Spleen | ENSG00000196735·7 | *HLA-DQA1* | 0·345 | 3·359E-12 | 59 |
| Spleen | ENSG00000179344·12 | *HLA-DQB1* | 0·688 | 1·406E-35 | 61 |
| Spleen | ENSG00000232629·4 | *HLA-DQB2* | 0·625 | 3·325E-36 | 39 |
| Spleen | ENSG00000008838·13 | *MED24* | 0·082 | 1·610E-03 | 20 |
| Spleen | ENSG00000186075·8 | *ZPBP2* | 0·105 | 2·889E-04 | 12 |
| Spleen | ENSG00000172057·5 | *ORMDL3* | 0·474 | 5·820E-20 | 46 |
| Spleen | ENSG00000073605·14 | *GSDMB* | 0·490 | 3·154E-19 | 12 |

| | | | | | |
|---|---|---|---|---|---|
| Spleen | ENSG00000164402·9 | *SEPT8* | 0·043 | 2·889E-02 | 11 |
| Spleen | ENSG00000188895·7 | *MSL1* | 0·182 | 1·104E-06 | 7 |
| Spleen | ENSG00000110801·9 | *PSMD9* | 0·071 | 4·123E-03 | 27 |
| Spleen | ENSG00000244731·3 | *C4A* | 0·196 | 1·001E-07 | 79 |
| Spleen | ENSG00000213719·4 | *CLIC1* | 0·105 | 5·656E-04 | 10 |
| Spleen | ENSG00000240065·3 | *PSMB9* | 0·131 | 4·269E-05 | 20 |
| Spleen | ENSG00000204650·9 | *CRHR1-IT1* | 0·583 | 8·850E-25 | 24 |
| Spleen | ENSG00000134824·9 | *FADS2* | 0·523 | 1·962E-21 | 26 |
| Spleen | ENSG00000237940·3 | *AC093642·3* | 0·120 | 5·348E-05 | 67 |
| Spleen | ENSG00000163216·6 | *SPRR2D* | 0·061 | 9·541E-03 | 8 |
| Spleen | ENSG00000204301·5 | *NOTCH4* | 0·159 | 1·474E-05 | 4 |
| Spleen | ENSG00000196301·3 | *HLA-DRB9* | 0·297 | 6·692E-11 | 130 |
| Spleen | ENSG00000197712·7 | *FAM114A1* | 0·144 | 3·654E-05 | 53 |
| Spleen | ENSG00000254810·1 | *RP11-672A2·4* | 0·054 | 1·327E-02 | 17 |
| Spleen | ENSG00000198502·5 | *HLA-DRB5* | 0·535 | 1·000E-22 | 13 |
| Spleen | ENSG00000241106·2 | *HLA-DOB* | 0·419 | 7·045E-16 | 77 |
| Spleen | ENSG00000204516·5 | *MICB* | 0·263 | 2·624E-10 | 41 |
| Spleen | ENSG00000204498·6 | *NFKBIL1* | 0·125 | 5·668E-05 | 47 |
| Spleen | ENSG00000196126·6 | *HLA-DRB1* | 0·300 | 1·488E-10 | 26 |
| Spleen | ENSG00000073584·14 | *SMARCE1* | 0·100 | 5·344E-04 | 13 |
| Spleen | ENSG00000197375·8 | *SLC22A5* | 0·115 | 1·439E-04 | 88 |
| Spleen | ENSG00000204267·9 | *TAP2* | 0·291 | 3·257E-10 | 48 |
| Spleen | ENSG00000229391·3 | *HLA-DRB6* | 0·620 | 1·822E-28 | 117 |
| Spleen | ENSG00000137310·7 | *TCF19* | 0·278 | 1·706E-11 | 31 |
| Spleen | ENSG00000264070·1 | *DND1P1* | 0·413 | 5·303E-15 | 166 |
| Spleen | ENSG00000145777·10 | *TSLP* | 0·485 | 6·884E-21 | 15 |
| Spleen | ENSG00000204525·10 | *HLA-C* | 0·298 | 3·360E-12 | 86 |
| Whole_Blood | ENSG00000131748·11 | *STARD3* | 0·079 | 5·283E-07 | 29 |
| Whole_Blood | ENSG00000115009·7 | *CCL20* | 0·136 | 9·696E-12 | 3 |
| Whole_Blood | ENSG00000197728·5 | *RPS26* | 0·587 | 4·336E-70 | 25 |
| Whole_Blood | ENSG00000139531·8 | *SUOX* | 0·142 | 4·811E-12 | 11 |
| Whole_Blood | ENSG00000174123·6 | *TLR10* | 0·033 | 1·334E-03 | 42 |
| Whole_Blood | ENSG00000240065·3 | *PSMB9* | 0·325 | 1·044E-29 | 66 |

| | | | | | |
|---|---|---|---|---|---|
| Whole_Blood | ENSG00000196301·3 | *HLA-DRB9* | 0·151 | 5·090E-13 | 27 |
| Whole_Blood | ENSG00000077238·9 | *IL4R* | 0·089 | 5·858E-08 | 9 |
| Whole_Blood | ENSG00000115604·6 | *IL18R1* | 0·039 | 4·249E-04 | 46 |
| Whole_Blood | ENSG00000204301·5 | *NOTCH4* | 0·436 | 3·812E-43 | 59 |
| Whole_Blood | ENSG00000255362·1 | *RP11-619A14·3* | 0·029 | 2·659E-03 | 9 |
| Whole_Blood | ENSG00000188895·7 | *MSL1* | 0·039 | 4·846E-04 | 28 |
| Whole_Blood | ENSG00000204267·9 | *TAP2* | 0·158 | 1·359E-13 | 35 |
| Whole_Blood | ENSG00000115607·5 | *IL18RAP* | 0·404 | 7·052E-39 | 90 |
| Whole_Blood | ENSG00000232629·4 | *HLA-DQB2* | 0·558 | 1·608E-63 | 46 |
| Whole_Blood | ENSG00000197375·8 | *SLC22A5* | 0·167 | 4·289E-14 | 46 |
| Whole_Blood | ENSG00000172057·5 | *ORMDL3* | 0·279 | 5·693E-25 | 30 |
| Whole_Blood | ENSG00000264198·1 | *IKZF3* | 0·023 | 6·740E-03 | 3 |
| Whole_Blood | ENSG00000073605·14 | *GSDMB* | 0·302 | 2·135E-27 | 20 |
| Whole_Blood | ENSG00000205716·4 | *AC007248·6* | 0·025 | 5·414E-03 | 8 |
| Whole_Blood | ENSG00000196735·7 | *HLA-DQA1* | 0·226 | 1·294E-19 | 8 |
| Whole_Blood | ENSG00000179344·12 | *HLA-DQB1* | 0·571 | 5·188E-66 | 42 |
| Whole_Blood | ENSG00000204520·8 | *MICA* | 0·189 | 1·583E-16 | 43 |
| Whole_Blood | ENSG00000244731·3 | *C4A* | 0·168 | 1·321E-14 | 13 |
| Whole_Blood | ENSG00000213676·6 | *ATF6B* | 0·025 | 5·859E-03 | 13 |
| Whole_Blood | ENSG00000008838·13 | *MED24* | 0·052 | 4·423E-05 | 8 |
| Whole_Blood | ENSG00000237541·3 | *HLA-DQA2* | 0·650 | 3·498E-84 | 105 |
| Whole_Blood | ENSG00000113520·6 | *IL4* | 0·019 | 1·475E-02 | 4 |
| Whole_Blood | ENSG00000204315·3 | *FKBPL* | 0·036 | 7·876E-04 | 88 |
| Whole_Blood | ENSG00000204390·8 | *HSPA1L* | 0·019 | 1·585E-02 | 5 |
| Whole_Blood | ENSG00000213171·2 | *LINGO4* | 0·059 | 1·395E-05 | 23 |
| Whole_Blood | ENSG00000164209·12 | *SLC25A46* | 0·065 | 5·360E-06 | 29 |
| Whole_Blood | ENSG00000204366·3 | *ZBTB12* | 0·291 | 7·007E-27 | 15 |
| Whole_Blood | ENSG00000204525·10 | *HLA-C* | 0·315 | 3·614E-29 | 19 |
| Whole_Blood | ENSG00000163221·7 | *S100A12* | 0·174 | 7·580E-15 | 37 |
| Whole_Blood | ENSG00000241106·2 | *HLA-DOB* | 0·480 | 1·752E-50 | 36 |
| Whole_Blood | ENSG00000196126·6 | *HLA-DRB1* | 0·159 | 7·772E-14 | 20 |
| Whole_Blood | ENSG00000204305·9 | *AGER* | 0·067 | 4·282E-06 | 26 |
| Whole_Blood | ENSG00000134824·9 | *FADS2* | 0·528 | 1·480E-57 | 25 |

| | | | | | |
|---|---|---|---|---|---|
| Whole_Blood | ENSG00000204463·8 | *BAG6* | 0·034 | 1·136E-03 | 29 |
| Whole_Blood | ENSG00000226644·1 | *RP11-128M1·1* | 0·059 | 8·861E-06 | 5 |
| Whole_Blood | ENSG00000119408·12 | *NEK6* | 0·121 | 2·158E-10 | 8 |
| Whole_Blood | ENSG00000263874·1 | *LINC00672* | 0·038 | 6·672E-04 | 33 |
| Whole_Blood | ENSG00000159352·11 | *PSMD4* | 0·014 | 3·904E-02 | 14 |
| Whole_Blood | ENSG00000110917·3 | *MLEC* | 0·038 | 4·668E-04 | 9 |
| Whole_Blood | ENSG00000213760·6 | *ATP6V1G2* | 0·018 | 2·002E-02 | 19 |
| Whole_Blood | ENSG00000204657·2 | *OR2H2* | 0·031 | 1·764E-03 | 15 |
| Whole_Blood | ENSG00000157837·11 | *SPPL3* | 0·020 | 1·290E-02 | 40 |
| Whole_Blood | ENSG00000254810·1 | *RP11-672A2·4* | 0·056 | 2·302E-05 | 31 |
| Whole_Blood | ENSG00000198502·5 | *HLA-DRB5* | 0·606 | 3·017E-74 | 50 |
| Whole_Blood | ENSG00000224389·4 | *C4B* | 0·222 | 1·305E-19 | 41 |
| Whole_Blood | ENSG00000137310·7 | *TCF19* | 0·072 | 1·815E-06 | 18 |
| Whole_Blood | ENSG00000254852·4 | *NPIPA2* | 0·024 | 6·332E-03 | 8 |
| Whole_Blood | ENSG00000166949·11 | *SMAD3* | 0·032 | 1·649E-03 | 54 |
| Whole_Blood | ENSG00000180902·12 | *D2HGDH* | 0·245 | 1·608E-21 | 14 |
| Whole_Blood | ENSG00000204650·9 | *CRHR1-IT1* | 0·455 | 1·385E-46 | 23 |
| Whole_Blood | ENSG00000169442·4 | *CD52* | 0·237 | 1·130E-20 | 41 |
| Whole_Blood | ENSG00000182134·11 | *TDRKH* | 0·086 | 1·691E-07 | 11 |
| Whole_Blood | ENSG00000264070·1 | *DND1P1* | 0·540 | 7·495E-60 | 21 |
| Whole_Blood | ENSG00000213719·4 | *CLIC1* | 0·026 | 4·221E-03 | 9 |
| Whole_Blood | ENSG00000137312·10 | *FLOT1* | 0·135 | 1·835E-11 | 40 |
| Whole_Blood | ENSG00000240053·8 | *LY6G5B* | 0·071 | 2·126E-06 | 37 |
| Whole_Blood | ENSG00000167914·6 | *GSDMA* | 0·122 | 6·011E-11 | 20 |
| Whole_Blood | ENSG00000204516·5 | *MICB* | 0·339 | 4·070E-31 | 45 |
| Whole_Blood | ENSG00000231852·2 | *CYP21A2* | 0·046 | 1·189E-04 | 19 |
| Whole_Blood | ENSG00000229391·3 | *HLA-DRB6* | 0·561 | 7·405E-65 | 77 |
| Whole_Blood | ENSG00000131435·8 | *PDLIM4* | 0·024 | 5·908E-03 | 10 |

### Table 2. Comparison of new loci in UKB asthma GWAS and Demenais et al[6] meta-analysis of asthma GWASs.

| Locus [a] | rsID | Position [b] | Nearby Genes [c] | Allele [d] | RAF [e] | UKB Childhood Onset Asthma GWAS | | | UKB Adult Onset Asthma GWAS | | | UKB Age of Onset GWAS | | | Demenais *et al* (TAGC)[6] | | | | | | | | | |
|---|---|---|---|---|---|---|---|---|---|---|---|---|---|---|---|---|---|---|---|---|---|---|---|---|
| | | | | | | | | | | | | | | | | | | | | | Multi-ancestry GWAS | | European GWAS | |
| | | | | | | OR | 95% CI | p-value | OR | 95% CI | p-value | Beta | SE | p-value | TAGC SNP | SNP position | $r^2$ between UKB and TAGC SNPs | Reference allele | Alternate allele | MAF | Beta [g] | p-value | Beta [h] | p-value |
| **Childhood Onset Specific** | | | | | | | | | | | | | | | | | | | | | | | | |
| **2q36·3** | rs10175070 | 2:228670575 | *CCL20, SLC18A3* | A/G | 0·251 | 1·122 | 1·086-1·16 | 4·31E-12 | 1·000 | 0·978-1·023 | 9·92E-01 | -0·862 | 0·154 | 2·24E-08 | rs13034664 [f] | 228672579 | 0·89 | A | G | 0·247 | -0·0143 | 4·27E-01 | -0·0104 | 5·78E-01 |
| **5q13·2** | rs10036789 | 5:71695918 | *PTCD2, ZNF366* | C/G | 0·460 | 1·085 | 1·054-1·117 | 4·51E-08 | 1·016 | 0·996-1·036 | 1·12E-01 | -0·511 | 0·137 | 1·95E-04 | rs10040192 [f] | 71695880 | 0·94 | T | C | 0·478 | 0·0107 | 3·87E-01 | 0·0175 | 1·74E-01 |
| **6p25·3** | rs9391997 | 6:409119 | *IRF4, DUSP22, EXOC2* | A/G | 0·527 | 1·102 | 1·070-1·135 | 6·89E-11 | 0·995 | 0·976-1·015 | 6·35E-01 | -0·597 | 0·136 | 1·06E-05 | rs9391997 | 409119 | 1·00 | A | G | 0·325 | 0·0388 | 5·11E-03 | 0·0342 | 1·47E-02 |
| **7p15·3** | rs34880821 | 7:2277545 | *IL6, TOMM7* | G/A | 0·283 | 1·106 | 1·071-1·141 | 4·73E-10 | 1·000 | 0·979-1·022 | 9·88E-01 | -0·667 | 0·149 | 7·59E-06 | N.A. | | | | | | | | | |
| **8q24·21** | rs13277355 | 8:128777719 | *MYC, TMEM75* | G/A | 0·274 | 1·110 | 1·075-1·146 | 1·65E-10 | 1·013 | 0·991-1·036 | 2·46E-01 | -0·660 | 0·151 | 1·20E-05 | rs6990534 [f] | 128814091 | 0·88 | A | G | 0·313 | -0·0351 | 6·29E-03 | -0·0301 | 3·00E-02 |
| **9q34·3** [f] | rs117137535 | 9:140500443 | *ARRDC1, ZMYND19, EHMT1* | G/A | 0·026 | 1·307 | 1·195-1·429 | 4·16E-09 | 0·977 | 0·913-1·046 | 5·10E-01 | -2·464 | 0·446 | 3·42E-08 | N.A. | | | | | | | | | |
| **10p15·1** | rs943451 | 10:6621773 | *PRKCQ, PFKFB3, SFMBT2* | C/T | 0·318 | 1·125 | 1·091-1·161 | 7·51E-14 | 1·021 | 1·000-1·043 | 5·30E-02 | -0·613 | 0·145 | 2·54E-05 | rs943451 | 6621773 | 1·00 | T | C | 0·280 | -0·0356 | 1·41E-02 | -0·0502 | 1·19E-03 |
| **11q13·1** | rs479844 | 11:65551957 | *AP5B1, OVOL1* | A/G | 0·555 | 1·106 | 1·074-1·139 | 1·48E-11 | 1·015 | 0·995-1·035 | 1·41E-01 | -0·657 | 0·135 | 1·22E-06 | rs479844 | 65551957 | 1·00 | A | G | 0·379 | 0·0532 | 2·98E-05 | 0·0546 | 2·74E-05 |
| **11q23·3** | rs12365699 | 11:118743286 | *CXCR5, DDX6* | A/G | 0·833 | 1·167 | 1·120-1·216 | 1·57E-13 | 1·004 | 0·978-1·031 | 7·43E-01 | -1·396 | 0·185 | 4·42E-14 | rs12365699 | 118743286 | 1·00 | G | A | 0·203 | -0·0494 | 4·69E-03 | -0·0430 | 1·60E-02 |
| **12q13·2** [f] | rs62623446 | 12:55368291 | *TESPA1, MUCL1, NEUROD4* | C/T | 0·072 | 1·188 | 1·127-1·252 | 1·40E-10 | 1·038 | 0·999-1·077 | 5·32E-02 | -1·079 | 0·253 | 1·95E-05 | N.A. | | | | | | | | | |
| **12q24·12** | rs10774625 | 12:111910219 | *ATXN2, SH2B3, BRAP* | A/G | 0·504 | 1·087 | 1·056-1·119 | 2·07E-08 | 0·997 | 0·978-1·017 | 7·92E-01 | -0·713 | 0·135 | 1·43E-07 | rs10774625 | 111910219 | 1·00 | A | G | 0·498 | 0·0450 | 3·52E-04 | 0·0494 | 1·20E-04 |
| **12q24·31** | rs1696361 | 12:121363823 | *SPPL3, HNF1A* | C/T | 0·359 | 1·109 | 1·076-1·142 | 1·19E-11 | 1·019 | 0·998-1·040 | 6·96E-02 | -0·696 | 0·140 | 6·84E-07 | rs558275 [f] | 121196891 | 0·87 | A | G | 0·390 | -0·0462 | 1·39E-04 | -0·0496 | 8·25E-05 |
| **17q21·2** | rs8066625 | 17:40390629 | *STAT5B, GHDC, STAT5A* | G/A | 0·100 | 1·151 | 1·098-1·206 | 4·90E-09 | 1·006 | 0·973-1·040 | 7·33E-01 | -0·846 | 0·227 | 1·88E-04 | rs4796770 [f] | 40264896 | 0·44 | C | A | 0·077 | -0·0240 | 5·20E-01 | -0·0568 | 1·82E-01 |
| **17q21·32** | rs56308324 | 17:45819206 | *TBX21, TBKBP1, OSBPL7* | A/T | 0·132 | 1·132 | 1·086-1·180 | 3·47E-09 | 1·025 | 0·996-1·055 | 8·72E-02 | -0·714 | 0·195 | 2·54E-04 | rs10514934 [f] | 45812124 | 1·00 | T | C | 0·143 | 0·0357 | 5·72E-02 | 0·0489 | 1·25E-02 |
| **18q21·33** | rs4574025 | 18:60009814 | *TNFRSF11A, KIAA1468, ZCCHC2* | C/T | 0·535 | 1·090 | 1·058-1·122 | 9·25E-09 | 0·986 | 0·967-1·006 | 1·69E-01 | -0·871 | 0·136 | 1·61E-10 | rs4369774 [f] | 60010448 | 1·00 | A | C | 0·456 | 0·0256 | 3·98E-02 | 0·0217 | 1·07E-01 |
| **18q21·33** | rs12964116 | 18:61442619 | *SERPINB7, SERPINB11, SERPINB2* | A/G | 0·036 | 1·336 | 1·247-1·432 | 2·56E-16 | 1·005 | 0·953-1·059 | 8·62E-01 | -1·916 | 0·351 | 4·87E-08 | N.A. | | | | | | | | | |
| **19p13·3** | rs4807630 | 19:1170445 | *SBNO2, GPX4, STK11* | C/T | 0·308 | 1·093 | 1·060-1·128 | 2·06E-08 | 1·013 | 0·992-1·035 | 2·32E-01 | -0·487 | 0·147 | 9·25E-04 | rs4807630 | 1170445 | 1·00 | C | T | 0·258 | 0·0498 | 2·21E-03 | 0·0447 | 1·05E-02 |
| **Adult Onset Specific** | | | | | | | | | | | | | | | | | | | | | | | | |
| **2q22·3** | rs12617922 | 2:146156679 | *TEX41, ACVR2A* | A/G | 0·518 | 0·983 | 0·955-1·012 | 2·36E-01 | 1·066 | 1·045-1·087 | 1·59E-10 | 0·478 | 0·136 | 4·46E-04 | rs6721116 [f] | 146156197 | 1·00 | G | A | 0·500 | 0·0131 | 2·68E-01 | 0·0083 | 5·14E-01 |
| **Shared** | | | | | | | | | | | | | | | | | | | | | | | | |
| **3p22·3** | rs35570272 | 3:33047662 | *GLB1, TRIM71, TMPPE* | G/T | 0·396 | 1·100 | 1·068-1·133 | 2·68E-10 | 1·046 | 1·025-1·067 | 9·84E-06 | -0·454 | 0·138 | 1·03E-03 | rs9828592 [f] | 33044339 | 0·65 | T | C | 0·418 | 0·0372 | 1·93E-03 | 0·0364 | 4·30E-03 |
| **5p15·2** | rs16903574 | 5:14610309 | *FAM105A, TRIO, OTULIN* | C/G | 0·077 | 1·236 | 1·174-1·302 | 9·01E-16 | 1·052 | 1·013-1·091 | 7·75E-03 | -1·055 | 0·250 | 2·51E-05 | N.A. | | | | | | | | | |
| **7p21·1** | rs4473914 | 7:20426263 | *ITGB6, MACC1, ABCB5* | C/T | 0·604 | 1·092 | 1·060-1·125 | 9·02E-09 | 1·035 | 1·014-1·056 | 8·44E-04 | -0·338 | 0·139 | 1·51E-02 | rs4473914 | 20426263 | 1·00 | T | C | 0·478 | -0·0118 | 3·20E-01 | -0·0155 | 2·23E-01 |
| **7p15·1** | rs917115 | 7:28172586 | *JAZF1, TAX1BP1, CREB5* | T/C | 0·208 | 1·112 | 1·074-1·152 | 1·73E-09 | 1·042 | 1·017-1·067 | 7·35E-04 | -0·577 | 0·164 | 4·17E-04 | rs917115 | 28172586 | 1·00 | T | C | 0·231 | 0·0478 | 7·08E-04 | 0·0457 | 2·21E-03 |
| **10p14** | rs7894791 | 10:8591369 | *GATA3, CELF2* | A/C | 0·587 | 1·103 | 1·071-1·137 | 9·43E-11 | 1·024 | 1·004-1·045 | 1·94E-02 | -0·494 | 0·137 | 3·24E-04 | rs7894791 | 8591369 | 1·00 | C | A | 0·352 | -0·0488 | 7·00E-05 | -0·0435 | 8·03E-04 |
| **11p15·5** | rs12788104 | 11:1123739 | *MUC6, MUC5AC* | A/G | 0·688 | 1·029 | 0·997-1·061 | 7·99E-02 | 1·064 | 1·041-1·087 | 1·41E-08 | 0·147 | 0·148 | 3·22E-01 | rs11245962 [f] | 1111164 | 0·95 | C | T (G) | 0·363 | 0·0596 [h] | 8·00E-05 | 0·0566 [i] | 7·19E-04 |
| **11q12·2** | rs174621 | 11:61630104 | *FADS2, FADS1, FADS3* | A/G | 0·772 | 1·013 | 0·978-1·049 | 4·63E-01 | 1·071 | 1·046-1·097 | 1·46E-08 | 0·508 | 0·163 | 1·87E-03 | rs174627 [f] | 61637466 | 0·62 | G | A | 0·165 | -0·0866 | 1·67E-06 | -0·0927 | 4·86E-07 |
| **12q21·1** | rs11178648 | 12:71533210 | *TSPAN8, PTPRR, LGR5* | T/C | 0·592 | 1·087 | 1·055-1·120 | 4·16E-08 | 1·052 | 1·031-1·073 | 6·62E-07 | -0·183 | 0·138 | 1·86E-01 | rs11178648 | 71533210 | 1·00 | C | T | 0·374 | -0·0454 | 1·76E-04 | -0·0429 | 6·61E-04 |
| **13q32·3** | rs1887704 | 13:99974492 | *UBAC2, DOCK9, TM9SF2* | C/G | 0·681 | 1·124 | 1·089-1·161 | 5·48E-13 | 1·048 | 1·026-1·070 | 1·57E-05 | -0·476 | 0·147 | 1·15E-03 | rs9517642 [f] | 99845912 | 0·89 | G | A | 0·264 | 0·0419 | 1·28E-03 | 0·0419 | 3·09E-03 |
| **21q22·12** | rs11088309 | 21:36464631 | *RUNX1, SETD4* | C/G | 0·143 | 1·039 | 0·997-1·083 | 6·82E-02 | 1·079 | 1·050-1·109 | 4·83E-08 | 0·063 | 0·189 | 7·38E-01 | rs17812917 [f] | 36459698 | 1·00 | G | A | 0·099 | 0·0630 | 3·57E-04 | 0·0693 | 1·20E-04 |

[a]Cytogenetic band. [b]SNP position, Genome Reference Consortium Build 37 (hg19). [c]The gene in which the SNP is located is indicated first, followed by the previous gene and the next gene; for intergenic SNPs, only the previous and next genes are shown. [d]Alleles are shown as non-risk/risk alleles, where the risk allele is the allele associated with increased asthma risk. [e]RAF, risk allele frequency in the UK Biobank. [f]The lead SNP in the UKB GWAS locus was not present and the TAGC SNP in highest LD is reported. [g]Beta values have not been adjusted for not-risk/risk allelic effects. [h]Beta value direction is discordant with UKB results. N.A. Not available; there was no SNP in the TAGC summary data with $r^2 > 0·40$

**Table 3. Summary statistics for childhood onset GWASs with and without individuals with allergic disease.** Childhood onset asthma GWAS columns are also shown in Table 2 of the manuscript.

| Locus | rsID | Position | Alleles | RAF | Childhood Onset Asthma GWAS | | | Childhood Onset Asthma GWAS (no allergies) | | |
|---|---|---|---|---|---|---|---|---|---|---|
| | | | | | OR | 95% CI | *P*-value | OR | 95% CI | *P*-value |
| 1q21.3 | rs61816761 | 1:152285861 | G/A | 0·024 | 1·970 | 1·823-2·129 | 1·88E-65 | 1·612 | 1·492-1·743 | 2·45E-19 |
| 1q25.1 | rs7518129 | 1:173163568 | A/G | 0·310 | 1·111 | 1·077-1·146 | 2·21E-11 | 1·093 | 1·06-1·128 | 4·22E-06 |
| 2q36.3 | rs10175070 | 2:228670575 | A/G | 0·251 | 1·122 | 1·086-1·160 | 4·31E-12 | 1·141 | 1·104-1·179 | 1·00E-10 |
| 3q28 | rs12634152 | 3:188121019 | C/T | 0·547 | 1·133 | 1·100-1·166 | 1·08E-16 | 1·121 | 1·089-1·155 | 4·33E-10 |
| 4p14 | rs5743618 | 4:38798648 | A/C | 0·774 | 1·249 | 1·203-1·295 | 3·19E-32 | 1·252 | 1·207-1·299 | 1·67E-22 |
| 5q31.1 | rs2051809 | 5:132056874 | C/A | 0·247 | 1·175 | 1·137-1·214 | 3·24E-22 | 1·182 | 1·144-1·221 | 1·92E-16 |
| 5q13.2 | rs10036789 | 5:71695918 | C/G | 0·460 | 1·085 | 1·054-1·117 | 4·51E-08 | 1·093 | 1·061-1·125 | 1·23E-06 |
| 6p25.3 | rs9391997 | 6:409119 | A/G | 0·527 | 1·102 | 1·070-1·135 | 6·89E-11 | 1·092 | 1·06-1·124 | 1·68E-06 |
| 7p15.3 | rs34880821 | 7:2277545 | G/A | 0·283 | 1·106 | 1·071-1·141 | 4·73E-10 | 1·092 | 1·058-1·128 | 8·39E-06 |
| 8q24.21 | rs13277355 | 8:128777719 | G/A | 0·274 | 1·110 | 1·075-1·146 | 1·65E-10 | 1·112 | 1·077-1·148 | 1·10E-07 |
| 9q34.3 | rs117137535 | 9:140500443 | G/A | 0·026 | 1·307 | 1·195-1·429 | 4·16E-09 | 1·253 | 1·146-1·37 | 7·44E-05 |
| 10p15.1 | rs943451 | 10:6621773 | C/T | 0·318 | 1·125 | 1·091-1·161 | 7·51E-14 | 1·113 | 1·079-1·148 | 3·36E-08 |
| 11q23.3 | rs12365699 | 11:118743286 | A/G | 0·833 | 1·167 | 1·120-1·216 | 1·57E-13 | 1·151 | 1·104-1·199 | 3·83E-08 |
| 12q13.2 | rs62623446 | 12:55368291 | C/T | 0·072 | 1·188 | 1·127-1·252 | 1·40E-10 | 1·156 | 1·097-1·219 | 1·24E-05 |
| 12q24.12 | rs10774625 | 12:111910219 | A/G | 0·504 | 1·087 | 1·056-1·119 | 2·07E-08 | 1·099 | 1·068-1·132 | 1·84E-07 |
| 12q24.31 | rs1696361 | 12:121363823 | C/T | 0·359 | 1·109 | 1·076-1·142 | 1·19E-11 | 1·112 | 1·079-1·145 | 1·50E-08 |
| 17q12 | rs4795399 | 17:38061439 | C/T | 0·529 | 1·406 | 1·365-1·448 | 1·45E-111 | 1·456 | 1·414-1·5 | 5·64E-90 |
| 17q21.2 | rs8066625 | 17:40390629 | G/A | 0·100 | 1·151 | 1·098-1·206 | 4·90E-09 | 1·157 | 1·104-1·213 | 6·82E-07 |
| 17q21.32 | rs56308324 | 17:45819206 | A/T | 0·132 | 1·132 | 1·086-1·180 | 3·47E-09 | 1·121 | 1·075-1·168 | 1·03E-05 |
| 18q21.33 | rs4574025 | 18:60009814 | C/T | 0·535 | 1·090 | 1·058-1·122 | 9·25E-09 | 1·092 | 1·061-1·125 | 1·46E-06 |
| 18q21.33 | rs12964116 | 18:61442619 | A/G | 0·036 | 1·336 | 1·247-1·432 | 2·56E-16 | 1·357 | 1·266-1·455 | 1·16E-12 |
| 19p13.3 | rs4807630 | 19:1170445 | C/T | 0·308 | 1·093 | 1·060-1·128 | 2·06E-08 | 1·079 | 1·046-1·113 | 9·83E-05 |
| 2q22.3 | rs12617922 | 2:146156679 | A/G | 0·518 | 0·983 | 0·955-1·012 | 2·36E-01 | 1·003 | 0·974-1·032 | 8·86E-01 |
| 1q32.1 | rs12023876 | 1:203093201 | T/G | 0·668 | 1·102 | 1·068-1·137 | 1·11E-09 | 1·100 | 1·066-1·135 | 1·05E-06 |
| 2p25.1 | rs13416555 | 2:8441735 | G/C | 0·705 | 1·130 | 1·094-1·167 | 1·94E-13 | 1·129 | 1·092-1·166 | 2·72E-09 |
| 2q12.1 | rs72823641 | 2:102936159 | A/T | 0·863 | 1·415 | 1·349-1·484 | 3·01E-46 | 1·430 | 1·364-1·5 | 3·33E-33 |
| 2q37.3 | rs34290285 | 2:242698640 | A/G | 0·745 | 1·177 | 1·137-1·219 | 2·06E-20 | 1·191 | 1·151-1·233 | 5·65E-16 |
| 3p22.3 | rs35570272 | 3:33047662 | G/T | 0·396 | 1·100 | 1·068-1·133 | 2·68E-10 | 1·089 | 1·058-1·122 | 3·86E-06 |
| 4q27 | rs2069763 | 4:123377482 | C/A | 0·334 | 1·126 | 1·092-1·160 | 1·59E-14 | 1·130 | 1·097-1·165 | 9·67E-11 |
| 5p15.2 | rs16903574 | 5:14610309 | C/G | 0·077 | 1·236 | 1·174-1·302 | 9·01E-16 | 1·188 | 1·128-1·251 | 1·52E-07 |
| 5q22.1 | rs1837253 | 5:110401872 | T/C | 0·740 | 1·211 | 1·169-1·253 | 2·33E-27 | 1·211 | 1·17-1·254 | 7·01E-19 |
| 5q31.1 | rs17622378 | 5:131778452 | G/A | 0·573 | 1·103 | 1·071-1·136 | 8·52E-11 | 1·100 | 1·068-1·133 | 2·77E-07 |
| 6p22.1 | rs1117490 | 6:30170510 | T/C | 0·234 | 1·100 | 1·063-1·137 | 2·78E-08 | 1·093 | 1·057-1·131 | 2·15E-05 |
| 6p21.33 | rs2428494 | 6:31322197 | T/A | 0·477 | 1·157 | 1·124-1·191 | 7·94E-23 | 1·160 | 1·127-1·194 | 3·59E-16 |
| 6p21.32 | rs28407950 | 6:32626348 | T/C | 0·756 | 1·354 | 1·306-1·405 | 1·27E-59 | 1·359 | 1·31-1·409 | 5·11E-41 |
| 6q15 | rs1321859 | 6:91011673 | T/C | 0·649 | 1·102 | 1·068-1·136 | 9·32E-10 | 1·099 | 1·066-1·134 | 9·91E-07 |
| 7p21.1 | rs4473914 | 7:20426263 | C/T | 0·604 | 1·092 | 1·060-1·125 | 9·02E-09 | 1·101 | 1·068-1·134 | 3·09E-07 |
| 7p15.1 | rs917115 | 7:28172586 | T/C | 0·208 | 1·112 | 1·074-1·152 | 1·73E-09 | 1·119 | 1·081-1·158 | 2·14E-07 |
| 8q21.13 | rs4739738 | 8:81291645 | A/G | 0·359 | 1·117 | 1·084-1·151 | 3·45E-13 | 1·113 | 1·08-1·147 | 1·02E-08 |
| 9p24.1 | rs992969 | 9:6209697 | G/A | 0·252 | 1·248 | 1·209-1·289 | 6·76E-42 | 1·268 | 1·228-1·309 | 1·46E-32 |
| 10p14 | rs7894791 | 10:8591369 | A/C | 0·587 | 1·103 | 1·071-1·137 | 9·43E-11 | 1·117 | 1·084-1·151 | 2·76E-09 |
| 10p14 | rs1775554 | 10:9054340 | C/A | 0·577 | 1·117 | 1·084-1·150 | 2·85E-13 | 1·130 | 1·097-1·164 | 4·83E-11 |
| 11p15.5 | rs12788104 | 11:1123739 | A/G | 0·688 | 1·029 | 0·997-1·061 | 7·99E-02 | 1·036 | 1·004-1·069 | 7·34E-02 |
| 11q12.2 | rs174621 | 11:61630104 | A/G | 0·772 | 1·013 | 0·978-1·049 | 4·63E-01 | 0·999 | 0·965-1·035 | 9·80E-01 |
| 11q13.1 | rs479844 | 11:65551957 | A/G | 0·555 | 1·106 | 1·074-1·139 | 1·48E-11 | 1·087 | 1·056-1·12 | 4·95E-06 |
| 11q13.5 | rs61894547 | 11:76248630 | C/T | 0·052 | 1·463 | 1·382-1·548 | 2·17E-39 | 1·436 | 1·357-1·52 | 6·29E-24 |
| 11q13.5 | rs7936312 | 11:76293726 | G/T | 0·477 | 1·194 | 1·160-1·230 | 4·89E-33 | 1·189 | 1·155-1·224 | 1·61E-21 |

| | | | | | | | | | | |
|---|---|---|---|---|---|---|---|---|---|---|
| 12q13.11 | rs56389811 | 12:48205358 | T/C | 0·761 | 1·022 | 0·988-1·058 | 2·11E-01 | 1·022 | 0·987-1·057 | 3·18E-01 |
| 12q13.2 | rs705699 | 12:56384804 | G/A | 0·425 | 1·106 | 1·074-1·139 | 1·43E-11 | 1·087 | 1·055-1·119 | 5·53E-06 |
| 12q13.3 | rs3122929 | 12:57509102 | C/T | 0·404 | 1·134 | 1·101-1·167 | 6·59E-17 | 1·155 | 1·121-1·189 | 5·34E-15 |
| 12q21.1 | rs11178648 | 12:71533210 | T/C | 0·592 | 1·087 | 1·055-1·120 | 4·16E-08 | 1·086 | 1·054-1·119 | 8·63E-06 |
| 13q32.3 | rs1887704 | 13:99974492 | C/G | 0·681 | 1·124 | 1·089-1·161 | 5·48E-13 | 1·121 | 1·086-1·157 | 1·02E-08 |
| 14q24.1 | rs1950897 | 14:68760141 | T/C | 0·287 | 1·096 | 1·062-1·131 | 1·38E-08 | 1·093 | 1·059-1·128 | 7·44E-06 |
| 15q22.2 | rs11071559 | 15:61069988 | T/C | 0·872 | 1·205 | 1·151-1·262 | 2·57E-15 | 1·215 | 1·16-1·273 | 1·66E-11 |

**Table 4. Results of GWAS-PW to test for co-localization of variants associated with childhood onset and adult onset asthma.** The childhood onset GWAS results were input for phenotype 1 and the adult onset GWAS results were input for phenotype 2. The childhood onset specific loci and adult onset specific locus did not colocalize (high probabilities of P1 for childhood onset, low probabilities of P2 for adult onset, and zero to very low probabilities of P3 for colocalization). The only exceptions are the childhood onset loci at 5q31.1, which has a P1 = 0 and a P4 = 1, and at 17q21.32, which has a P1 = 0.05 and a P4 = 0.95. P4 indicates that the locus is shared but different variants are associated with risk in the childhood onset and adult onset GWASs. This is evident in the locus zoom plots of these two regions (appendix pages 23 and 34, respectively) where it is clear that the variants that are genome-wide significant in the childhood onset GWAS (over *KIF3A* at the 5q31.1 locus and over *TBX21* at the 17q21.32 locus) are not significant in the adult onset GWAS (lead SNP shown as a blue diamond and SNPs in high LD with the lead SNPs shown as red dots). At the 17q21.32 locus, the lead genome-wide significant variant in the childhood onset GWAS had a p=0.08 in the adult onset GWAS, although other variants in this region that were not in LD with the lead childhood onset variant had p-values ~$10^{-6}$ in the adult onset GWAS. Because those variants did not reach genome-wide significance, a second 17q21.32 locus was not included as a shared or adult onset specific locus in our analysis. In contrast, the 5q31.1 locus was listed as both childhood onset specific and shared because a second peak over *IRF1* was significant in both GWAS. Among the shared loci, nearly all have high probabilities for P3 (indicating co-localization) except for a few, such as 5q31.1 that have low probabilities of P3 and high probabilities of P4 as discussed above, or have similar P3 and P4 probabilities. This indicates that these are complex regions that likely harbor multiple independent susceptibility variants, not all of which reached genome wide significance in our GWASs. Notably, we do not find high probabilities of P1 or P2 for the shared loci, indicating that the shared loci are not misclassified.

| Locus | rsID | Position | P1 | P2 | P3 | P4 |
|---|---|---|---|---|---|---|
| **Childhood onset loci** | | | | | | |
| 1q21.3 | rs61816761 | 1:152285861 | 0.954 | 0.000 | 0.000 | 0.046 |
| 1q25.1 | rs7518129 | 1:173163568 | 0.961 | 0.000 | 0.000 | 0.039 |
| 2q36.3 | rs10175070 | 2:228670575 | 0.941 | 0.000 | 0.000 | 0.059 |
| 3q28 | rs12634152 | 3:188121019 | 0.951 | 0.000 | 0.000 | 0.048 |
| 4p14 | rs5743618 | 4:38798648 | 0.965 | 0.000 | 0.000 | 0.035 |
| 5q13.2 | rs10036789 | 5:71695918 | 0.954 | 0.000 | 0.000 | 0.045 |
| 5q31.1 | rs2051809 | 5:132056874 | 0.000 | 0.000 | 0.000 | 1.000 |
| 6p25.3 | rs9391997 | 6:409119 | 0.974 | 0.000 | 0.000 | 0.026 |
| 7p15.3 | rs34880821 | 7:22775450 | 0.958 | 0.000 | 0.000 | 0.042 |
| 8q24.21 | rs13277355 | 8:128777719 | 0.898 | 0.000 | 0.000 | 0.102 |
| 9q34.3 | rs117137535 | 9:140500443 | 1.000 | 0.000 | 0.000 | 0.000 |
| 10p15.1 | rs943451 | 10:6621773 | 0.939 | 0.000 | 0.001 | 0.060 |
| 11q13.1 | rs479844 | 11:65551957 | 0.972 | 0.000 | 0.000 | 0.028 |
| 11q23.3 | rs12365699 | 11:118743286 | 0.959 | 0.000 | 0.000 | 0.041 |
| 12q13.2 | rs62623446 | 12:55368291 | 0.963 | 0.000 | 0.001 | 0.036 |
| 12q24.12 | rs10774625 | 12:111910219 | 0.957 | 0.000 | 0.000 | 0.042 |
| 12q24.31 | rs1696361 | 12:121363823 | 0.956 | 0.000 | 0.000 | 0.044 |
| 17q12 | rs4795399 | 17:38061439 | 0.929 | 0.000 | 0.000 | 0.071 |
| 17q21.2 | rs8066625 | 17:40390629 | 0.920 | 0.000 | 0.000 | 0.080 |
| 17q21.32 | rs56308324 | 17:45819206 | 0.051 | 0.000 | 0.000 | 0.949 |
| 18q21.33 | rs4574025 | 18:60009814 | 0.951 | 0.000 | 0.000 | 0.049 |
| 18q21.33 | rs12964116 | 18:61442619 | 0.953 | 0.000 | 0.000 | 0.047 |
| 19p13.3 | rs4807630 | 19:1170445 | 0.964 | 0.000 | 0.000 | 0.036 |
| **Adult onset locus** | | | | | | |
| 2q22.3 | rs12617922 | 2:146156679 | 0.000 | 0.984 | 0.003 | 0.013 |
| **Shared loci** | | | | | | |
| 1q32.1 | rs12023876 | 1:203093201 | 0.000 | 0.000 | 0.938 | 0.062 |
| 2p25.1 | rs13416555 | 2:8441735 | 0.000 | 0.000 | 0.988 | 0.012 |
| 2q12.1 | rs72823641 | 2:102936159 | 0.000 | 0.000 | 0.183 | 0.817 |
| 2q37.3 | rs34290285 | 2:242698640 | 0.000 | 0.000 | 0.994 | 0.006 |
| 3p22.3 | rs35570272 | 3:33047662 | 0.000 | 0.000 | 0.999 | 0.001 |
| 4q27 | rs2069763 | 4:123377482 | 0.000 | 0.000 | 0.648 | 0.352 |

| | | | | | | |
|---|---|---|---|---|---|---|
| 5p15.2 | rs16903574 | 5:14610309 | 0.000 | 0.000 | 0.961 | 0.038 |
| 5q22.1 | rs1837253 | 5:110401872 | 0.000 | 0.000 | 1.000 | 0.000 |
| 5q31.1 | rs17622378 | 5:131778452 | 0.000 | 0.000 | 0.000 | 1.000 |
| 6p22.1 | rs1117490 | 6:30170510 | 0.000 | 0.000 | 0.995 | 0.005 |
| 6p21.33 | rs2428494 | 6:31322197 | 0.000 | 0.000 | 1.000 | 0.000 |
| 6p21.32 | rs28407950 | 6:32626348 | 0.000 | 0.000 | 0.000 | 1.000 |
| 6q15 | rs1321859 | 6:91011673 | 0.000 | 0.000 | 0.978 | 0.022 |
| 7p21.1 | rs4473914 | 7:20426263 | 0.000 | 0.000 | 0.952 | 0.048 |
| 7p15.1 | rs917115 | 7:28172586 | 0.000 | 0.000 | 0.965 | 0.034 |
| 8q21.13 | rs4739738 | 8:81291645 | 0.000 | 0.000 | 0.964 | 0.036 |
| 9p24.1 | rs992969 | 9:6209697 | 0.000 | 0.000 | 0.958 | 0.042 |
| 10p14 | rs7894791 | 10:8591369 | 0.000 | 0.000 | 0.602 | 0.398 |
| 10p14 | rs1775554 | 10:9054340 | 0.000 | 0.000 | 0.879 | 0.121 |
| 11p15.5 | rs12788104 | 11:1123739 | 0.000 | 0.001 | 0.441 | 0.588 |
| 11q12.2 | rs174621 | 11:61630104 | 0.000 | 0.003 | 0.659 | 0.338 |
| 11q13.5 | rs61894547 | 11:76248630 | 0.000 | 0.000 | 0.984 | 0.016 |
| 11q13.5 | rs7936312 | 11:76293726 | 0.000 | 0.000 | 0.063 | 0.937 |
| 12q13.11 | rs56389811 | 12:48205358 | 0.000 | 0.001 | 0.275 | 0.724 |
| 12q13.2 | rs705699 | 12:56384804 | 0.000 | 0.000 | 0.973 | 0.027 |
| 12q13.3 | rs3122929 | 12:57509102 | 0.000 | 0.000 | 0.880 | 0.120 |
| 12q21.1 | rs11178648 | 12:71533210 | 0.000 | 0.000 | 0.969 | 0.031 |
| 13q32.3 | rs1887704 | 13:99974492 | 0.000 | 0.000 | 0.987 | 0.013 |
| 14q24.1 | rs1950897 | 14:68760141 | 0.000 | 0.000 | 0.918 | 0.082 |
| 15q22.2 | rs11071559 | 15:61069988 | 0.000 | 0.000 | 0.992 | 0.008 |
| 15q22.33 | rs56062135 | 15:67455630 | 0.000 | 0.000 | 0.992 | 0.008 |
| 16p13.13 | rs35032408 | 16:11215424 | 0.000 | 0.000 | 0.997 | 0.003 |
| 16p12.1 | rs3785356 | 16:27349168 | 0.000 | 0.000 | 0.090 | 0.910 |
| 16q12.1 | rs2066844 | 16:50745926 | 0.001 | 0.000 | 0.925 | 0.075 |
| 17q21.33 | rs28406364 | 17:47454507 | 0.000 | 0.000 | 0.982 | 0.018 |
| 19q13.11 | rs10414065 | 19:33721455 | 0.000 | 0.000 | 0.981 | 0.019 |
| 21q22.12 | rs11088309 | 21:36464631 | 0.000 | 0.001 | 0.831 | 0.168 |

**Table 5. Summary statistics for the age of asthma onset GWAS and the childhood and adult onset GWAS.** Information for the most significant SNP at each of 19 independent loci in the age of onset GWAS is shown.

| Locus | rsID | Position | Alleles | MAF | Age of Asthma Onset GWAS | | | Childhood Onset Asthma GWAS | | | Adult Onset Asthma GWAS | | |
|---|---|---|---|---|---|---|---|---|---|---|---|---|---|
| | | | | | Beta | SE | p-value | OR | 95% CI | p-value | OR | 95% CI | p-value |
| 1q21.3 | rs61816761 | rs61816761 | G/A | 0·024 | -4·571 | 0·426 | 8·145E-27 | 1·970 | 1·823-2·129 | 1·879E-65 | 1·016 | 0·948-1·088 | 6·563E-01 |
| 1q25.1 | rs7518129 | rs7518129 | A/G | 0·310 | -0·849 | 0·145 | 4·887E-09 | 1·111 | 1·077-1·146 | 2·213E-11 | 0·997 | 0·977-1·019 | 8·168E-01 |
| 2q12.1 | rs3771175 | rs3771175 | A/T | 0·863 | -1·726 | 0·207 | 7·661E-17 | 1·410 | 1·344-1·478 | 1·529E-45 | 1·104 | 1·072-1·137 | 2·810E-11 |
| 2q36.3 | rs10187276 | rs10187276 | C/T | 0·251 | -0·865 | 0·154 | 1·976E-08 | 1·122 | 1·086-1·159 | 4·543E-12 | 1·000 | 0·977-1·023 | 9·853E-01 |
| 2q37.3 | rs78147778 | rs78147778 | C/T | 0·748 | -0·914 | 0·162 | 1·638E-08 | 1·168 | 1·128-1·210 | 3·464E-18 | 1·070 | 1·045-1·095 | 9·790E-09 |
| 3q28 | rs2889896 | rs2889896 | T/C | 0·547 | -0·975 | 0·136 | 8·066E-13 | 1·129 | 1·096-1·162 | 6·485E-16 | 1·007 | 0·987-1·027 | 5·076E-01 |
| 4p14 | rs5743618 | rs5743618 | A/C | 0·774 | -1·578 | 0·163 | 4·526E-22 | 1·249 | 1·203-1·295 | 3·188E-32 | 1·008 | 0·985-1·032 | 5·090E-01 |
| 5q31.1 | rs4705962 | rs4705962 | C/T | 0·229 | -0·985 | 0·159 | 5·571E-10 | 1·170 | 1·132-1·210 | 2·238E-20 | 1·010 | 0·987-1·034 | 3·814E-01 |
| 6p21.33 | rs12207974 | rs12207974 | G/C | 0·776 | -1·066 | 0·164 | 8·862E-11 | 1·100 | 1·062-1·141 | 1·614E-07 | 0·978 | 0·956-1·002 | 6·758E-02 |
| 6p21.33 | rs1093 | rs1093 | A/G | 0·212 | -0·994 | 0·162 | 8·405E-10 | 1·138 | 1·100-1·178 | 1·459E-13 | 1·036 | 1·012-1·061 | 3·152E-03 |
| 6p21.32 | rs9274659 | rs9274659 | T/A | 0·670 | -1·249 | 0·145 | 6·707E-18 | 1·175 | 1·138-1·213 | 2·512E-23 | 1·004 | 0·983-1·025 | 7·226E-01 |
| 9p24.1 | rs7848215 | rs7848215 | C/T | 0·261 | -1·031 | 0·151 | 7·510E-12 | 1·246 | 1·207-1·287 | 7·805E-42 | 1·098 | 1·074-1·122 | 1·157E-16 |
| 9q34.3 | rs117137535 | rs117137535 | G/A | 0·026 | -2·464 | 0·446 | 3·419E-08 | 1·307 | 1·195-1·429 | 4·160E-09 | 0·977 | 0·913-1·046 | 5·097E-01 |
| 11q13.5 | rs61894547 | rs61894547 | C/T | 0·052 | -2·227 | 0·284 | 4·421E-15 | 1·463 | 1·382-1·548 | 2·167E-39 | 1·093 | 1·047-1·141 | 5·462E-05 |
| 11q23.3 | rs12365699 | rs12365699 | A/G | 0·833 | -1·396 | 0·185 | 4·423E-14 | 1·167 | 1·120-1·216 | 1·575E-13 | 1·004 | 0·978-1·031 | 7·434E-01 |
| 17q12 | rs4795399 | rs4795399 | C/T | 0·529 | -2·286 | 0·134 | 6·760E-65 | 1·406 | 1·365-1·448 | 1·448E-111 | 1·005 | 0·985-1·025 | 6·279E-01 |
| 17q21.2 | rs11658582 | rs11658582 | C/G | 0·626 | -0·905 | 0·141 | 1·373E-10 | 1·106 | 1·073-1·140 | 7·912E-11 | 1·012 | 0·992-1·033 | 2·358E-01 |
| 18q21.33 | rs4574025 | rs4574025 | C/T | 0·535 | -0·871 | 0·136 | 1·605E-10 | 1·090 | 1·058-1·122 | 9·251E-09 | 0·986 | 0·967-1·006 | 1·690E-01 |
| 18q21.33 | rs12964116 | rs12964116 | A/G | 0·036 | -1·916 | 0·351 | 4·866E-08 | 1·336 | 1·247-1·432 | 2·561E-16 | 1·005 | 0·953-1·059 | 8·623E-01 |

**Table 6. Asthma age of onset GWAS comparison.** The lead SNPs reported by Sarnowski et al[7] are shown with their p values in the three UKB asthma GWASs.

| Sarnowski et al | | | | | | UKB | | |
|---|---|---|---|---|---|---|---|---|
| **Reported SNP** | **Locus** | **Effect/reference alleles** | **Effect frequency** | **Hazard ratio** | **P value** | **Childhood onset GWAS p value** | **Adult onset GWAS p value** | **Age of onset GWAS p value** |
| GWS | | | | | | | | |
| rs10208293 | 2q12 | G/A | 0·73 | 1·14 | 3·10E-08 | 3·68E-38 | 3·11E-06 | 1·36E-15 |
| rs9272346 | 6p21 | A/G | 0·59 | 1·13 | 1·60E-08 | 4·17E-37 | 2·48E-47 | 7·18E-08 |
| rs928413 | 9p24 | G/A | 0·25 | 1·19 | 6·50E-16 | 2·01E-40 | 4·61E-17 | 5·13E-11 |
| rs1861760 | 16q22 | A/C | 0·04 | 1·28 | 4·20E-08 | 5·68E-01 | 2·96E-01 | 8·80E-01 |
| rs9901146 | 17q12-q21 | G/A | 0·51 | 1·18 | 1·90E-16 | 8·14E-99 | 4·43E-01 | 4·46E-57 |
| Suggestive loci | | | | | | | | |
| rs12468899 | 2q11-q12 | G/A | 0·69 | 1·12 | 1·70E-07 | 4·45E-03 | 8·83E-01 | 4·08E-02 |
| rs11071559 | 15q22 | C/T | 0·85 | 1·16 | 8·30E-07 | 2·57E-15 | 1·22E-05 | 5·11E-06 |
| rs1805013 | 16p12-p11 | T/C | 0·05 | 1·22 | 8·00E-07 | 9·62E-01 | 3·76E-01 | 5·75E-01 |

**Table 7. Tissue enrichment p-values of GWAS loci.** Bonferroni adjusted 1-sided p-values for higher expression from FUMA[5] are shown.

| | Childhood onset p-value | Adult onset p-value |
|---|---|---|
| **Skin** | 5·72E-03 | 1·00E+00 |
| **Lung** | 9·79E-01 | 6·39E-03 |
| **Blood** | 9·13E-03 | 4·00E-04 |
| **Spleen** | 1·22E-01 | 2·38E-03 |
| **Small intestine** | 2·03E-02 | 4·87E-03 |

**Table 8. Results of PrediXcan analysis in 5 tissues.** Results sorted by locus category and chromosomal band within each category. Only predicted genes with significant associations with asthma ($<1{\cdot}4\times10^{-6}$) in at least one tissue are shown. Gene categories: A, adult onset; C, childhood onset; S, shared (see above Methods for details).

| Locus Category | Gene | Chr. Band (Gene) | Chr. Band (Locus) | Gene Category | Childhood Onset - P-values | | | | | | Childhood Onset - Z-scores | | | | | | Childhood Onset - Betas | | | | | | Adult Onset - P-values | | | | | | Adult Onset - Z-scores | | | | | | Adult Onset - Betas | | | | | |
|---|---|---|---|---|---|---|---|---|---|---|---|---|---|---|---|---|---|---|---|---|---|---|---|---|---|---|---|---|---|---|---|---|---|---|---|---|---|---|---|---|
| | | | | | Lung | Skin (not sun exposed-suprapubic) | Skin (sun exposed-lower leg) | Small Intestine (terminal ilium) | Spleen | Whole blood | Lung | Skin (not sun exposed-suprapubic) | Skin (sun exposed-lower leg) | Small Intestine (terminal ilium) | Spleen | Whole blood | Lung | Skin (not sun exposed-suprapubic) | Skin (sun exposed-lower leg) | Small Intestine (terminal ilium) | Spleen | Whole blood | Lung | Skin (not sun exposed-suprapubic) | Skin (sun exposed-lower leg) | Small Intestine (terminal ilium) | Spleen | Whole blood | Lung | Skin (not sun exposed-suprapubic) | Skin (sun exposed-lower leg) | Small Intestine (terminal ilium) | Spleen | Whole blood | Lung | Skin (not sun exposed-suprapubic) | Skin (sun exposed-lower leg) | Small Intestine (terminal ilium) | Spleen | Whole blood |
| Childhood | CRCT1 | 1q21.3 | 1q21.3 | C | | 2.8E-11 | | | | | | 6.65592 | | | | | | 0.31804 | | | | | | 0.87436 | | | | | | 0.15812 | | | | | | 0.0053 | | | | |
| Childhood | CRNN | 1q21.3 | 1q21.3 | C | | 4.1E-09 | 0.00262 | | | | | 5.88137 | 3.00967 | | | | | 0.23988 | 0.07501 | | | | | 0.57488 | 0.69656 | | | | | 0.56088 | -0.39 | | | | | 0.0175 | -0.0047 | | | |
| Childhood | FLG | 1q21.3 | 1q21.3 | C | 8.5E-14 | 1.7E-16 | 0.0723 | | 2.4E-10 | | -7.4618 | -8.2421 | -1.7972 | | -6.3302 | | -0.2152 | -0.43 | -0.0828 | | -0.2829 | | 0.1837 | 0.2094 | 0.18283 | | 0.77249 | | -1.3294 | 1.25522 | 1.33208 | | -0.2891 | | -0.0256 | 0.04843 | 0.0595 | | -0.0087 | |
| Childhood | LINGO4 | 1q21.3 | 1q21.3 | C | 3.2E-12 | 0.00013 | 1.9E-08 | | | 3.3E-27 | 6.96918 | 3.83579 | 5.62437 | | | 10.805 | 0.23411 | 0.09659 | 0.08699 | | | 0.58797 | 0.94526 | 0.47137 | 0.39294 | | | 0.66914 | -0.0687 | -0.7202 | -0.8543 | | | 0.42733 | -0.003 | -0.0123 | -0.011 | | | 0.0165 |
| Childhood | PSMD4 | 1q21.3 | 1q21.3 | C | 7.5E-08 | | | | | 0.12948 | 5.37817 | | | | | 1.51616 | 0.29395 | | | | | 0.12555 | 0.13637 | | | | | 0.19887 | 1.48943 | | | | | 1.28477 | 0.05371 | | | | | 0.07321 |
| Childhood | S100A12 | 1q21.3 | 1q21.3 | C | 8.6E-07 | | | | | 0.785 | -4.9221 | | | | | -0.2728 | -0.2387 | | | | | -0.0075 | 0.0847 | | | | | 0.80808 | -1.724 | | | | | -0.2429 | -0.0599 | | | | | -0.0049 |
| Childhood | SPRR2D | 1q21.3 | 1q21.3 | C | | 2.2E-08 | 0.0002 | | 0.25746 | | | -5.5977 | -3.7131 | | -1.1324 | | | -0.3674 | -0.1215 | | -0.0682 | | | 0.18306 | 0.84034 | | 0.96243 | | | -1.3314 | -0.2015 | | -0.0471 | | | -0.0595 | -0.0041 | | 0.0001 | |
| Childhood | TDRKH | 1q21.3 | 1q21.3 | C | 5E-09 | 1.3E-08 | 1.3E-07 | | 6.8E-09 | 1.5E-07 | -5.8487 | -5.6814 | -5.2802 | | -5.7965 | -5.2469 | -0.2712 | -0.2909 | -0.2428 | | -0.1861 | -0.2874 | 0.38593 | 0.0116 | 0.10471 | | 0.36571 | 0.13753 | -0.867 | -2.524 | -1.6224 | | -0.9045 | -1.485 | -0.0271 | -0.0869 | -0.05 | | -0.0194 | -0.0545 |
| Childhood | THEM5 | 1q21.3 | 1q21.3 | C | | 8.6E-08 | 0.28529 | | | | | 5.35473 | -1.0685 | | | | | 0.45414 | -0.0816 | | | | | 0.31027 | 0.20794 | | | | | 1.01466 | 1.25926 | | | | | 0.06018 | 0.06266 | | | |
| Childhood | CCL20 | 2q36.3 | 2q36.3 | C | | | | | | 3.4E-11 | | | | | | -6.6269 | | | | | | -0.3057 | | | | | | 0.50143 | | | | | | -0.6722 | | | | | | -0.0211 |
| Childhood | FAM114A1 | 4p14 | 4p14 | C | 7.4E-11 | 0.03577 | 0.02561 | 0.8541 | 0.04131 | | -6.5119 | -2.0995 | -2.2321 | -0.1839 | -2.0404 | | -0.2944 | -0.0541 | -0.0449 | -0.0183 | -0.0415 | | 0.8888 | 0.80663 | 0.52889 | 0.6266 | 0.48267 | | 0.13982 | 0.24477 | 0.6297 | 0.48651 | 0.70201 | | 0.00589 | 0.00446 | 0.00852 | 0.03928 | 0.01019 | |
| Childhood | TLR10 | 4p14 | 4p14 | C | | | | | | 9.4E-11 | | | | | | 6.47706 | | | | | | 0.26329 | | | | | | 0.76021 | | | | | | 0.30521 | | | | | | 0.00536 |
| Childhood | TLR6 | 4p14 | 4p14 | C | | 1.7E-12 | 0.00413 | | | | | 7.0589 | 2.86775 | | | | | 0.24776 | 0.09865 | | | | | 0.43459 | 0.65707 | | | | | 0.78136 | 0.44396 | | | | | 0.01725 | 0.01004 | | | |
| Childhood | STEAP1B | 7p15.3 | 7p15.3 | C | 1.3E-07 | | 0.0001 | | | | -5.2848 | | -3.8814 | | | | -0.2646 | | -0.32 | | | | 0.10445 | | 0.64613 | | | | -1.6237 | | -0.4591 | | | | -0.0542 | | -0.0259 | | | |
| Shared | AP5B1 | 11q13.1 | 11q13.1 | C | 6E-08 | | 0.62619 | | | | -5.4192 | | -0.4871 | | | | -0.2151 | | -0.0356 | | | | 0.03659 | | 0.99713 | | | | -2.0903 | | 0.0036 | | | | -0.0567 | | -0.0008 | | | |
| Childhood | MLEC | 12q24.31 | 12q24.31 | C | | 2.4E-06 | 3.5E-08 | | | 0.0555 | | 4.71449 | 5.51362 | | | 1.91495 | | 0.11842 | 0.16299 | | | 0.12957 | | 0.20498 | 0.11452 | | | 0.23374 | | 1.26748 | 1.5782 | | | -1.1908 | | 0.02091 | 0.03076 | | | -0.0504 |
| Childhood | PSMD9 | 12q24.31 | 12q24.31 | C | | | 0.53502 | | 4.6E-07 | | | | -0.6204 | | 5.04025 | | | | -0.051 | | 0.17301 | | | | 0.52571 | | 0.29009 | | | | -0.6346 | | 1.05792 | | | | -0.0346 | | 0.02208 | |
| Childhood | SPPL3 | 12q24.31 | 12q24.31 | C | 2.1E-07 | | | 4.2E-09 | | 0.02285 | 5.18955 | | | 5.87614 | | 2.27588 | 0.21651 | | | 0.31682 | | 0.09334 | 0.10213 | | | 0.05378 | | 0.81007 | 1.63461 | | | 1.92859 | | -0.2403 | 0.04579 | | | 0.0704 | | -0.0107 |
| Childhood | AC087491.2 | 17q12 | 17q12 | C | | 4.2E-23 | | | | | | -9.8985 | | | | | | -0.4606 | | | | | | 0.4942 | | | | | | 0.68364 | | | | | | 0.02383 | | | | |
| Childhood | CSF3 | 17q21.1 | 17q12 | C | 0.30861 | | | | 1.2E-10 | | -1.0181 | | | | 6.44543 | | -0.1429 | | | | 0.18669 | | 0.36395 | | | | 0.98288 | | -0.9079 | | | | 0.02146 | | -0.0845 | | | | -0.0004 | |
| Childhood | ERBB2 | 17q12 | 17q12 | C | | 2E-49 | 4.3E-47 | | | | | 14.7784 | 14.4131 | | | | | 0.99477 | 0.83492 | | | | | 0.32963 | 0.37545 | | | | | 0.97487 | 0.88632 | | | | | 0.04851 | 0.03623 | | | |
| Childhood | FBXL20 | 17q12 | 17q12 | C | | | | 3.2E-15 | | | | | | -7.8845 | | | | | | -0.2834 | | | | | | 0.39305 | | | | | | 0.85409 | | | | | | 0.02087 | | |
| Childhood | GSDMA | 17q21.1 | 17q12 | C | 2.2E-37 | | 5.9E-63 | 4.2E-55 | 4.5E-23 | 0.00069 | -12.776 | | -16.747 | -15.636 | -9.8931 | 3.39297 | -0.2426 | | -1.1794 | -0.4181 | -0.4365 | 0.12375 | 0.22441 | | 0.17168 | 0.47831 | 0.11681 | 0.15111 | -1.2149 | | -1.3668 | -0.709 | -1.5683 | -1.4356 | -0.0159 | | -0.0674 | -0.0121 | -0.0472 | -0.0372 |
| Childhood | GSDMB | 17q12 | 17q12 | C | 3.1E-93 | 0.32736 | 5.9E-15 | 2E-113 | 8E-104 | 2E-99 | 20.4828 | -0.9794 | 7.80594 | 22.6266 | 21.6384 | 21.159 | 0.76275 | -0.1021 | 0.31933 | 0.63137 | 0.4165 | 0.4463 | 0.29517 | 0.22959 | 0.1102 | 0.37469 | 0.41889 | 0.51745 | 1.04685 | 1.20143 | 1.59731 | 0.88772 | 0.80834 | 0.64729 | 0.02644 | 0.08681 | 0.05025 | 0.01646 | 0.01041 | 0.00905 |
| Childhood | IKZF3 | 17q12 | 17q12 | C | | | | | | 5E-100 | | | | | | -21.232 | | | | | | -2.1919 | | | | | | 0.59442 | | | | | | -0.5324 | | | | | | -0.0364 |
| Childhood | KRT12 | 17q21.2 | 17q12 | C | | | | 3.3E-14 | | | | | | -7.5856 | | | | | | -0.1943 | | | | | | 0.93375 | | | | | | -0.0831 | | | | | | -0.0033 | | |
| Childhood | LINC00672 | 17q12 | 17q12 | C | 1.8E-07 | | 0.00311 | 0.64836 | | 0.12636 | -5.2204 | | 2.95681 | 0.45604 | | -1.5286 | -0.3466 | | 0.11118 | 0.03079 | | -0.0632 | 0.99206 | | 0.57366 | 0.06953 | | 0.62432 | -0.0099 | | -0.5627 | 1.81499 | | 0.48974 | -0.0004 | | -0.0148 | 0.0846 | | 0.01499 |
| Childhood | MED24 | 17q21.1 | 17q12 | C | 8.3E-11 | 1.9E-14 | 4.2E-13 | | 1.3E-38 | 8.6E-25 | -6.4954 | -7.654 | -7.248 | | -12.995 | -10.28 | -0.2458 | -0.2834 | -0.194 | | -0.4079 | -0.7968 | 0.17614 | 0.06839 | 0.07659 | | 0.07477 | 0.10344 | -1.3527 | -1.8224 | -1.7708 | | -1.7819 | -1.6284 | -0.0333 | -0.0458 | -0.0319 | | -0.0353 | -0.086 |
| Childhood | MSL1 | 17q21.1 | 17q12 | C | 6.1E-10 | 2E-17 | 1.7E-16 | | 5.2E-20 | 4.2E-12 | -6.188 | -8.4922 | -8.245 | | -9.1599 | -6.9296 | -0.227 | -0.3632 | -0.2476 | | -0.3042 | -0.3148 | 0.77356 | 0.66028 | 0.74836 | | 0.853 | 0.44005 | -0.2877 | 0.43952 | -0.3208 | | 0.18529 | -0.7721 | -0.0074 | 0.01375 | -0.0046 | | 0.00441 | -0.0263 |
| Childhood | ORMDL3 | 17q12 | 17q12 | C | 5E-101 | 1.2E-24 | 7.4E-58 | | 7E-99 | 1E-109 | 21.3423 | 10.2462 | 16.0343 | | 21.1093 | 22.2441 | 0.93875 | 0.48972 | 1.08485 | | 0.35909 | 0.479 | 0.31519 | 0.43096 | 0.25949 | | 0.57734 | 0.50004 | 1.00439 | 0.78755 | 1.12759 | | 0.55727 | 0.67442 | 0.02953 | 0.02496 | 0.04614 | | 0.0062 | 0.0094 |
| Childhood | PGAP3 | 17q12 | 17q12 | C | 1.2E-53 | 1.6E-39 | 9.1E-41 | 8.8E-52 | | | 15.4183 | 13.1559 | 13.37 | 15.1403 | | | 0.70112 | 0.48048 | 0.43426 | 0.78134 | | | 0.69701 | 0.97681 | 0.46588 | 0.74084 | | | 0.38935 | -0.0291 | 0.7292 | 0.33075 | | | 0.01552 | -0.0044 | 0.01537 | 0.01218 | | |
| Childhood | PNMT | 17q12 | 17q12 | C | 6.6E-30 | | | | | | 11.3599 | | | | | | 0.65293 | | | | | | 0.4901 | | | | | | 0.69014 | | | | | | 0.02752 | | | | | |
| Childhood | PPP1R1B | 17q12 | 17q12 | C | | 7.1E-43 | 9.8E-45 | | | | | -13.726 | -14.033 | | | | | -1.218 | -0.8001 | | | | | 0.25265 | 0.02168 | | | | | -1.1439 | -2.2959 | | | | | -0.0711 | -0.0924 | | | |
| Childhood | RP11-387H17 | 17q12 | 17q12 | C | | 2E-33 | 6.9E-69 | | | | | -12.049 | -17.542 | | | | | -1.3511 | -0.914 | | | | | 0.84605 | 0.89335 | | | | | -0.1942 | 0.13406 | | | | | -0.0149 | 0.00566 | | | |
| Childhood | RP11-690G19 | 17q12 | 17q12 | C | | 4.6E-07 | | 0.00202 | | | | -5.0433 | | -3.0876 | | | | -0.3785 | | -0.0619 | | | | 0.24149 | | 0.05735 | | | | -1.1713 | | 1.90064 | | | | -0.0618 | | 0.02738 | | |
| Childhood | SMARCE1 | 17q21.2 | 17q12 | C | 0.00011 | 7.4E-09 | 0.915 | | 0.00139 | | -3.8683 | -5.7806 | -0.1067 | | -3.1975 | | -0.105 | -0.2262 | -0.0074 | | -0.1444 | | 0.22998 | 0.83625 | 0.46718 | | 0.50437 | | -1.2004 | -0.2067 | -0.7271 | | -0.6676 | | -0.0219 | -0.0057 | -0.0106 | | -0.0217 | |
| Childhood | STARD3 | 17q12 | 17q12 | C | | | | | | 2.4E-11 | | | | | | 6.68198 | | | | | | 0.32104 | | | | | | 0.04066 | | | | | | 2.04696 | | | | | | 0.06761 |
| Childhood | ZPBP2 | 17q12 | 17q12 | C | | | | | 2.9E-61 | | | | | | -16.515 | | | | | | -0.7371 | | | | | | 0.59617 | | | | | | -0.5299 | | | | | | -0.0163 | |
| Childhood | SERPINB7 | 18q21.33 | 18q21.33 | C | | 1.4E-16 | 3.5E-09 | | | | | -8.2621 | -5.9054 | | | | | -0.3299 | -0.3448 | | | | | 0.51863 | 0.59915 | | | | | -0.6455 | 0.52562 | | | | | -0.0175 | 0.02489 | | | |
| Mixed | AFF4 | 5q31.1 | 5q31.1 | S | | 0.00946 | | 4.8E-10 | 0.00018 | 5.4E-06 | | 2.59514 | | 6.22654 | 3.7412 | 4.5476 | | 0.09577 | | 0.21266 | 0.14596 | 0.21764 | | 0.89165 | | 0.0143 | 0.73239 | 0.41935 | | -0.1362 | | 2.44971 | 0.34195 | 0.80755 | | -0.0065 | | 0.05803 | 0.01027 | 0.0273 |
| Mixed | IL4 | 5q31.1 | 5q31.1 | S | 2.5E-09 | 4.3E-05 | 5.7E-12 | | | 6.9E-16 | -5.9593 | -4.0887 | -6.8872 | | | -8.0726 | -0.5181 | -0.1522 | -0.4037 | | | -1.1157 | 0.85453 | 0.13594 | 0.04963 | | | 0.00233 | 0.18334 | -1.4911 | -1.9631 | | | -3.0443 | 0.01245 | -0.0363 | -0.082 | | | -0.3013 |
| Mixed | IRF1 | 5q31.1 | 5q31.1 | S | | | 4.2E-13 | 0.11837 | 5E-11 | | | | -7.2505 | -1.5617 | -6.5707 | | | | -0.976 | -0.0382 | -0.3843 | | | | 5E-13 | 0.84952 | 9E-09 | | | | -7.2247 | -0.1897 | -5.7481 | | | | -0.6544 | -0.009 | -0.2258 | |
| Mixed | KIF3A | 5q31.1 | 5q31.1 | S | 3.3E-12 | | 9.3E-20 | | | | -6.9658 | | -9.0968 | | | | -0.7831 | | -0.5218 | | | | 2.1E-05 | | 0.00114 | | | | -4.2502 | | -3.2524 | | | | -0.3325 | | -0.1279 | | | |
| Mixed | PDLIM4 | 5q31.1 | 5q31.1 | S | | 3.4E-11 | 8.2E-10 | | | 0.08723 | | 6.62672 | 6.14116 | | | 1.71017 | | 0.34198 | 0.23582 | | | 0.18289 | | 1.1E-08 | 2.1E-10 | | | 0.16492 | | 5.7173 | 6.35687 | | | 1.38871 | | 0.19835 | 0.16435 | | | 0.10011 |
| Mixed | RAD50 | 5q31.1 | 5q31.1 | S | 0.01242 | | 6.6E-10 | 0.02752 | | | 2.49993 | | -6.1758 | 2.20413 | | | 0.08099 | | -0.2549 | 0.03086 | | | 0.00083 | | 0.0005 | 0.43951 | | | 3.34358 | | -3.4782 | 0.77303 | | | 0.07316 | | -0.1111 | 0.00609 | | |
| Mixed | SEPT8 | 5q31.1 | 5q31.1 | S | | 4.7E-14 | 1.6E-13 | | 0.7728 | | | 7.54131 | 7.38131 | | -0.2887 | | | 0.20924 | 0.16412 | | -0.0448 | | | 3.7E-05 | 2.1E-05 | | 0.41714 | | | 4.12287 | 4.25549 | | -0.8114 | | | 0.08028 | 0.06597 | | -0.0607 | |
| Mixed | SLC22A5 | 5q31.1 | 5q31.1 | S | 2.9E-12 | 1.5E-10 | 5.2E-10 | 1.3E-08 | 0.00066 | 2.3E-14 | 6.98165 | 6.40887 | 6.21261 | 5.68085 | 3.4039 | 7.6343 | 0.24423 | 0.15399 | 0.14097 | 0.19251 | 0.04631 | 0.20283 | 3.1E-11 | 1E-11 | 4.4E-11 | 0.00018 | 3.8E-07 | 1.6E-11 | 6.64398 | 6.80448 | 6.59077 | 3.73896 | 5.08106 | 6.73469 | 0.15632 | 0.11014 | 0.10057 | 0.08612 | 0.05425 | 0.11958 |
| Shared | AC007248.6 | 2q12.1 | 2q12.1 | S | | | | | | 2.9E-46 | | | | | | -14.28 | | | | | | -2.0008 | | | | | | 1.8E-10 | | | | | | -6.3743 | | | | | | -0.551 |
| Shared | IL18R1 | 2q12.1 | 2q12.1 | S | 3.3E-08 | 3.7E-36 | 8.3E-29 | | | 1.6E-08 | 5.52493 | 12.555 | 11.1371 | | | 5.65211 | 0.13628 | 0.64634 | 0.77756 | | | 0.30695 | 0.54928 | 4.7E-09 | 5.7E-06 | | | 0.7316 | -0.5988 | 5.85605 | 4.53566 | | | -0.343 | -0.0093 | 0.19933 | 0.21 | | | -0.0115 |
| Shared | IL18RAP | 2q12.1 | 2q12.1 | S | 4.3E-06 | 2.5E-19 | 3.8E-27 | | | 5.6E-14 | -4.5945 | 8.98982 | 10.7914 | | | -7.5181 | -0.2799 | 0.3913 | 0.48368 | | | -0.1532 | 0.07468 | 1.4E-12 | 1E-08 | | | 0.39856 | 1.78243 | 7.08699 | 5.7283 | | | -0.8442 | 0.07386 | 0.19586 | 0.17037 | | | -0.01 |
| Shared | IL1RL2 | 2q12.1 | 2q12.1 | S | | 1.7E-13 | 4.9E-10 | | | | | 7.37192 | 6.22134 | | | | | 0.22762 | 0.21461 | | | | | 1.7E-05 | 0.00013 | | | | | 4.29685 | 3.82045 | | | | | 0.08838 | 0.08964 | | | |
| Shared | AC093642.3 | 2q37.3 | 2q37.3 | S | 0.08872 | 9.4E-08 | 1.1E-06 | 5.6E-05 | 0.34405 | | -1.7022 | -5.3388 | -4.8685 | -4.0299 | 0.94619 | | -0.1077 | -0.2245 | -0.3548 | -0.2441 | 0.01767 | | 0.73776 | 0.00231 | 0.05257 | 0.01092 | 0.15158 | | -0.3348 | -3.047 | -1.9384 | -2.5454 | -1.434 | | -0.0129 | -0.0864 | -0.0971 | -0.104 | -0.0149 | |
| Shared | D2HGDH | 2q37.3 | 2q37.3 | S | 2.5E-11 | 8.9E-10 | 1.4E-10 | 9.6E-05 | 8.1E-13 | 7.1E-07 | 6.67617 | 6.12796 | 6.41218 | 3.90015 | 7.15879 | 4.95794 | 0.13622 | 0.09913 | 0.11573 | 0.06488 | 0.15448 | 0.12921 | 2.6E-10 | 3.6E-10 | 2.7E-10 | 1.6E-05 | 1.4E-10 | 4E-07 | 6.31916 | 6.26884 | 6.31307 | 4.3182 | 6.41487 | 5.06891 | 0.08735 | 0.06879 | 0.07709 | 0.04881 | 0.09349 | 0.08992 |
| Shared | GAL3ST2 | 2q37.3 | 2q37.3 | S | 0.00021 | 1.1E-13 | 1E-10 | | | | -3.7079 | -7.4277 | -6.4675 | | | | -0.1317 | -0.6808 | -0.4956 | | | | 0.09081 | 9.2E-10 | 3.8E-08 | | | | -1.6912 | -6.1222 | -5.4983 | | | | -0.0381 | -0.3767 | -0.2826 | | | |
| Shared | SLC25A46 | 5q22.1 | 5q22.1 | S | 0.65692 | | 4E-06 | | | 1.4E-07 | -0.4442 | | -4.6093 | | | -5.2697 | -0.0509 | | -0.2127 | | | -0.3848 | 0.10261 | | 0.02359 | | | 0.01843 | 1.63235 | | -2.2637 | | | -2.3569 | 0.10589 | | -0.0733 | | | -0.1201 |
| Shared | TMEM232 | 5q22.1 | 5q22.1 | S | 0.08754 | 5.9E-06 | 5.6E-07 | | 4.3E-12 | | 1.70853 | 4.52969 | 5.00543 | | 6.92816 | | 0.08259 | 0.24499 | 0.38505 | | 0.3026 | | 0.78056 | 0.00309 | 0.00947 | | 0.00078 | | -0.2786 | 2.95858 | 2.59458 | | 3.36137 | | -0.0075 | 0.11388 | 0.13729 | | 0.09977 | |
| Shared | TSLP | 5q22.1 | 5q22.1 | S | | 2.7E-10 | 4E-12 | | 0.02286 | | | 6.31634 | 6.93835 | | 2.27584 | | | 0.16547 | 0.17214 | | 0.0481 | | | 1.4E-05 | 1.4E-05 | | 0.76783 | | | 4.34315 | 4.34489 | | 0.29521 | | | 0.07626 | 0.07252 | | 0.00457 | |
| Shared | HLA-DRA | 6p21.32 | 6p21.33 / 6p21.3 | A | | | 0.24106 | | | | | | 1.17233 | | | | | | 0.09336 | | | | | | 5.5E-11 | | | | | | 6.5571 | | | | | | 0.31521 | | | |
| Shared | PRRT1 | 6p21.32 | 6p21.33 / 6p21.3 | A | | 0.41422 | 0.75586 | | | | | -0.8165 | -0.3109 | | | | | -0.0145 | -0.0043 | | | | | 8.4E-10 | 1.3E-06 | | | | | 6.13639 | 4.84208 | | | | | 0.09124 | 0.08139 | | | |
| Shared | VPS52 | 6p21.32 | 6p21.33 / 6p21.3 | A | 0.07438 | | 0.62176 | | | | -1.7843 | | 0.49336 | | | | -0.1925 | | 0.06681 | | | | 0.70093 | | 3.8E-08 | | | | 0.38407 | | -5.4989 | | | | 0.02864 | | -0.255 | | | |
| Shared | HLA-DPA1 | 6p21.32 | 6p21.33 / 6p21.3 | C | 7.9E-09 | 2.1E-05 | 1.3E-06 | | | | -5.7712 | -4.2528 | -4.8449 | | | | -0.3044 | -0.1592 | -0.1407 | | | | 0.18196 | 0.28423 | 0.57561 | | | | -1.3347 | 1.07086 | 0.55981 | | | | -0.0395 | 0.02767 | 0.01128 | | | |
| Shared | ABHD16A | 6p21.33 | 6p21.33 / 6p21.3 | S | 1.8E-08 | 1.3E-05 | 0.0002 | | | | 5.63403 | 4.366 | 3.71896 | | | | 0.25226 | 0.35732 | 0.28042 | | | | 2.8E-10 | 6.1E-09 | 2.7E-05 | | | | 6.31145 | 5.8145 | 4.19847 | | | | 0.17842 | 0.29417 | 0.19872 | | | |
| Shared | AGER | 6p21.32 | 6p21.33 / 6p21.3 | S | 2.5E-12 | | 0.00989 | | | 0.69909 | -7.0053 | | 2.57951 | | | 0.38655 | -0.2725 | | 0.12698 | | | 0.01212 | 9.9E-06 | | 5E-08 | | | 0.21298 | -4.4186 | | 5.45169 | | | 1.24541 | -0.1157 | | 0.20506 | | | 0.07425 |
| Shared | APOM | 6p21.33 | 6p21.33 / 6p21.3 | S | 0.76974 | | 0.00037 | | | | 0.29271 | | -3.5624 | | | | 0.11585 | | -0.2154 | | | | 0.24534 | | 1.4E-07 | | | | -1.1617 | | -5.2666 | | | | -0.2277 | | -0.2083 | | | |
| Shared | ATF6B | 6p21.33 | 6p21.33 / 6p21.3 | S | 1.6E-09 | 2.3E-12 | 1.2E-06 | | | 2.6E-24 | 6.03628 | 7.01636 | 4.86266 | | | 10.1736 | 0.18957 | 0.22167 | 0.1205 | | | 0.84494 | 1E-11 | 4E-14 | 4.2E-10 | | | 2.7E-20 | 6.80614 | 7.56197 | 6.24792 | | | 9.23036 | 0.14231 | 0.15927 | 0.10751 | | | 0.49825 |
| Shared | BAG6 | 6p21.33 | 6p21.33 / 6p21.3 | S | 0.72086 | | 0.0009 | | | 0.50199 | 0.35731 | | 3.31999 | | | 0.67136 | 0.04133 | | 0.14322 | | | 0.05453 | 0.41343 | | 7E-09 | | | 1.9E-07 | 0.81787 | | 5.79097 | | | 5.21087 | 0.03959 | | 0.16759 | | | 0.21641 |
| Shared | BTNL2 | 6p21.32 | 6p21.33 / 6p21.3 | S | | | | 1.9E-06 | | | | | | 4.76518 | | | | | | 0.08572 | | | | | | 7.7E-10 | | | | | | 6.15167 | | | | | | 0.07155 | | |
| Shared | C4A | 6p21.33 | 6p21.33 / 6p21.3 | S | 0.00432 | 0.00048 | 4.7E-06 | 1.6E-06 | 0.01668 | 1.1E-07 | -2.8539 | -3.4942 | -4.5772 | -4.7997 | -2.3937 | -5.3081 | -0.0511 | -0.0622 | -0.0869 | -0.112 | -0.0425 | -0.1362 | 4.5E-06 | 2.3E-07 | 6.9E-11 | 2.9E-05 | 0.09489 | 7.3E-10 | -4.5868 | -5.1739 | -6.5236 | -4.1803 | -1.6701 | -6.1587 | -0.0561 | -0.0666 | -0.0864 | -0.0655 | -0.0195 | -0.1119 |
| Shared | C4B | 6p21.33 | 6p21.33 / 6p21.3 | S | 0.99253 | 0.00026 | 4E-06 | | | 0.00291 | 0.00937 | 3.64918 | 4.6106 | | | 2.97659 | 0.05739 | 0.08913 | 0.10412 | | | 0.07037 | 0.14921 | 1.2E-05 | 2.3E-07 | | | 3.7E-08 | -1.4423 | 4.37645 | 5.1703 | | | 5.50494 | -0.0345 | 0.07206 | 0.08008 | | | 0.09007 |
| Shared | C6orf48 | 6p21.33 | 6p21.33 / 6p21.3 | S | 0.00045 | 2.2E-05 | 0.02681 | 0.13031 | 0.09294 | 2.3E-06 | 3.50607 | 4.24853 | 2.21425 | -1.5129 | 1.6801 | 4.72361 | 0.17719 | 0.34736 | 0.20825 | -0.0413 | 0.04905 | 0.61045 | 0.0838 | 0.12315 | 0.29324 | 5.2E-07 | 0.00396 | 0.01945 | 1.72905 | 1.54167 | 1.05103 | -5.02 | 2.88143 | 2.33686 | 0.05417 | 0.08632 | 0.07032 | -0.1241 | 0.0555 | 0.19833 |
| Shared | CLIC1 | 6p21.33 | 6p21.33 / 6p21.3 | S | | | | | 3.3E-06 | 8.7E-06 | | | | | -4.6486 | -4.446 | | | | | -0.2361 | -0.4071 | | | | | 7.1E-05 | 2.4E-07 | | | | | -3.9718 | -5.1665 | | | | | -0.1404 | -0.3229 |
| Shared | CYP21A2 | 6p21.33 | 6p21.33 / 6p21.3 | S | 0.07392 | 0.00033 | 0.00088 | 0.7015 | | 0.00164 | 1.78709 | 3.58982 | 3.32524 | 0.38329 | | 3.14796 | 0.03801 | 0.10061 | 0.06827 | 0.00832 | | 0.10998 | 1E-06 | 4.8E-09 | 0.00093 | 2.9E-08 | | 0.0001 | 4.89016 | 5.85345 | 3.31121 | 5.55022 | | 3.88738 | 0.12605 | 0.12097 | 0.05412 | 0.07841 | | 0.11417 |
| Shared | DHFRP2 | 6p21.33 | 6p21.33 / 6p21.3 | S | | | | | | 1.3E-06 | | | | | | -4.8421 | | | | | | -0.566 | | | | | | 1.4E-05 | | | | | | -4.3365 | | | | | | -0.3442 |
| Shared | FKBPL | 6p21.33 | 6p21.33 / 6p21.3 | S | | 0.00326 | 5.9E-05 | | | 1.4E-15 | | 2.94234 | 4.01602 | | | -7.9899 | | 0.35315 | 0.30861 | | | -0.341 | | 0.10752 | 0.09543 | | | 1E-16 | | 1.60946 | 1.66744 | | | -8.2998 | | 0.12802 | 0.08447 | | | -0.2381 |
| Shared | FLOT1 | 6p21.33 | 6p21.33 / 6p21.3 | S | 1.1E-07 | | 0.04893 | 0.3358 | | 2.3E-05 | 5.30555 | | 1.96922 | 0.96249 | | 4.23561 | 0.18308 | | 0.13607 | 0.0198 | | 0.17714 | 8.5E-07 | | 0.51388 | 0.59415 | | 2.1E-05 | 4.9236 | | 0.65281 | 0.53284 | | 4.25484 | 0.11291 | | 0.02098 | -0.0031 | | 0.1175 |
| Shared | GPANK1 | 6p21.33 | 6p21.33 / 6p21.3 | S | 0.0146 | 0.02311 | | | | | -2.4422 | -2.2717 | | | | | -0.1655 | -0.0777 | | | | | 5.8E-11 | 0.00202 | | | | | -6.5495 | -3.0868 | | | | | -0.3183 | -0.0726 | | | | |
| Shared | HLA-C | 6p21.33 | 6p21.33 / 6p21.3 | S | 1.7E-06 | 0.00136 | 0.00015 | 3.2E-05 | 7.4E-06 | 1.3E-07 | -4.7835 | -3.2025 | -3.7939 | -4.1616 | -4.4809 | -5.2817 | -0.1208 | -0.0763 | -0.0791 | -0.1591 | -0.0711 | -0.1268 | 2E-06 | 0.00292 | 0.00316 | 0.00158 | 0.00034 | 0.00013 | -4.7517 | -2.9757 | -2.9513 | -3.1601 | -3.5839 | -3.8183 | -0.0809 | -0.048 | -0.0412 | -0.0785 | -0.0384 | -0.0604 |
| Shared | HLA-DOB | 6p21.32 | 6p21.33 / 6p21.3 | S | 7.8E-09 | 1.4E-05 | 7E-07 | 1.2E-16 | 0.27329 | 0.77996 | 5.77328 | 4.34776 | 4.96173 | 8.28488 | 1.09551 | 0.27937 | 0.11239 | 0.07503 | 0.08945 | 0.38569 | 0.01542 | 0.00036 | 4.2E-09 | 4.9E-09 | 1E-10 | 0.0016 | 0.0015 | 0.00262 | 5.87622 | 5.84973 | 6.45976 | 3.15607 | 3.17509 | 3.00921 | 0.07631 | 0.07033 | 0.07724 | 0.09604 | 0.03644 | 0.03864 |
| Shared | HLA-DQA1 | 6p21.32 | 6p21.33 / 6p21.3 | S | 4.1E-37 | 4E-20 | 2.1E-20 | 5.5E-23 | 2E-23 | 1.3E-36 | -12.728 | -9.187 | -9.2594 | -9.8723 | -9.9736 | -12.635 | -0.3218 | -0.1734 | -0.1748 | -0.4431 | -0.2321 | -0.4692 | 1.4E-47 | 1.9E-47 | 4E-44 | 4.2E-47 | 1.9E-47 | 4.5E-44 | -14.492 | -14.469 | -13.934 | -14.414 | -14.468 | -13.924 | -0.2452 | -0.1792 | -0.1712 | -0.4308 | -0.2199 | -0.3433 |
| Shared | HLA-DQA2 | 6p21.32 | 6p21.33 / 6p21.3 | S | 3.9E-26 | 1E-24 | 2.5E-25 | 3.6E-29 | 1.4E-20 | 8.4E-21 | 10.5752 | 10.2645 | 10.3992 | 11.2106 | 9.29823 | 9.35403 | 0.14865 | 0.1701 | 0.15737 | 0.15134 | 0.15097 | 0.12916 | 8.5E-61 | 3.9E-54 | 1.7E-46 | 5.8E-56 | 1E-47 | 6.6E-45 | 16.4494 | 15.493 | 14.3182 | 15.761 | 14.5131 | 14.0614 | 0.15474 | 0.17041 | 0.14181 | 0.14098 | 0.16183 | 0.13006 |
| Shared | HLA-DQB1 | 6p21.32 | 6p21.33 / 6p21.3 | S | 7.1E-19 | 1.4E-30 | 3.6E-33 | 1.9E-18 | 2.7E-24 | 1.4E-29 | -8.874 | -11.492 | -12 | -8.7651 | -10.172 | -11.296 | -0.1717 | -0.2688 | -0.2609 | -0.1929 | -0.1878 | -0.307 | 5.2E-37 | 2.5E-42 | 5.2E-41 | 1.2E-39 | 4.2E-36 | 9.1E-41 | -12.71 | -13.634 | -13.411 | -13.177 | -12.546 | -13.37 | -0.1687 | -0.2136 | -0.1951 | -0.1895 | -0.1542 | -0.2448 |
| Shared | HLA-DQB2 | 6p21.32 | 6p21.33 / 6p21.3 | S | 2.3E-46 | 0.04445 | 4.1E-12 | 8.1E-44 | 6.7E-27 | 1.2E-28 | 14.2971 | 2.00982 | 6.93358 | 13.8825 | 10.7383 | 11.1018 | 0.22394 | 0.09094 | 0.3352 | 0.30172 | 0.19033 | 0.19481 | 3.2E-56 | 0.00996 | 5.6E-10 | 1.9E-46 | 3.5E-44 | 7.8E-43 | 15.7976 | 2.57735 | 6.201 | 14.3104 | 13.943 | 13.7191 | 0.16461 | 0.0714 | 0.19702 | 0.20413 | 0.16115 | 0.16078 |
| Shared | HLA-DRB1 | 6p21.32 | 6p21.33 / 6p21.3 | S | 8.2E-16 | 3.8E-15 | 2.5E-11 | 1.2E-11 | 0.00219 | 0.66688 | -8.0513 | -7.8615 | -6.6758 | -6.778 | 3.06268 | -0.4304 | -0.1637 | -0.245 | -0.1862 | -0.3254 | 0.11611 | -0.0212 | 7.3E-37 | 1.5E-39 | 5.5E-28 | 1.9E-29 | 0.09564 | 4.9E-14 | -12.684 | -13.161 | -10.968 | -11.269 | 1.66638 | -7.5333 | -0.1739 | -0.2696 | -0.2054 | -0.3623 | 0.04011 | -0.1786 |
| Shared | HLA-DRB5 | 6p21.32 | 6p21.33 / 6p21.3 | S | 0.08884 | 0.02276 | 0.01725 | 0.79988 | 0.02142 | 0.00545 | 1.70155 | 2.27749 | 2.38129 | 0.2535 | 2.30052 | 2.77899 | 0.0262 | 0.03958 | 0.04207 | 0.00317 | 0.04392 | 0.04145 | 2.6E-06 | 0.00031 | 0.00016 | 1.2E-09 | 1.1E-06 | 9.5E-05 | -4.6986 | -3.6097 | -3.7768 | -6.0759 | -4.88 | -3.9019 | -0.0481 | -0.0377 | -0.0426 | -0.0826 | -0.0608 | -0.0421 |
| Shared | HLA-DRB6 | 6p21.32 | 6p21.33 / 6p21.3 | S | 3.9E-07 | 0.00013 | 4.3E-05 | 0.02759 | 2.9E-05 | 5.6E-06 | 5.07584 | 3.82242 | 4.08919 | 2.20307 | 4.18016 | 4.54132 | 0.06951 | 0.05519 | 0.0588 | 0.03988 | 0.05087 | 0.06825 | 3.2E-33 | 5.1E-30 | 4E-29 | 1.1E-19 | 1.9E-21 | 1.5E-27 | 12.0094 | 11.3835 | 11.2007 | 9.07478 | 9.50987 | 10.8744 | 0.11504 | 0.1126 | 0.10687 | 0.09948 | 0.07991 | 0.1153 |
| Shared | HLA-DRB9 | 6p21.32 | 6p21.33 / 6p21.3 | S | 4.1E-15 | 0.00467 | 0.79295 | 0.04456 | 0.21597 | 2.3E-09 | 7.85315 | -2.8292 | -0.2625 | 2.00879 | 1.23733 | 5.97425 | 0.14058 | -0.0786 | -0.0097 | 0.05096 | 0.01254 | 0.20564 | 6.1E-29 | 2.1E-10 | 3.8E-11 | 1.9E-14 | 1.7E-13 | 1.5E-27 | 11.1639 | 6.35362 | 6.61295 | 7.65786 | 7.37323 | 10.8776 | 0.12898 | 0.10589 | 0.15099 | 0.11802 | 0.05935 | 0.2582 |
| Shared | HSPA1L | 6p21.33 | 6p21.33 / 6p21.3 | S | | | | | | 5.5E-15 | | | | | | -7.8151 | | | | | | -1.3924 | | | | | | 1E-08 | | | | | | -5.7303 | | | | | | -0.6664 |
| Shared | LY6G5B | 6p21.33 | 6p21.33 / 6p21.3 | S | 0.00118 | 4.8E-05 | 2.4E-05 | | 1.8E-07 | 8.6E-05 | 3.24406 | 4.067 | 4.22503 | | 5.21884 | 3.92625 | 0.07969 | 0.17511 | 0.12276 | | 0.45443 | 0.20099 | 0.92947 | 0.01789 | 0.05264 | | 0.00054 | 0.88877 | -0.0885 | 2.36791 | 1.93784 | | 3.45786 | -0.1399 | -0.0006 | 0.0697 | 0.03867 | | 0.19977 | -0.0002 |
| Shared | MDC1 | 6p21.33 | 6p21.33 / 6p21.3 | S | 1.2E-06 | | | | 0.70675 | | -4.8552 | | | | -0.3762 | | -0.2448 | | | | -0.0113 | | 0.00453 | | | | 0.00088 | | -2.839 | | | | -3.326 | | -0.0954 | | | | -0.0753 | |
| Shared | MICA | 6p21.33 | 6p21.33 / 6p21.3 | S | 2.6E-19 | 2.3E-09 | 2.3E-18 | 1.7E-08 | 8.8E-16 | 1.5E-14 | 8.98504 | 5.97647 | 8.74119 | 5.63859 | 8.04219 | 7.68534 | 0.18798 | 0.13509 | 0.17963 | 0.09412 | 0.15012 | 0.19794 | 1.3E-17 | 1.4E-15 | 2.2E-18 | 1.9E-08 | 1.7E-16 | 1.6E-22 | 8.54727 | 7.98628 | 8.74725 | 5.62078 | 8.24341 | 9.76557 | 0.11617 | 0.1211 | 0.11416 | 0.06028 | 0.10103 | 0.166 |
| Shared | MICB | 6p21.33 | 6p21.33 / 6p21.3 | S | 9.2E-06 | 1.7E-05 | 2E-05 | | 0.00272 | 0.00118 | 4.43495 | 4.30053 | 4.26189 | | 2.99805 | 3.24265 | 0.08952 | 0.06252 | 0.07673 | | 0.05971 | 0.07059 | 7.8E-12 | 3.3E-05 | 1.9E-07 | | 5.9E-08 | 9E-08 | 6.84157 | 4.14897 | 5.20757 | | 5.42227 | 5.34533 | 0.09322 | 0.03888 | 0.06068 | | 0.07514 | 0.07802 |
| Shared | NFKBIL1 | 6p21.33 | 6p21.33 / 6p21.3 | S | | | | | 0.0022 | | | | | | 3.06145 | | | | | | 0.08786 | | | | | | 6.1E-07 | | | | | | 4.98877 | | | | | | 0.09252 | |
| Shared | NOTCH4 | 6p21.32 | 6p21.33 / 6p21.3 | S | 0.12084 | 0.76981 | 0.14301 | 0.31658 | 0.23671 | 3.2E-08 | -1.5513 | 0.29262 | -1.4647 | -1.0015 | 1.18326 | -5.5297 | -0.0347 | 0.0079 | -0.0317 | -0.0475 | 0.07476 | -0.1163 | 3E-06 | 0.00139 | 7.7E-05 | 0.00192 | 0.05912 | 0.00105 | -4.6686 | -3.196 | -3.9544 | -3.1026 | -1.8873 | -3.276 | -0.0719 | -0.0531 | -0.0602 | -0.099 | -0.0827 | -0.0443 |
| Shared | PPP1R18 | 6p21.33 | 6p21.33 / 6p21.3 | S | 5.3E-13 | | | | | | -7.2166 | | | | | | -0.3431 | | | | | | 2.1E-07 | | | | | | -5.1912 | | | | | | -0.1654 | | | | | |
| Shared | PSMB9 | 6p21.32 | 6p21.33 / 6p21.3 | S | 2.9E-07 | 4E-10 | 5.1E-12 | 1.1E-05 | 0.56222 | 5.2E-09 | 5.12689 | 6.25366 | 6.90288 | 4.38926 | 0.57955 | 5.84166 | 0.1211 | 0.15618 | 0.22852 | 0.13556 | 0.01651 | 0.11867 | 0.00255 | 8.6E-10 | 4.1E-07 | 0.0022 | 0.26007 | 6.4E-06 | 3.01675 | 6.13258 | 5.06338 | 3.06198 | -1.1262 | 4.51286 | 0.04866 | 0.10084 | 0.11267 | 0.06335 | -0.037 | 0.06148 |
| Shared | SAPCD1 | 6p21.33 | 6p21.33 / 6p21.3 | S | | | 1.9E-05 | | | | | | -4.2781 | | | | | | -0.1976 | | | | | | 5.2E-13 | | | | | | -7.2191 | | | | | | -0.2276 | | | |
| Shared | STK19P | 6p21.33 | 6p21.33 / 6p21.3 | C | 9E-06 | 0.01841 | 0.00187 | 1.7E-11 | | 0.02112 | -4.4398 | -2.3572 | -3.1094 | -6.7301 | | -2.3059 | -0.1318 | -0.0605 | -0.0639 | -0.2687 | | -0.0722 | 0.07623 | 0.99803 | 0.74426 | 0.04829 | | 0.69496 | -1.773 | -0.0025 | -0.3262 | -1.9748 | | -0.3921 | -0.0322 | 0.00234 | -0.0048 | -0.0541 | | -0.0101 |
| Shared | TAP2 | 6p21.32 | 6p21.33 / 6p21.3 | S | 0.58528 | 0.42317 | 0.2621 | | 7.4E-05 | 1.8E-10 | 0.54568 | 0.80093 | 1.12143 | | -3.9636 | -6.3811 | 0.00414 | -0.0033 | -0.0062 | | -0.098 | -0.1759 | 0.00638 | 0.00031 | 0.13327 | | 1.6E-11 | 7.6E-14 | -2.7276 | 3.60596 | 1.50132 | | -6.7402 | -7.4774 | -0.0751 | 0.07797 | 0.01344 | | -0.1158 | -0.1391 |
| Shared | TCF19 | 6p21.33 | 6p21.33 / 6p21.3 | S | 0.00131 | 0.00442 | 4.1E-10 | | 1.5E-05 | 2.6E-07 | -3.2131 | 2.84649 | 6.24942 | | -4.3328 | -5.1503 | -0.1818 | 0.12499 | 0.17205 | | -0.142 | -0.2994 | 0.50938 | 0.00501 | 3.1E-06 | | 0.00038 | 0.00016 | -0.6598 | 2.8066 | 4.66447 | | -3.5527 | -3.779 | -0.0415 | 0.08059 | 0.08829 | | -0.0768 | -0.1365 |
| Shared | TNXA | 6p21.33 | 6p21.33 / 6p21.3 | S | 2.8E-25 | 6.8E-15 | 4.1E-05 | 1.1E-09 | | | -10.387 | -7.7882 | -4.1036 | -6.0923 | | | -0.2189 | -0.2544 | -0.2121 | -0.1926 | | | 1.2E-15 | 4.4E-12 | 7.7E-06 | 2.4E-05 | | | -8.0069 | -6.9221 | -4.4725 | -4.2213 | | | -0.1151 | -0.1474 | -0.1485 | -0.0893 | | |
| Shared | ZBTB12 | 6p21.33 | 6p21.33 / 6p21.3 | S | | | | | | 1E-05 | | | | | | -4.4143 | | | | | | -0.1108 | | | | | | 5E-07 | | | | | | -5.026 | | | | | | -0.0833 |
| Shared | OR2H2 | 6p22.1 | 6p22.1 | S | | 2.2E-05 | 0.05362 | | | 0.02763 | | 4.24301 | 1.92992 | | | 2.20248 | | 0.37514 | 0.09784 | | | 0.1379 | | 7.6E-08 | 0.02638 | | | 2.6E-08 | | 5.37658 | 2.22062 | | | 5.568 | | 0.31756 | 0.07553 | | | 0.22307 |
| Shared | KIAA1432 | 9p24.1 | 9p24.1 | S | | 4.2E-05 | 0.00274 | | | | | 4.09375 | 2.99497 | | | | | 0.1051 | 0.08903 | | | | | 5.3E-08 | 0.00021 | | | | | 5.44053 | 3.7033 | | | | | 0.09755 | 0.07552 | | | |
| Shared | RP11-672A2.[illegible] | 11q13.5 | 11q13.5 | C | 7.8E-07 | | | | 0.02406 | 0.00607 | -4.9394 | | | | -2.2562 | 2.74396 | -0.291 | | | | -0.1669 | 0.1051 | 0.49133 | | | | 0.9502 | 0.75949 | -0.6882 | | | | 0.06246 | 0.30615 | -0.0251 | | | | 0.00028 | 0.00521 |
| Shared | RP11-619A14 | 11q13.5 | 11q13.5 | S | | | | 2E-05 | | 4.4E-08 | | | | 4.26918 | | 5.47521 | | | | 0.12679 | | 0.47501 | | | | 0.05051 | | 0.01551 | | | | 1.95565 | | 2.4203 | | | | 0.03829 | | 0.14208 |
| Shared | RPS26 | 12q13.2 | 12q13.2 | S | 1.1E-10 | 3E-11 | 3.8E-10 | 3E-11 | 2E-10 | 3.8E-11 | 6.45377 | 6.64513 | 6.26127 | 6.64698 | 6.36391 | 6.61007 | 0.11263 | 0.09396 | 0.08913 | 0.15217 | 0.12777 | 0.12458 | 1.5E-07 | 1.5E-07 | 2.2E-07 | 1.5E-07 | 3.7E-07 | 1.2E-07 | 5.24847 | 5.25418 | 5.17902 | 5.25358 | 5.08489 | 5.28827 | 0.06191 | 0.05019 | 0.05011 | 0.08118 | 0.06944 | 0.06707 |
| Shared | SDR9C7 | 12q13.3 | 12q13.2 | S | | 1.8E-10 | 6.6E-08 | | | | | -6.3795 | -5.4008 | | | | | -0.3771 | -0.4533 | | | | | 2E-05 | 0.00016 | | | | | -4.2625 | -3.7712 | | | | | -0.163 | -0.2045 | | | |
| Shared | SUOX | 12q13.2 | 12q13.2 | S | 4.7E-10 | 5.4E-11 | 6E-12 | 3.2E-11 | 6E-10 | 6.9E-11 | -6.2299 | -6.5582 | -6.8805 | -6.6374 | -6.1914 | -6.5219 | -0.1479 | -0.2152 | -0.148 | -0.2413 | -0.1532 | -0.2503 | 5E-07 | 3.2E-07 | 1.9E-07 | 5.5E-07 | 7.3E-06 | 5.8E-07 | -5.025 | -5.1101 | -5.2085 | -5.0073 | -4.4843 | -4.9984 | -0.0813 | -0.1136 | -0.0757 | -0.123 | -0.0748 | -0.1294 |
| Shared | LRP1 | 12q13.3 | 12q13.3 | S | | | 3.9E-09 | | | | | | -5.8866 | | | | | | -0.3275 | | | | | | 0.00129 | | | | | | -3.2186 | | | | | | -0.1222 | | | |
| Shared | SMAD3 | 15q22.33 | 15q22.33 | S | 0.00116 | 0.03641 | 8.2E-08 | | | 6.2E-07 | -3.2493 | -2.0923 | -5.3637 | | | -4.984 | -0.2042 | -0.1294 | -0.3964 | | | -0.2148 | 0.11728 | 0.06965 | 2.1E-07 | | | 0.00026 | -1.5663 | -1.8142 | -5.188 | | | -3.6492 | -0.0667 | -0.0784 | -0.2593 | | | -0.1095 |
| Shared | IL4R | 16p12.1 | 16p12.1 | S | 0.00177 | | | | | 2.1E-09 | -3.126 | | | | | -5.988 | -0.1135 | | | | | -0.2999 | 0.03609 | | | | | 0.01376 | -2.0959 | | | | | -2.4634 | -0.0503 | | | | | -0.0837 |
| Shared | NSMCE1 | 16p12.1 | 16p12.1 | S | | | 1.3E-07 | | | | | | 5.27225 | | | | | | 0.57745 | | | | | | 0.00216 | | | | | | 3.06716 | | | | | | 0.22662 | | | |
| Shared | CLEC16A | 16p13.13 | 16p13.13 | S | 3E-09 | | | | | | 5.93035 | | | | | | 0.52066 | | | | | | 5E-05 | | | | | | 4.05387 | | | | | | 0.23652 | | | | | |

**Table 9. Loci that were genome-wide significant in three previous large GWASs[6,8,9] that were not genome-wide significant in the childhood onset or adult onset GWASs.** For regions in which multiple SNPs were reported in previous GWAS, the SNP with the smallest p-value is shown.

| | | | UK Biobank | | Zhu et al[7] | Pickrell et al[6] | Demenais et al[5] |
|---|---|---|---|---|---|---|---|
| **Locus** | **Position** | **rsID** | **Childhood Onset Asthma *P*-value** | **Adult Onset Asthma *P*-value** | ***P*-value** | ***P*-value** | ***P*-value** |
| 1p36.22 | 1:10557251 | rs662064 | 3·538e-5 | 9·393e-3 | n.r. | 3·2e-8 | n.r. |
| 1p36.11 | 1:25299426 | rs6600246 | 1·105e-4 | 8·870e-3 | 5·04e-9 | n.r. | n.r. |
| 1q23.3 | 1:161159147 | rs4233366 | 1·717e-5 | 2·959e-1 | n.r. | 4·8e-15 | n.r. |
| 1q24.2 | 1:167433420 | rs1723018 | 8·302e-6 | 7·141e-6 | n.r. | 1·4e-8 | n.r. |
| | 1:167420425 | rs2056626 | 1·048e-4 | 1·547e-7 | 8·80e-9 | n.r. | n.r. |
| 3p24.3 | 3:23653570 | rs115913567 | 1·846e-1 | 3·748e-1 | 1·89e-8 | n.r. | n.r. |
| 5q31.3 | 5:141492419 | rs7705042 | 3·814e-7 | 6·530e-4 | n.r. | 2·5e-8 [a] | 7·9e-9 [a] |
| 6p22.1 | 6:28712247 | rs1233578 | 5·013e-5 | 3·224e-5 | n.r. | n.r. | 5·9e-7 |
| 7p21.1 | 7:20560996 | rs6461503 | 9·210e-8 | 1·570e-1 | 2·13e-9 | n.r. | n.r |
| 7q22.3 | 7:105676505 | rs6959584 | 4·581e-4 | 6·970e-1 | n.r. | 2·0e-8 | n.r. |
| 20q13.33 | 20:62333022 | rs1623866 | 8·822e-6 | 4·930e-2 | 4·04e-9 | n.r. | n.r. |

n.r., not reported as genome-wide significant. [a]The lead SNP reported in this region (rs200634877 at 5:141529512) in Zhu *et al*. is not present in the 1000 genomes database or the imputed UK Biobank data; instead we show the nearest lead SNP at 5q31·3 in Pickrell *et al.* and Demenais *et al*.
